# Prediction of Hemolytic Peptides and their Hemolytic Concentration (HC_50_)

**DOI:** 10.1101/2024.07.23.604887

**Authors:** Anand Singh Rathore, Nishant Kumar, Shubham Choudhury, Naman Kumar Mehta, Gajendra P. S. Raghava

## Abstract

Several peptide-based drugs fail in clinical trials due to their toxicity or hemolytic activity against red blood cells (RBCs). Existing methods predict hemolytic peptides but not the concentration (HC50) required to lyse 50% of RBCs. In this study, we developed a classification model and regression model to identify and quantify the hemolytic activity of peptides. Our models were trained and validated on 1924 peptides with experimentally determined HC50 against mammalian RBCs. Analysis indicates that hydrophobic and positively charged residues were associated with higher hemolytic activity. Our classification models achieved a maximum AUC of 0.909 using a hybrid model of ESM-2 and a motif-based approach. Regression models using compositional features achieved R of 0.739 with R² of 0.543. Our models outperform existing methods and are implemented in the web-based platform HemoPI2 and standalone software for designing hemolytic peptides with desired HC50 values (http://webs.iiitd.edu.in/raghava/hemopi2/).

**Highlights:**

- Developed classification and regression models to predict hemolytic activity and HC50 values of peptides.
- A hybrid model combining machine learning and motif prediction excels in accuracy.
- Benchmarking of the existing classification methods on independent datasets.
- Web server, standalone software, and pip package for hemolytic activity prediction of peptides/proteins.

## 1 Introduction

The process of developing and testing drugs is intricate, expensive, time-intensive, and laden with risks. Drug development encompasses several stages, which can be categorized into various phases, including disease-related genomics, identification and validation of targets, lead discovery and optimization, preclinical testing, and clinical trials^1^. In the past few decades, peptide-based drugs have boomed in drug discovery and development because of their advantages over traditional drugs, including greater efficacy, specificity, high tissue penetration ability, low immunogenicity, ease of modification, reduced risk of drug-drug interactions as their degradation product is amino acids, and low cost^2–5^. This trend is evidenced by the approval of 31 peptides as drugs by the Food and Drug Administration (FDA) between 2016 and 2023, alongside 370 new drugs approved during the same period, accounting for more than 8% of the total^6–8^. Additionally, there are over 200 peptides currently in clinical development and approximately 600 peptides undergoing preclinical studies, further highlighting the growing importance and utilization of peptides in pharmaceutical research and development^9,10^. The primary causes of failure of peptide-based drugs during preclinical trials are due to unacceptable safety and efficacy, which is mainly caused by absorption, distribution, metabolism, excretion, and toxicity (ADMET) profiles^11^. Thus, there is a need to effectively screen and enhance the ADMET properties of drugs at an early stage. Many pharmaceutical companies have adopted a “fail early, fail cheap” strategy to address these challenges^12^. Adopting an in-silico strategy for predicting ADMET properties has gained considerable traction due to its cost-saving advantages and ability to provide high-throughput alternatives to traditional experimental measurement methods^13^.

Toxicity, a major obstacle in designing peptide-based therapeutics, encompasses three primary categories: cytotoxicity, hemotoxicity responsible for lysing RBCs, and immunotoxicity allergenicity^14–21^. Hemolytic concentration (HC_50_) serves as a common indicator of peptide toxicity, representing the concentration at which 50% lysis of normal human erythrocytes occurs under physiological conditions^22^. Peptides rich in positively charged amino acids can bind to the erythrocyte’s negatively charged lipid bilayer, leading to membrane disintegration and allowing water and solute molecules to enter the cell. This influx increases the osmotic gradient inside the erythrocyte, resulting in cell swelling and ultimately bursting^23^. Several computational methods have been developed for predicting hemolytic peptides in recent years^24–26^. Most of these methods were trained and tested on datasets derived from a dedicated database of hemolytic peptides referred as Hemolytik^17^. In 2016, Chaudhary et al. attempted to develop a machine learning (ML)-based model and motif-based tool for predicting hemolytic peptides called HemoPI^24^. A method called HemoPred^25^, based on the RF model, was developed utilizing the identical dataset employed in HemoPI. It incorporates a linear combination of amino acid composition (AAC) and dipeptide composition (DPC) as features. HLPpred-Fuse^27^ proposed a two-layer prediction framework. They share the same positive dataset developed by Chaudhry et al., while the negative dataset is sourced from the PEPred-SUITE method^28^. By using the HemoPI dataset, Plisson et al.^29^ developed another ML-based method that predicts the hemolytic nature of peptide sequences using gradient-boosting classifiers. Their multivariate outlier detection models led to the discovery of high-confidence non-hemolytic natural antimicrobial peptides (AMPs), facilitated the de novo design of non-hemolytic peptides, and provided guidelines for designing non-hemolytic peptides. HemoPImod^18^ is a prediction model for chemically modified hemolytic peptides. It utilizes a Random Forest (RF) model that integrates various peptide features, including atom and diatom compositions, 2D and 3D descriptors, as well as fingerprints. HAPPENN^26^ and HemoNet^30^ are two neural network-based classifier models using the DBAASP and Hemolytik databases for dataset construction. HAPPENN represents a state-of-the-art model for predicting hemolytic activity, leveraging features selected via an ensemble of RF model and support vector machines (SVMs). In the HemoNet tool, SMILES-based fingerprints are used as a feature so that it can capture N/C terminal modification. Plisson et al.^29^ explored 14 binary classifiers to predict hemolytic activity across three datasets developed by Chaudhary et al. They utilized 56 sequence-based physicochemical descriptors and employed a ML model called XGBC (Extreme Gradient Boosting Classifier) to discover and design non-hemolytic peptides. In 2021, Capecchi et al.^31^ introduced a recurrent neural network classifier to identify membrane-disruptive amphiphilic antimicrobial peptides. Additionally, they developed a model for designing short non-hemolytic antimicrobial peptides, leveraging data sourced from DBAASP. Ansari et al.^32^ developed a recurrent neural network (Bi-LSTM) with concatenated amino acid frequencies to predict whether a peptide is hemolytic. They utilized the DBAASPv3 database to create their dataset, comprising 9316 positive sequences (length range: 1-77) and negative sequences (length range: 1-190) in the training data. Salem et al.^33^ utilized transfer learning to address the issue of small data and employed a protein language model (PLMs) based on LLM (Large Language Model), employing a tool named AMPDeep. Perveen et al.^34^ introduced an ML approach named Hemolytic-pred, designed for predicting hemolytic proteins. The dataset used in this method was collected from UniProtKB-SwissProt^35^ [18287689], and it employs position and composition-based features. Mendieta et al.^36^ utilize network science and data mining to analyze hemolytic peptides, creating scaffolds to represent their chemical space and uncovering putative hemolytic motifs. Zhuang et al.^37^ developed a tool employing the quantum support vector machine (QSVM) to classify peptides as hemolytic or non-hemolytic, utilizing the dataset from the HemoPI tool. A recently developed PeptideBERT^38^ based approach for hemolytic peptide identification utilizes the ProtBERT pre-trained transformer model featuring 12 attention heads and 12 hidden layers. Several other computational tools have been developed for different types of toxicity. In Table 1, we present a comprehensive list of toxicity prediction methods. While the methods mentioned above have contributed to advancing the discovery of potential hemolytic peptides, further improvement remains necessary. This is primarily because the datasets utilized in these methods are overly generalized to all vertebrates. Moreover, none of these methods have the capability to predict the HC_50_ value of peptides. It is important for drug development to identify the peptide concentration at which any peptide exhibits significant hemolytic activity.

**Table 1:** Computational tools for predicting hemolytic activity of peptides and proteins.

| Tool (year) | Description (link) | Reference |
| --- | --- | --- |
| HemoPI (2016) | Website and mobile application designed for computing the hemolytic activity of peptides.<br>(https://webs.iitd.edu.in/raghava/hemopi/) | <sup>24</sup> |
| HemoPred (2017) | RF-based Web server dedicated to predicting the hemolytic potency of peptides.<br>(http://codes.bio/hemopred/) | <sup>25</sup> |
| HemoPImod (2020) | A web-based prediction model for chemically modified hemolytic peptides.<br>(https://webs.iiitd.edu.in/raghava/hemopimod/) | <sup>18</sup> |
| HLPPred-Fuse (2020) | Enhanced and resilient prediction of hemolytic peptide activity through the fusion of multiple feature representations. (http://thegleelab.org/HLPPred-Fuse) | <sup>27</sup> |
| HAPPENN (2020) | An innovative tool for predicting hemolytic potency in therapeutic peptides utilizing neural networks.<br>(https://research.timmons.eu/happenn) | <sup>26</sup> |
| Plisson et al. (2020) | ML models alongside outlier detection techniques boosts the accuracy in predicting antimicrobial peptides (AMPs) that exhibit diminished hemolytic activity.<br>(https://github.com/plissonf/ML-guided-discovery-and-design-of-non-hemolytic-peptides) | <sup>29</sup> |
| Capecchi et al. (2021) | RNN model trained on antimicrobial peptide data from DBAASP is utilized to design short, non-hemolytic peptides effective against multidrug-resistant bacteria, offering promising alternatives for drug design.<br>(https://github.com/reymond-group/MLpeptide) | <sup>31</sup> |
| HemoNet (2021) | Predicting hemolytic activity of peptides through integrated feature learning techniques.<br>(https://github.com/adibayaseen/HemoNet/) | <sup>30</sup> |
| AMPDeep (2022) | PLMs based hemolytic activity prediction of antimicrobial peptides<br>(https://zenodo.org/records/6992526) | <sup>33</sup> |
| Hemolytic-Pred<br>(2023) | A web server for predicting hemolytic proteins utilizing position and composition-based features with an XGBoost model. (http://ec2-54-160-229-10.compute-1.amazonaws.com) | <sup>34</sup> |
| Ansari et al. (2023) | Serverless prediction of peptide properties using recurrent neural networks, focusing on classification tasks such as hemolysis, nonfouling, and solubility. (https://peptide.bio) | <sup>32</sup> |
| Mendieta et al.<br>(2023) | In Silico Discovery of Hemolytic Peptides Through a Novel Approach Based on Network Science and Similarity Searching Methods. | <sup>36</sup> |
| PeptideBERT (2023) | A transformer-based language model for peptide property prediction, specializing in classification tasks such as hemolysis, nonfouling, and solubility. (https://github.com/ChakradharG/PeptideBERT) | <sup>38</sup> |
| QSVM (2024) | Quantum machine learning (QML) based quantum support vector machine (QSVM) model to classify *hemolytic* and *non Hemolytic peptides*. | <sup>39</sup> |

In order to address challenges faced by the scientific community, we proposed a novel method for predicting hemolytic peptides as well as HC_50_ value against mammalian RBCs. The proposed method, HemoPI2, is trained and evaluated on experimentally validated 1926 hemolytic peptides. We have developed various classification models and regression models using ML, as well as PLMs. We have also developed ML-based models using word embeddings extracted from LLMs. These models have undergone rigorous benchmarking against independent dataset. Our HemoPI2 proposed in this study is an improved version of HemoPI, which has been widely utilized by the scientific community. Figure 1 provides a visual representation of the algorithm and processes undertaken in the study.

**Figure 1:**
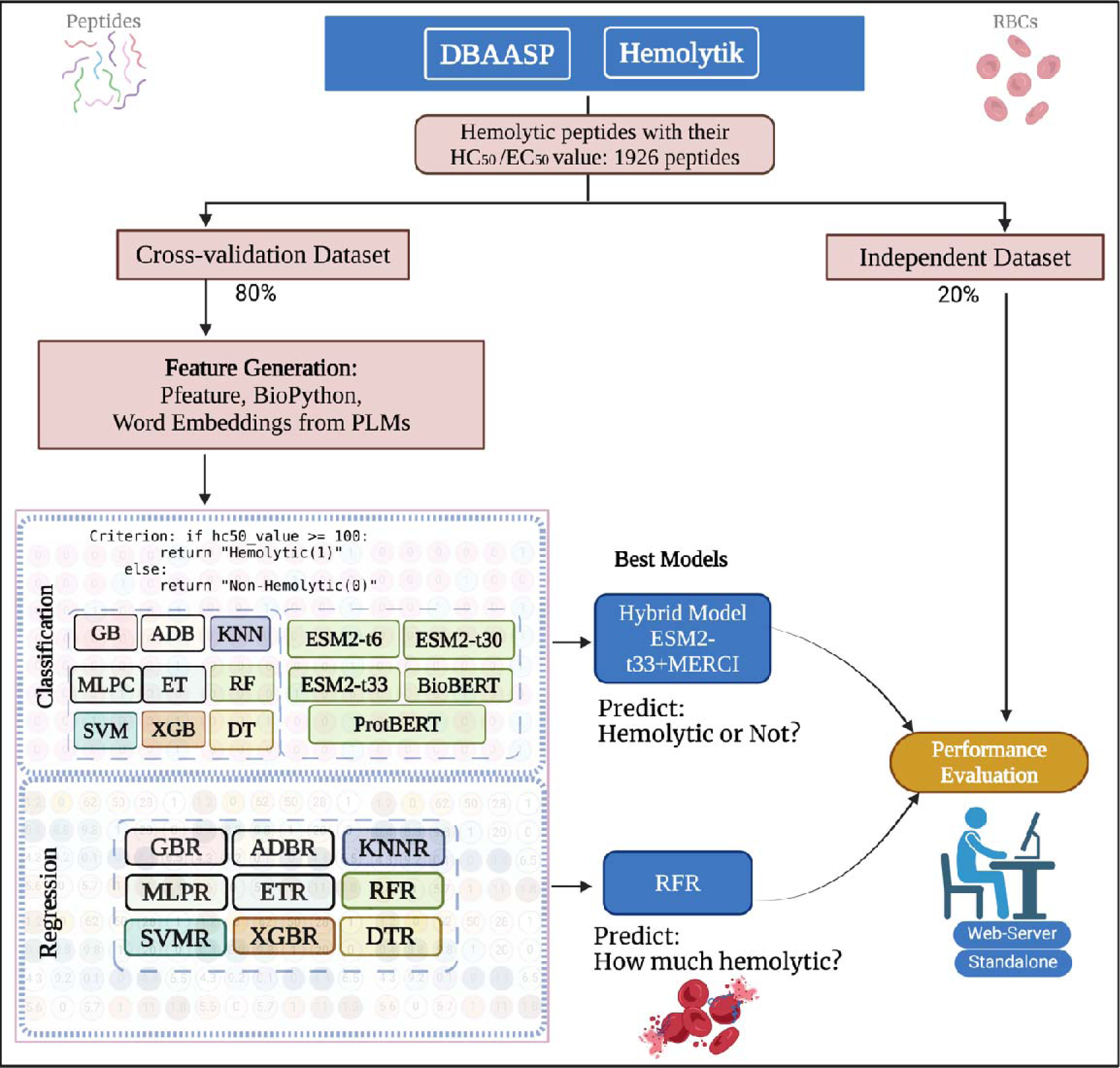
Illustration of the comprehensive workflow adopted throughout this study.

## 2 Methods

### 2.1 Data Collection

We acquired experimentally validated hemolytic peptides from DBAASP version 3 and Hemolytik database ^17,40^. These databases offer comprehensive details on the hemolytic activity of peptides that have undergone experimental validation. Peptide activity is assessed by extrapolating measurements from dose-response curves to determine the concentration at which 50% of RBCs are lysed, known as the HC_50_ value. We collected 3147 peptides from DBAASP and 560 peptides from the Hemolytik database, whose HC_50_ value is available. We implemented several preprocessing steps that included the removal of peptides containing non-natural amino acids and the removal of peptides containing less than six residues. In cases where a peptide sequence has multiple HC_50_ values or a range of HC_50_ values, we computed the average of these values. This mean activity measure represents the overall hemolytic activity of the peptide under various experimental conditions. By averaging, we ensure that our model captures the general behavior of the peptide’s hemolytic activity rather than specific instances, thereby enhancing the robustness of our predictions. The final HemoPI2 dataset comprises 1926 unique experimentally validated hemolytic peptides along with their corresponding hemolytic concentrations measured in μM, available as Supplementary Table S1. For classification purposes, we establish binary labels distinguishing strong hemolytic (positive) from weak hemolytic (negative) peptides; we utilized the criterion that peptides with an HC_50_ of ≤ 100 μM are classified as hemolytic. Peptides with HC_50_ values ≤ 100 μM were labeled as hemolytic, while those with values > 100 μM were classified as non-hemolytic. Following this criterion, we got 891 hemolytic peptides and 1035 non-hemolytic peptides in the dataset.

We standardized the HC_50_ values by converting them into a uniform measurement unit (µM). Following this, we transformed these HC_50_ values into pHC_50_ values using a specific equation (Equation. 1). This standardization and transformation process ensures consistency in our data, which is crucial for the accuracy and reliability of our regression model. It allows us to compare and analyze the hemolytic activity of different peptides on a common scale, thereby enhancing the predictive capabilities of our model.

#### Predictive Target for Regression Analysis

In this study, we have chosen the negative logarithmic HC_50_ (pHC_50_) as the target for our regression prediction. The pHC_50_ value is calculated using the following equation:

This equation transforms the HC_50_ values into a logarithmic scale, which can help in handling a wide range of HC_50_ values and can potentially improve the performance of the regression model. This transformation is commonly used in bioinformatics and cheminformatics for handling bioactivity data^41–43^. It allows us to compare and analyze the hemolytic activity of different peptides on a common scale, thereby enhancing the predictive capabilities of our model.

### 2.2 Cross-validation approach

This study followed established bioinformatics protocols. Initially, our data was randomly divided into training and independent datasets, with the cross-validation set comprising 80% of the data and the independent set containing the remaining 20%. We employed a five-fold cross-validation technique within the cross-validation dataset for training and testing to assess model performance. This process involved randomly dividing sequences into five subsets, using four for training and one for testing iteratively across five repetitions. Notably, the independent dataset was held aside throughout training, testing, and hyperparameter tuning. Only the final models were evaluated using the independent or unseen dataset. Comparison of models on this independent dataset is critical to any predictive methodology.

### 2.3 Feature extraction

In crafting a sequence-based predictor to elucidate the biological properties of peptides, a fundamental consideration is how to effectively represent the peptides to provide a comprehensive description of their features, reflecting their functions accurately. To extract features, we utilized the Pfeature^44^ tool, which yielded a diverse range of descriptors. These include AAC (20 descriptors), DPC (order 1) (400 descriptors), Atom Type Composition (5 descriptors), Bond Type Composition (4 descriptors), and Physico-chemical properties (30 descriptors), among others. Notably, this encompassed various indices and compositions, such as the Residue Repeats Index (20 descriptors), Property Repeats Index (19 descriptors), Distance Distribution of Repeats (20 descriptors), Shannon Entropy measures (40 descriptors), Conjoint Triad Descriptors (343 descriptors), and Composition enhanced Transition and Distribution (189 descriptors). In total, we gathered 1092 features for each peptide, which are the foundation for developing our predictive models.

### 2.4 Comprehensive analysis

To conduct the preliminary analysis of experimentally validated hemolytic peptides, we employed several analytical approaches. Initially, we conducted sequence-based analysis, which involved scrutinizing amino acid composition, positional distribution, and motif patterns. This exploration spanned the entire peptide sequences, including both the N-terminal and C-terminal regions. Following this, we employed Two Sample Logo (TSL)^45^ to discern specific preferences for amino acid residues at distinct positions within the peptides. In motif-based analysis, we employed the Motif-EmeRging and Classes-Identification (MERCI) tool^46^ to uncover recurring patterns contributing to hemolytic activity. Such motifs serve as pivotal regions within peptides responsible for their hemolytic effects. Identifying these motifs enhances our understanding of the molecular mechanisms underlying toxicity, offering insights for drug development and therapeutic strategies^47–49^. Lastly, we conducted a correlation analysis between features extracted by Pfeature and their corresponding HC_50_ concentrations. This facilitated the identification of features closely associated with peptide hemolytic activity, elucidating meaningful relationships crucial for further research^50^.

### 2.5 Word embeddings from protein language models

Recent strides in natural language processing (NLP) have catalyzed the emergence of PLMs, which harness individual amino acids and their combinations (doublets or triplets) as tokens or words. These models yield fixed-size vectors, referred to as embeddings, to encapsulate specific peptide sequences. These protein embeddings serve as pivotal inputs for a spectrum of tasks, spanning structure prediction, novel sequence generation, and protein classification^51^. In our study, we employed three widely recognized LLMs: ESM-2^52^, ProtBERT, and BioBERT, to produce embeddings for peptide sequences. ProtBERT and BioBERT are built upon the BERT model^53^ and are pre-trained on extensive datasets of protein sequences in a self-supervised manner. Conversely, ESM-2 (Evolutionary Scale Modeling) is a transformer-based PLMs initially developed for protein structure prediction, trained on sequences sourced from the UniRef protein sequence database^54^. ESM-2, renowned as a state-of-the-art protein model, is trained on a masked language modeling objective. This model proves adept at fine-tuning across an extensive array of tasks that entail protein sequences as inputs. Several ESM-2 checkpoints with varying sizes are available on HuggingFace, where larger sizes generally yield slightly better accuracy but necessitate significantly more memory and training time. We opted to utilize the ESM-2_t36_3B_UR50D, which consists of 36 transformer blocks with 3B parameters trained from UniRef50^55^, and ESM-2_t33_650M_UR50D^56^, which consists of 33 transformer blocks with 650M parameters trained from UniRef50^55^, checkpoints to generate embeddings. These checkpoints were deemed suitable for our objectives, offering a balance between accuracy and resource requirements. Subsequently, the embeddings derived from these ESM-2 checkpoints were employed as features for an ML regressor and classifier, facilitating the development of a robust model. This approach empowered us to exploit the rich contextual information encoded within the embeddings to enhance the predictive capabilities of the regressor and classifier models.

### 2.6 Classification Models

In our study, we employed a diverse set of ML classifiers to enhance the robustness and accuracy of our models. These include Extra Trees (ET), which are known for their ability to reduce over-fitting and bias; Support Vector Machines (SVM), effective in high-dimensional spaces; Extreme Gradient Boosting (XGB), renowned for its speed and performance; Random Forest (RF), appreciated for its handling of unbalanced datasets; Multi-Layer Perceptron Classifier (MLPC), a type of neural network known for its flexibility; Gradient Boosting (GB), recognized for reducing errors; and Decision Trees (DT) and Linear Regressor (LR), both fundamental to understanding feature importance and relationships. Each of these classifiers contributes unique strengths to our ML pipeline, resulting in a more robust and accurate predictive model.

#### 2.6.2 Protein language models

In our hemolytic peptide classification study, we utilized PLMs, which are computational frameworks that leverage natural language processing techniques to analyze protein structures, functions, and interactions. Specifically, we employed models such as ESM2-t33, ESM2-t30, and ESM2-t12 from the Evolutionary Scale Modeling (ESM) series, which are pre-trained on large protein sequence corpora and excel in tasks like structure prediction and variant effect prediction^57^. We also used BioBERT, a domain-specific model pre-trained on large-scale biomedical corpora^58^, and ProtBERT, a protein-specific model from the BERT series^59^, pre-trained on a vast corpus of protein sequences^60^. These models, each with their unique strengths, significantly enhanced the precision and reliability of our hemolytic peptide classification model.

### 2.7 Hybrid/Ensemble Model for classification

In our further investigation, we investigated a combined strategy, fusing an ensemble technique to bolster the predictive prowess of our concluding model. This amalgamated strategy merges a motif-based technique utilizing MERCI and models constructed using ML and PLMs methodologies. The initial step involved the application of the MERCI motif-based technique for peptide sequence categorization. Each prediction was allocated a weight: ‘+0.5’ for hemolytic predictions, ‘-0.5’ for non-hemolytic predictions, and ‘0’ when no matches were identified. This weighting scheme provides a numerical confidence indicator for each prediction. In the combined strategy, the scores from both the MERCI method and the ML were integrated to compute a comprehensive score. Depending on various threshold values, this cumulative score could classify the peptide sequence as toxic or non-toxic. This combined approach has been widely applied in numerous scientific investigations, validating its effectiveness in enhancing prediction accuracy.

### 2.8 Regression Models

To predict the HC_50_ of hemolytic peptides, we employed a diverse array of ML models along with PLMs. Our model development process involved leveraging various categories of features extracted by Pfeature. Additionally, we harnessed embeddings derived from fine-tuned PLMs, incorporating them as features for ML model development. This comprehensive approach enabled us to explore the predictive potential of both traditional features and advanced language model embeddings in modeling peptide hemolytic activity.

#### 2.8.1 Machine learning regressor models

The predictive performance of the computational models developed in this study relies not only on the chosen feature representations but also on the specific regression models employed. To explore the predictive capabilities comprehensively, we utilized a range of popular regressors, including XGBoost Regressor (XGBR), Random Forest Regressor (RFR), Gradient Boosting Regressor (GBR), Extra Trees Regressor (ETR), Decision Tree Regressor (DTR), AdaBoost Regressor (ADBR), Support Vector Regressor (SVR), K-Nearest Neighbors Regressor (KNNR), Linear Regressor (LR) and Multi-Layer Perceptron Regressor (MLPR).

Each of these regression models operates differently and leverages distinct mathematical algorithms to make predictions. For instance, XGBR is an implementation of gradient-boosted decision trees designed for speed and performance, while RFR utilizes an ensemble of decision trees to improve prediction accuracy and mitigate overfitting. GBR sequentially fits weak learners to the residuals of the previous models, gradually improving prediction accuracy. Similarly, ETR builds an ensemble of randomized decision trees to enhance prediction robustness further. DTR constructs a tree-like model of decisions based on feature inputs, recursively splitting data into subsets to minimize variance. ADBR combines multiple weak learners to create a strong learner iteratively, focusing on instances that previous models misclassified. SVR identifies the optimal hyperplane that maximizes the margin between data points and minimizes prediction error. KNNR predicts the output of a query point based on the majority vote of its k nearest neighbors, while LR assumes a linear relationship between input features and the target variable. These regression techniques have demonstrated success in predicting various functions and properties of peptides, as well as other biological or chemical entities in previous studies, as evidenced by the cited literature^61–64^.

### 2.9 Performance Metrics

The validation of the empirical predictive model is of paramount importance for evaluating its robustness. In the realm of pattern recognition, predicting hemolytic activity is approached as both a regression and classification problem. The regression analysis employs four standard statistical parameters. These include the Pearson Correlation Coefficient (R), which measures the linear correlation between predicted and actual values, and the Coefficient of Determination (R²), which indicates the fit of the data to the regression model (the closer to 1, the better the fit). Additionally, the Mean Absolute Error (MAE) provides the average of the absolute differences between predicted and actual values, while the Mean Squared Error (MSE) offers measures of the differences between these values. The formula of the following statistical parameters is given below:

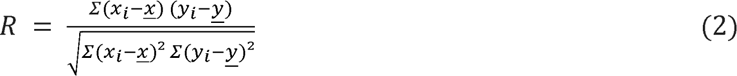

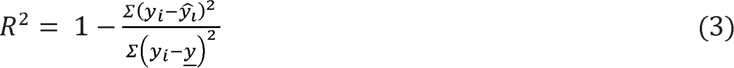

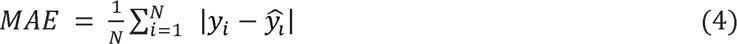

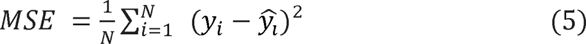

where *y_i_* and *x_i_* are the data points, N is the number of data points, 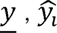, and 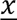 and indicates the mean value of y, the predicted value of actual value (y), and an average of x, respectively.

The efficacy of different classification models was gauged using established evaluation metrics, both threshold-dependent and independent. Threshold-dependent metrics, including sensitivity, specificity, accuracy, and the Matthews correlation coefficient (MCC), are influenced by the classification threshold. Conversely, the area under the receiver operating characteristic curve (AUC) is a threshold-independent metric that offers a holistic view of a model’s discriminative capacity. These metrics, extensively validated in previous studies, are crucial for reliable performance assessment.

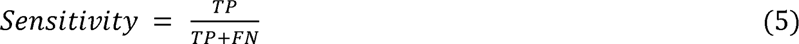

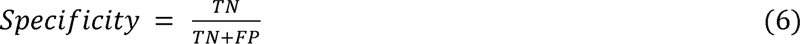

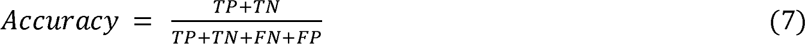

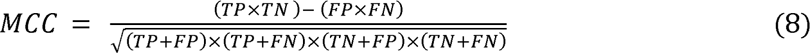

- True Positives (TP): Correctly predicted hemolytic peptides.
- False Positives (FP): Non-hemolytic peptides wrongly classified as hemolytic.
- True Negatives (TN): Correctly predicted non-hemolytic peptides.
- False Negatives (FN): Hemolytic peptides wrongly classified as non-hemolytic.

## 3 Results

The dataset included peptide sequences with experimentally validated hemolytic activity levels. We performed comprehensive analyses, including compositional, positional, and motif analyses, along with feature extraction using the Pfeature tool. Feature vectors are utilized to study their correlation with HC_50_ values and develop diverse ML models encompassing regression and classification. In addition to traditional ML models, a PLMs was also implemented. Embeddings were extracted from LLMs and utilized as feature vectors in the ML models. The predictive performance of these models was evaluated using an independent or unseen dataset, providing insights into their efficacy in predicting peptide hemolytic activity.

### 3.1 Analysis of Hemolytic peptides

#### 3.1.1 Amino acid composition analysis

The investigation into the amino acid composition of hemolytic and non-hemolytic peptides revealed distinct patterns that underscore their functional differences. In the comparative analysis in Figure 2, certain amino acid residues emerged as dominant features of hemolytic peptides, notably Cysteine, Phenylalanine, Glycine, and Leucine. These residues exhibited significantly higher proportions within hemolytic peptides compared to their non-hemolytic counterparts. Furthermore, the study noted the presence of other residues, such as Lysine and Tryptophan, albeit in lesser quantities, further distinguishing hemolytic peptides. Interestingly, the comparison extended to the termini composition, where the N and C termini exhibited almost identical overall compositions to the entire peptide. However, disparities surfaced in the distribution of Phenylalanine and Glycine, with heightened concentrations observed exclusively at the N-terminal end. Conversely, the C-terminal region displayed a near-equivalent distribution of Phenylalanine and Glycine between hemolytic and non-hemolytic peptides. These findings illuminate the nuanced amino acid profiles that contribute to the hemolytic properties of peptides, offering insights into their molecular mechanisms and potential applications in various biomedical contexts.

**Figure 2:**
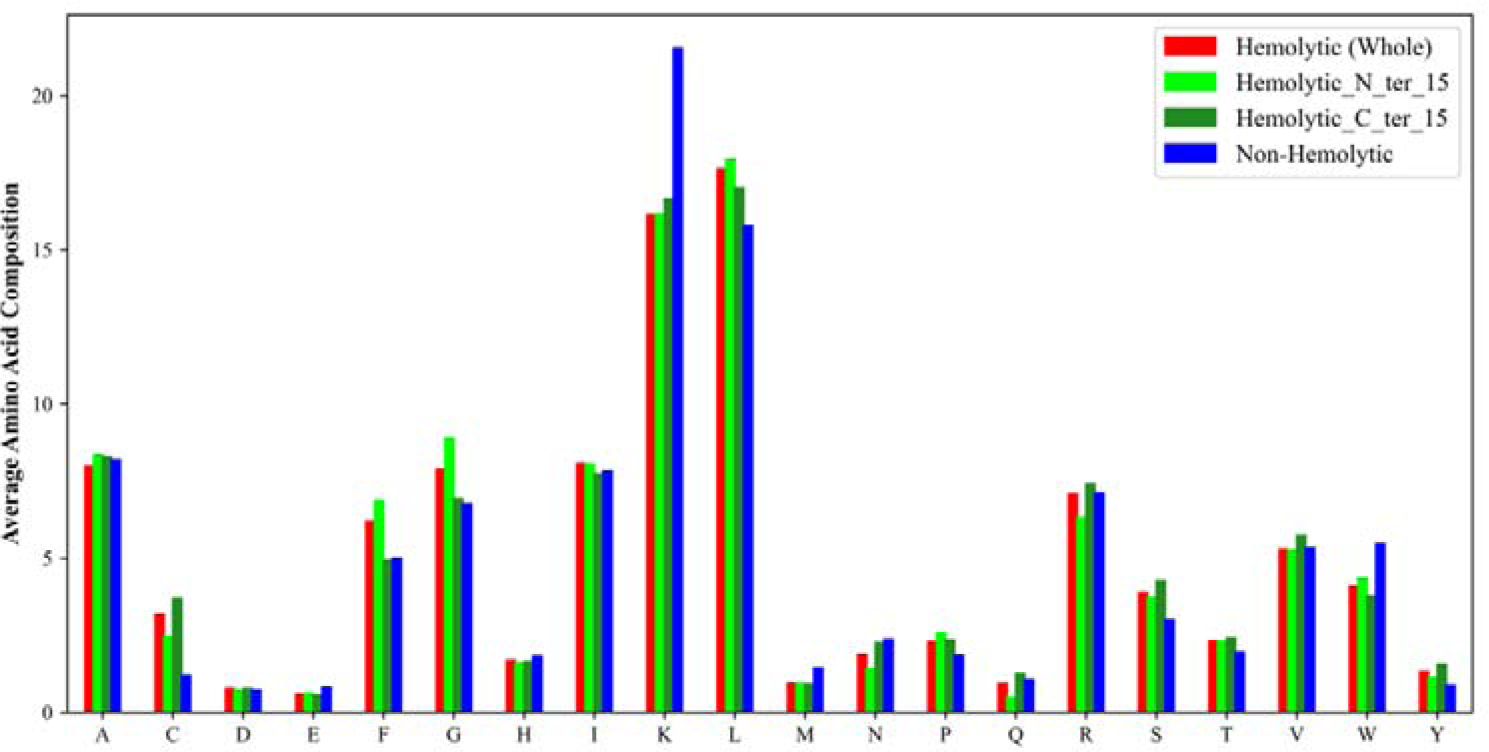
Representation of the average percentage composition of amino acid residues in various parts of experimentally validated peptides. Red, Cyan, and Green represent full sequences, N-terminal 15 residues, and C-terminal 15 residues of hemolytic peptides, respectively. Blue represents non-hemolytic peptides.

#### 3.1.2 Positional analysis

During our analysis, we aimed to identify any potential preferences for specific amino acid residues at particular positions within the peptide sequence. We constructed TSL for both hemolytic and non-hemolytic peptides, as illustrated in Figure 3. The TSL provides valuable insights into the relative abundance of amino acid residues and their significance within the sequence. Enriched residues are more prevalent at a given position in hemolytic peptides compared to the non-hemolytic class, while depleted residues are more prevalent at a given position in non-hemolytic peptides. Upon initial examination of the TSL, we observed findings consistent with the amino acid composition analysis: hemolytic peptides exhibited enrichment in hydrophobic residues and were predominantly depleted in positively charged residues. Further scrutiny revealed position-specific enrichments. In the N-terminal 15 residues (Figure 3A), hemolytic peptides were enriched in hydrophobic residues like Phenylalanine in position 1; Leucine in positions 2, 6, 8, and 12; Glycine in position 3; Isoleucine in position 5, and Proline in position 15, while in non-hemolytic peptides Lysine is preferred at most positions. Similarly, in the C-terminal 15 residues (Figure 3B), amino acids such as Alanine at position 1, Proline at position 5, Lysine at positions 6 and 14, and Arginine at positions 12 and 15 were more prevalent in hemolytic peptides, while Lysine is preferred in most positions but at positions 6 and 14 Alanine is preferred in non-hemolytic peptides. These observations shed light on the differential distribution of amino acids along the peptide sequence and provide valuable insights into the structural and functional characteristics of hemolytic as well as non-hemolytic peptides.

**Figure 3:**
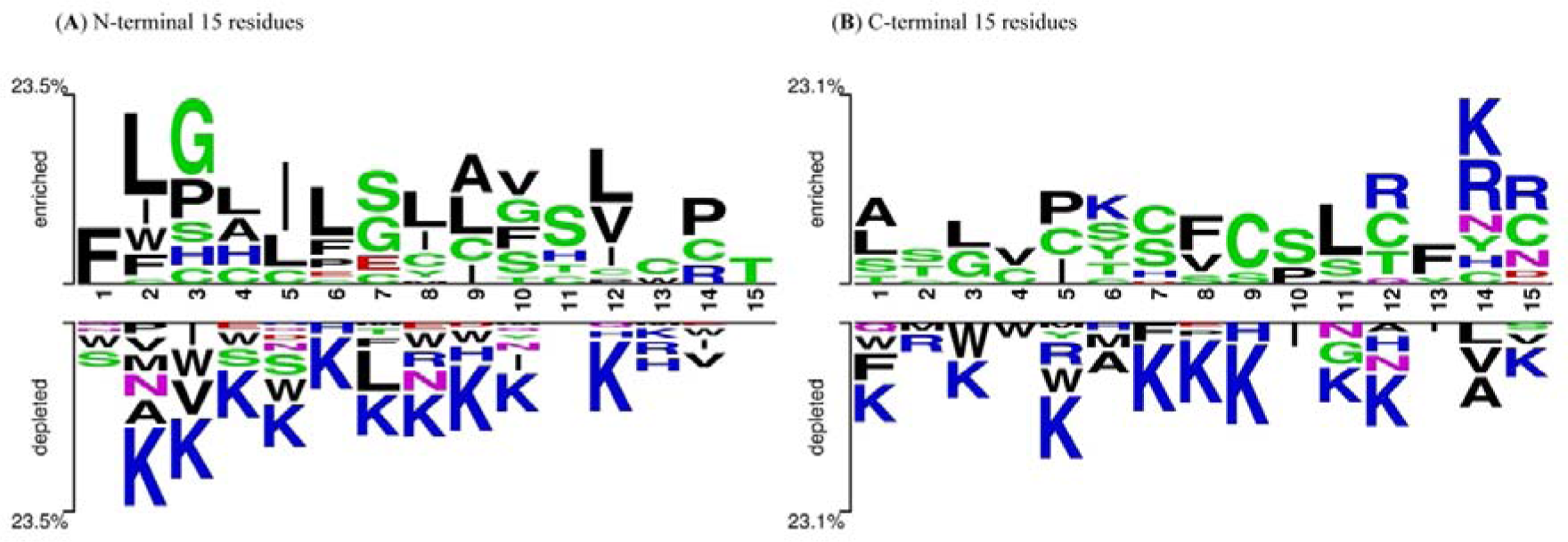
This illustration presents TLS that depict the residue preferences in hemolytic and non-hemolytic peptides. Logo A) represents the first 15 residues of the N-terminal, while Logo B) represents the last 15 residues of the C-terminal.

#### 3.1.3 Motif analysis

Motif analysis was conducted to pinpoint motifs present exclusively in either hemolytic or non-hemolytic peptides. This analysis identifies specific segments or patterns within peptides that contribute to their hemolytic activity. Consequently, motif analysis not only enhances prediction accuracy but also aids in identifying the precise motif or region responsible for hemolysis. The top ten motifs unique to hemolytic peptides include ‘CGET’, ‘CGETC’, ‘TLLKKVLKA’, ‘TPGC’, ‘AIH’, ‘GETC’, ‘TLLKKVLKA’, ‘GGLFS’, ‘IGGLF’ and ‘TLLKK’.

Conversely, the top ten motifs exclusive to non-hemolytic peptides consist of ‘AKD’, ‘SKIK’, ‘DLA’, ‘SKIKK’, ‘HVQ’, ‘NKL’, ‘HRK’, ‘INKQ’, ‘KDLA’ and ‘KINKQ’. Supplementary Table S2 contains the full list of motifs.

#### 3.1.4 Correlation analysis

Past research shows correlation analysis is crucial in predicting protein functions, understanding disease mechanisms, and discovering drugs^65^. It quantifies the strength and direction of relationships between variables, highlighting key influencers and potential patterns. In Table 2, we have highlighted the top features of AAC, DPC, and physico-chemical properties that exhibit a correlation with the HC_50_ concentration of hemolytic peptides. Features that exhibit a positive correlation with HC_50_ concentration of hemolytic peptides indicate that as specific attributes (e.g., composition of positively charged residues) increase, the HC50 value of hemolytic peptides also rises, suggesting a reduction in hemolytic potency. On the other hand, a negative correlation (e.g., the composition of neutral and hydrophobic residues) indicates that an increase in a feature’s value leads to a decrease in the HC_50_ value, implying an increase in hemolytic potency. A comprehensive list of correlation analysis of each feature is provided in Supplementary Table S3.

**Table 2.** Top amino acid, di-peptide, and physico-chemical features correlated to the HC_50_ value of hemolytic peptides.

| **AAC Features** | **Correlation** | **DPC Features** | **Correlation** | **Physico-chemical Features** | **Correlation** |
| --- | --- | --- | --- | --- | --- |
| AAC_K | 0.214 | DPC1_KK | 0.207 | Positively charged residues | 0.236 |
| AAC_R | 0.043 | DPC1_IK | 0.191 | Basic residues | 0.236 |
| AAC_W | 0.018 | DPC1_AK | 0.165 | Hydrophilic residues | 0.220 |
| AAC_A | 0.011 | DPC1_LY | 0.143 | Exposed | 0.166 |
| AAC_Y | 0.010 | DPC1_YR | 0.101 | Helix | 0.157 |
| AAC_P | -0.075 | DPC1_FL | -0.088 | Hydrophobic residues | -0.154 |
| AAC_C | -0.075 | DPC1_VL | -0.091 | Buried residue | -0.167 |
| AAC_S | -0.092 | DPC1_GG | -0.094 | PCP_Z3 | -0.204 |
| AAC_F | -0.093 | DPC1_LF | -0.102 | Neutral charged residues | -0.233 |
| AAC_G | -0.102 | DPC1_IG | -0.132 | Neutral residues based on pH | -0.233 |

### 3.2 Classification Models

We developed various classifiers using a combination of ML techniques and PLMs. In order to develop a classification model, we employed a dataset comprising 1924 distinct experimentally validated hemolytic peptides. We assigned binary labels to differentiate between strong hemolytic (positive) and weak hemolytic (negative) peptides. Our classification criteria categorized peptides with an HC_50_ of ≤ 100 μM as hemolytic.

#### 3.2.1 Machine Learning Models

We developed various classifiers to classify high hemolytic peptides and weak hemolytic peptides, including ET, SVM, XGBC, RF, MLPC, GB, DT, and LR. Initially, we calculated the features of the hemolytic peptides utilizing the compositional-based module of Pfeature. Additionally, molecular weight and peptide length were included as feature vector components. This procedure yielded a comprehensive set of 1092 feature vectors for each peptide. Once again, the tree-based classifiers outperformed other classifiers, particularly RF. In Table 3, the performance was compared based on the AUC using different features and evaluated on an independent dataset. RF classifier achieved the best performance on the ALLCOMP excluding SOC (1090 descriptors) with an AUC of 0.888, which is very close to the performance on combined features AAC+DPC+PCP (450 descriptors) with an AUC of 0.881, AAC+DPC (420 descriptors) with an AUC of 0.875, and CeTD (189 descriptors) with an AUC score of 0.856. The detailed evaluation metrics are provided in Supplementary Table S4.

**Table 3:** Evaluation performance metrics of RF classifier model on independent dataset using various features derived from Pfeature and ProtPram.

| Features | Sp (%) | Sn (%) | Acc (%) | MCC | AUC |
| --- | --- | --- | --- | --- | --- |
| AAC | 83.9 | 66.3 | 75.9 | 0.512 | 0.831 |
| DPC | 87.7 | 65.1 | 77.5 | 0.547 | 0.837 |
| PCP | 84.8 | 67.4 | 76.9 | 0.534 | 0.854 |
| CeTD | 86.7 | 72.0 | 80.1 | 0.597 | 0.856 |
| AAC+DPC | 74.9 | 85.5 | 80.6 | 0.609 | 0.875 |
| AAC+DPC+CeTD | 76.2 | 78.8 | 77.5 | 0.598 | 0.841 |
| AAC+DPC+PCP | 73.7 | 85.0 | 79.8 | 0.623 | 0.881 |
| ALLCOMP | 74.9 | 83.9 | 80.1 | 0.610 | 0.878 |
| **ALLCOMP-ex SOC** | **76.5** | **86.9** | **82.1** | **0.640** | **0.888** |

#### 3.2.2 Protein Language Models

Our study evaluated the performance of various PLMs in classifying hemolytic peptides, measured by the evaluation metric. The results highlight the effectiveness of different PLMs in accurately classifying hemolytic peptides, with ProtBERT exhibiting particularly strong performance with AUC 0.875 (See Table 4). The varied AUC values provide insights into the relative strengths of each model in our classification task, allowing for informed model selection and optimization in future studies. Table 4 shows the detailed performance of each model.

**Table 4:** Comprehensive performance analysis of PLMs on an independent dataset.

| PLM | Sp (%) | Sn (%) | Acc (%) | MCC | AUC |
| --- | --- | --- | --- | --- | --- |
| ESM2-t33 | 86.1 | 70.0 | 79.1 | 0.573 | 0.870 |
| ESM2-t30 | 84.2 | 78.0 | 81.3 | 0.599 | 0.851 |
| ESM2-t12 | 70.0 | 79.0 | 74.2 | 0.490 | 0.831 |
| ESM2-t6 | 76.1 | 84.0 | 79.5 | 0.591 | 0.870 |
| **ProtBERT** | **84.9** | **73.7** | **80.0** | **0.602** | **0.875** |
| BioBERT | 63.7 | 80.5 | 71.5 | 0.474 | 0.800 |

#### 3.2.3 ML models based on word embeddings

Word embeddings were generated using various checkpoints of ESM2 (ESM2-t36, ESM2-t33, ESM2-t30, ESM2-t6), ProtBERT, BioBERT, and ProtBERT + BiLSTM, mirroring the approach adopted to develop regression models. The ML classifiers, including ET, SVM, XGB, RF, MLPC, GB, and DT, were trained to predict the hemolytic activity of peptides. MLPC with ProtBERT embedding achieved maximum performance with AUC 0.882 (See Table 5)

**Table 5:** Evaluation of ML classifiers’ performance utilizing various PLM embedding sources on an independent dataset.

| **Embedding Source** | **ML classifier** | **Sp (%)** | **Sn (%)** | **Acc (%)** | **MCC** | **AUC** |
| --- | --- | --- | --- | --- | --- | --- |
| ESM2-t36 | XGBC | 74.9 | 86.0 | 80.8 | 0.614 | 0.869 |
| ESM2-t33 | ET | 73.2 | 85.5 | 79.8 | 0.604 | 0.873 |
| ESM2-t6 | XGBC | 74.3 | 79.7 | 77.2 | 0.541 | 0.865 |
| **ProtBERT** | **MLPC** | **84.0** | **83.9** | **83.9** | **0.677** | **0.882** |
| BioBERT | MLPC | 68.5 | 81.5 | 75.6 | 0.507 | 0.792 |
| ProtBERT+BiLSTM | ET | 71.4 | 82.8 | 77.4 | 0.547 | 0.859 |

### 3.3 Hybrid model for classification

In order to boost the predictive power of our top-performing model, we adopted a hybrid strategy for the classification of hemolytic peptides. This strategy involved the use of a weighted scoring technique to construct hybrid models based on motifs, integrating MERCI with ML and PLMs. Each prediction made by the MERCI method was assigned a weight: ‘+0.5’ for hemolytic predictions, ‘-0.5’ for non-hemolytic predictions, and ‘0’ when no matches were found. This weighting scheme provides a numerical confidence indicator for each prediction, allowing us to quantify the confidence in the prediction made by ML and PLMs, leading to a substantial enhancement in the AUC, a key metric for evaluating the performance of our predictive models. Simultaneously, we carefully calibrated the threshold value integral to this classification. This pivotal element strikes a balance between sensitivity (true positive detection) and specificity (avoidance of false positives). Adjusting this threshold allowed us to optimize the model’s performance, enhancing sensitivity at the cost of reduced specificity or vice versa. Table 6 presents the performance of the best-performing model when combined with the MERCI motif-based method, evaluated on an independent dataset. Among the ML models, the RF model, when used with ALLCOMP features with an AUC of 0.906, and among the PLMs, ESM2-t6 achieved the best performance with an AUC score of 0.909. It’s worth noting that an AUC score of 0.90 is considered excellent, suggesting that these models have a high true positive rate and a low false positive rate. This is a significant improvement in the field of peptide classification. Supplementary Table S5 provides detailed performance metrics of hybrid models.

**Table 6:** Performance of hybrid model on independent dataset developed by combining best-performing classification models with MERCI.

| Model | Threshold | Sp | Sn (%) | Acc (%) | MCC | AUC |
| --- | --- | --- | --- | --- | --- | --- |

|  |  | **d** | **(%)** |  |  |  |  |
| --- | --- | --- | --- | --- | --- | --- | --- |
| **ML** | **RF**<br>**(ALLCOMP**<br>**-SOC)** | **0.46** | **81.6** | **84.0** | **82.6** | **0.651** | **0.906** |
|  | RF<br>(AAC+DPC<br>+PCPF) | 0.47 | 79.9 | 83.6 | 81.9 | 0.641 | 0.890 |
|  | XGBC<br>(ESM2t36<br>embeddings) | 0.50 | 78.8 | 86.0 | 82.6 | 0.651 | 0.892 |
|  | ET<br>(ESM2t33<br>embeddings) | 0.49 | 81.6 | 81.6 | 81.6 | 0.631 | 0.899 |
|  | MLPC<br>(ProtBERT<br>embeddings) | 0.48 | 76.5 | 87.4 | 82.4 | 0.646 | 0.892 |
|  | ET<br>(ESM2t6<br>embeddings) | 0.49 | 77.7 | 86.5 | 82.4 | 0.646 | 0.892 |
| **PLM** | ESM2-t33 | 0.46 | 68.7 | 94.2 | 82.4 | 0.658 | 0.890 |
|  | ESM2-t36 | 0.48 | 78.7 | 79.7 | 79.2 | 0.584 | 0.875 |
|  | **ESM2-t6** | **0.55** | **82.1** | **83.7** | **83.4** | **0.657** | **0.909** |
|  | ProtBERT | 0.46 | 65.9 | 90.3 | 79.0 | 0.585 | 0.871 |

### 3.4 Regression Models

Initial investigations have suggested the feasibility of distinguishing hemolytic peptides from non-hemolytic ones based on factors such as amino acid composition, binary profiles, motifs, and physicochemical properties. Our novel study utilized several popular ML regressors to predict the hemolytic activity of peptides using features derived from their primary sequences. The predictive models underwent training and testing via fivefold cross-validation on the training dataset (80%), with the final models being evaluated using an independent dataset (20%). We reported key statistical parameters, including R, R^2^, MAE, and MSE.

#### 3.4.1 ML Regressors using Pfeature

In this study, we developed prediction models using a range of regressors, including XGBR, RFR, GBR, ETR, DTR, ADBR, SVR, KNNR, and LR. The various features we used for the regression models are the same as those used for the classification models. This comprehensive approach ensures the robustness and accuracy of our hemolytic activity prediction model. While experimenting with various categories of features and feature combinations, tree-based regressors such as RFR and ETR consistently emerged as the top performers among the various ML algorithms we evaluated. Detailed performance of each model is provided in Supplementary Table S6. Although the performance of RFR slightly outshone ETR, the difference was marginal. Table 7 displays the performance of the RFR with different sets of features on an independent dataset. The RFR demonstrated its superior performance when employing the ALLCOMP excluding SPC (1167 descriptors), achieving a R of 0.739 and an R^2^ value of 0.543.

**Table 7:** Performance of RFR-based model developed utilizing various composition-based features of hemolytic peptides on an independent dataset.

| Features | R | R<sup>2</sup> | MAE | MSE |
| --- | --- | --- | --- | --- |
| AAC | 0.681 | 0.456 | 0.826 | 1.168 |
| DPC | 0.702 | 0.469 | 0.837 | 1.139 |
| PCP | 0.697 | 0.477 | 0.798 | 1.123 |
| CeTD | 0.716 | 0.507 | 0.769 | 1.058 |
| AAC+DPC | 0.719 | 0.515 | 0.772 | 1.043 |
| AAC+DPC+CeTD | 0.735 | 0.531 | 0.752 | 1.007 |
| AAC+DPC+CeTD+PCP | 0.729 | 0.532 | 0.730 | 1.004 |
| ALLCOMP | 0.737 | 0.540 | 0.733 | 0.986 |
| **ALLCOMP -ex SPC** | **0.739** | **0.543** | **0.734** | **0.981** |

#### 3.4.2 ML models based on word embeddings

Over the past decade, the application of language models in the field of bioinformatics has seen a significant surge. A multitude of protein and nucleotide language models have been developed with the aim of predicting the function of biological macromolecules^66–68^. In this study, we employed PLMs such as ESM-2 (with checkpoints: esm2_t36_3B_UR50D, esm2_t33_650M_UR50D, esm2_t30_150M_UR50D, and esm2_t6_8M_UR50D), ProtBERT, and BioBERT to predict the HC_50_ of hemolytic peptides. Given that these models are not specifically tuned for any particular property of peptides, we optimized the hyperparameters of these models on our dataset of hemolytic peptides for classification tasks. Subsequently, these tuned models were utilized to extract embeddings from hemolytic peptides. These embeddings were then used as input for ML regression models.

We generated 2560, 1280, 640, 320, 1024, and 767 embeddings from ESM2-t36, ESM2-t33, ESM2-t30, ESM2-t6, ProtBERT, and BioBERT, respectively, for each peptide. In the case of ProtBERT, we also employed a combination of a fine-tuned ProtBERT model with BiLSTM to extract high-quality embeddings. These embeddings were used as features for developing ML models to predict the HC_50_ peptides. Table 8 illustrates the performance of the top-performing ML regression model with the corresponding PLMs on an independent dataset, utilizing embeddings derived from various language model architectures. Once more, among the array of ML regressors, tree-based methods like the EFR and ETR consistently stood out as strong performers. ESM embeddings, particularly those generated by ESM2-t33, demonstrated the best performance, especially with the Extra Trees Regressor achieving an R of 0.711 and an R^2^ of 0.495.

**Table 8:**
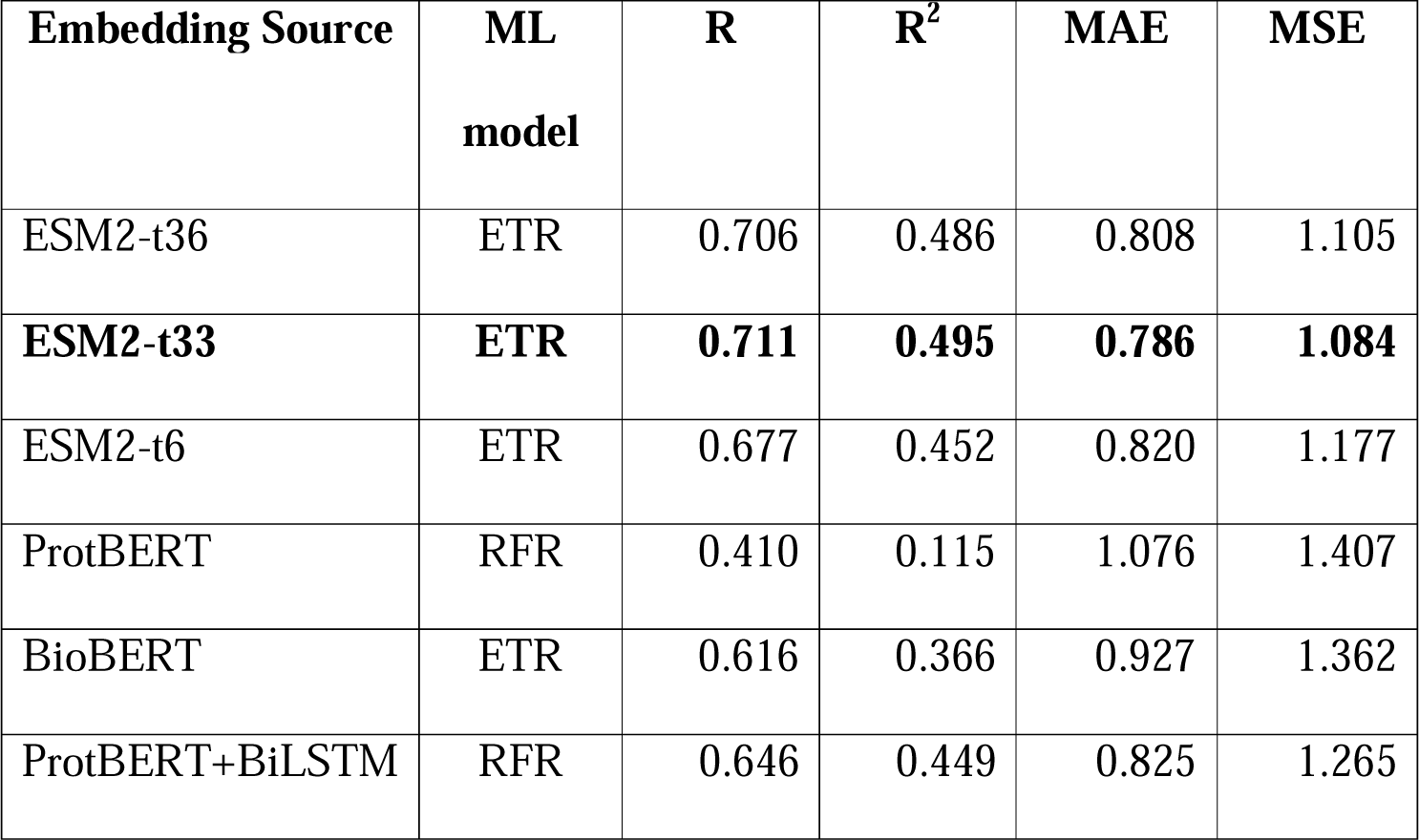
Evaluation of ML regressor models constructed using word embeddings derived from PLMs on an independent dataset.

| Embedding Source | ML model | R | R<sup>2</sup> | MAE | MSE |
| --- | --- | --- | --- | --- | --- |
| ESM2-t36 | ETR | 0.706 | 0.486 | 0.808 | 1.105 |
| **ESM2-t33** | **ETR** | **0.711** | **0.495** | **0.786** | **1.084** |
| ESM2-t6 | ETR | 0.677 | 0.452 | 0.820 | 1.177 |
| ProtBERT | RFR | 0.410 | 0.115 | 1.076 | 1.407 |
| BioBERT | ETR | 0.616 | 0.366 | 0.927 | 1.362 |
| ProtBERT+BiLSTM | RFR | 0.646 | 0.449 | 0.825 | 1.265 |

### 3.5 Model Finalization

In our research, we constructed several regression and classification models, utilizing a diverse set of features derived from Pfeature. Additionally, we incorporated word embeddings extracted from PLMs as input features. We experimented with various combinations of these features to optimize our models. All developed models were ultimately evaluated on an independent dataset, and their performances were compared. Among all the regression models, the RFR model exhibited superior performance with an R 0.739 and an R^2^ 0.543. A comparative analysis of the performance of the best ML and PLM regression models is presented in the tables.

In our classification model development, we diversified our approach similar to the regression models. Notably, we observed promising performance with RF (ALLCOMP-ex SOC) and ProtBERT models. Moreover, we extended our classification methodologies by integrating hybrid approaches that combine ML and PLM classifiers with MERCI. These hybrid strategies were meticulously crafted to leverage the complementary strengths of both methods. Subsequent evaluation revealed enhancements in model performance following the incorporation of MERCI. A comparative assessment of these hybrid models on independent datasets is detailed in the table. The ESM2-t6 and RF (ALLCOMP-ex SOC) models are particularly noteworthy, which demonstrated optimal performance with AUC scores of 0.909 and 0.906, respectively. These regression and classification models have been seamlessly integrated into our prediction software and web services, streamlining the quantification and classification of hemolytic peptides.

### 3.6 Benchmarking

Benchmarking a newly developed method against existing ones is crucial to comprehend its significance and potential improvements. In the context of a regression model, to the best of our knowledge, no tool has been specifically designed to quantify hemolytic peptides. However, several tools have been developed for classification (as detailed in Table 1). We conducted an evaluation of these current tools using an independent/unseen dataset from our study. This evaluation provides a comparative analysis with other existing approaches, offering insights into their relative effectiveness. In Table 6, we present a comparison of the proposed approach, HemoPI2.0, with other methods currently in use, as reported in the literature. In the independent dataset employed in HemoPI2.0, HAPPENN, PeptideBERT, and Plisson et al. achieved AUC values of 0.736, 0.739, and 0.740, respectively. However, HemoPI2.0 notably outperformed the others, demonstrating an AUC of 0.909. (See Table 9) The subpar performance of tools like HemoPI and HemoPred could potentially be attributed to the utilization of randomly generated negative datasets during training. Furthermore, we faced challenges in comparing the performance of certain tools such as HLPpred-FUSE, HemoNet, haemolytic-Pred, and RNN (Capecchi et al.) due to limitations in their services. Specifically, HLPpred-FUSE and haemolytic-Pred suffered from improper functioning of their web services, while HemoNet and RNN (Capecchi et al.) had non-functional GitHub code due to the absence of input files.

**Table 9:** Benchmarking of classification tools on an independent dataset of HemoPI2.0.

| Tools | Model | Sp (%) | Sn (%) | Acc (%) | MCC | AUC |
| --- | --- | --- | --- | --- | --- | --- |
| HemoPI | D1 | 16.7 | 90.8 | 58.4 | 0.099 | 0.538 |
|  | D2 | 59.9 | 60.8 | 60.3 | 0.207 | 0.662 |
|  | ModelAll | 49.2 | 69.8 | 58.8 | 0.194 | 0.669 |
| HemoPred | Default | 31.5 | 87.0 | 57.2 | 0.221 | - |
| HAPPENN | Main_Dataset | 53.7 | 84.9 | 68.1 | 0.402 | 0.736 |
| AMP_deep | Prot-BERT-BFD | 67.0 | 53.0 | 59.0 | 0.413 | 0.602 |
| Ansari et al. | Embedding + Bi-LSTM | 73.3 | 32.3 | 54.5 | 0.07 | 0.512 |
| PeptideBERT | Default | 79.1 | 62.4 | 71.0 | 0.422 | 0.739 |
| Plisson et al. | XGBC | 69.0 | 61.4 | 70.0 | 0.470 | 0.740 |
| **HemoPI2.0**<br>**(Proposed model)** | **ESM2-t6** | **83.7** | **82.1** | **83.4** | **0.657** | **0.909** |
|  | **RF** | **84.0** | **81.6** | **82.6** | **0.651** | **0.906** |

The data clearly demonstrates that HemoPI2.0 outperforms the alternative existing methods. This superior performance underscores the potential of HemoPI2.0 as a valuable tool in the field of therapeutic peptide development, particularly in the classification and quantification of hemolytic peptides.

### 3.7 Community contribution through HemoPI2.0

To contribute to the scientific community, we have made our regression and classification algorithms accessible online via our user-friendly web server, HemoPI2.0, which is freely available for academic researchers at https://webs.iiitd.edu.in/raghava/hemopi2/. The web server is organized into four main modules: Home, Prediction, Protein scanning, Motif scan, and Design. The prediction modules enable users to predict the hemolytic potency of their peptides by submitting multiple peptide sequences in FASTA format. However, predictions are restricted to peptides composed of the 20 natural amino acids, with non-natural amino acids not being supported. The protein scanning module provides the functionality to identify or scan toxic regions within a protein. The motif scan module enables users to scan or map hemolytic motifs within the query sequence using MERCI. The design module creates non-toxic peptides from the primary sequence, generating mutant peptides with a single mutation for hemolytic activity prediction. Additionally, Download and Help modules are available to facilitate data download and provide user assistance. Alongside the web server, we have also developed a standalone package that can be downloaded from our web server. This standalone software and pip package, designed for large-scale hemolytic peptide prediction, offers a versatile solution for comprehensive analysis.

## 4. Discussion

In recent decades, the focus on therapeutic peptide development has intensified. However, the inherent toxicity of many potential peptide lead molecules, specifically hemolytic activity, often hinders their progression into drug molecules. Most clinically approved therapeutic peptide drugs exhibit a high therapeutic index, meaning they possess high therapeutic activity and minimal hemolytic activity. Yet, a significant number of potential therapeutic peptides have been found to exhibit varying degrees of hemolysis. Therefore, the development of a drug with a high therapeutic index is of paramount importance. Over the years, the spotlight has primarily been on the classification of hemolytic peptides. As mentioned in Table 1, numerous tools have been developed for this purpose. To expedite the lead molecule design and optimization pipeline, this study aimed to create an improved in silico method for not only classifying but also quantifying therapeutic peptides as hemolytic or non-hemolytic based on their primary sequence. This approach is expected to significantly contribute to the field of therapeutic peptide development by providing a more nuanced understanding of peptide toxicity, thereby facilitating the design of safer and more effective therapeutic peptides.

The efficacy and dependability of an ML classifier heavily rely on the accurate delineation of its task. In this instance, the task involves defining positive and negative datasets sourced from the DBAASP (version: 3) and Hemolytik databases, containing experimentally validated peptide sequences. Prioritizing peptides with HC_50_ values, indicative of the concentration leading to 50% red blood cell lysis, was part of the selection criteria. Peptides with non-natural amino acids or fewer than six residues were excluded from upholding dataset integrity. For peptides with multiple or ranged HC_50_ values, averaging was employed to bolster prediction robustness. This meticulous process culminated in the creation of the HemoPI2.0 dataset, housing 1924 unique peptides alongside their respective hemolytic concentrations (in μM). This comprehensive dataset lays a strong groundwork for prediction model development. For feature extraction, we employed Pfeature, along with PLMs, to extract embeddings from peptide sequences for ML model development. Regression models, including ET, RF, DT, SVM, XGB, GB, MLPC, and LR, were assessed alongside PLMs like ESM2-t36, ESM2-t33, ESM2-t30, ESM2-t6, ProtBERT, and BioBERT, with performance evaluated on an independent dataset. RFR excelled in regression, while for classification, peptides with HC_50_ ≤ 100 μM were classified as hemolytic. RF and ProtBERT were top performers deployed on the web server. A hybrid model incorporating motif information improved performance. Leading regression and classification models are integrated into the web server.

Various analytical methods were employed for the preliminary analysis of experimentally validated hemolytic peptides. This included sequence-based scrutiny of AAC, positional distribution, and motif patterns across the peptide sequences. TSL were utilized to identify amino acid residue preferences at different positions. Motif-based analysis using MERCI tool revealed recurring patterns contributing to hemolytic activity, aiding in understanding molecular mechanisms. Correlation analysis between Pfeature-extracted features and HC_50_ concentrations helped identify features closely linked to peptide hemolytic activity, offering insights for further research. The amino acid composition analysis revealed distinct patterns distinguishing hemolytic and non-hemolytic peptides, with certain hydrophobic residues such as Cysteine, Phenylalanine, Glycine, and Leucine being predominant in hemolytic peptides. Terminally, the composition was similar, yet differences arose in Phenylalanine and Glycine distribution. Positional analysis through Two Sample Logos highlighted enriched hydrophobic residues in hemolytic peptides, particularly in the N-terminal region, while motif analysis identified unique motifs like ‘CGET’ and ‘TLLKKVLKA’ in hemolytic peptides and ‘AKD’ and ‘SKIK’ in non-hemolytic ones. Furthermore, correlation analysis showcased features like the composition of positively charged residues (AAC_K, AAC_R, DPC1_KK, etc.) and composition of basic residues positively correlating with HC_50_ concentration in hemolytic peptides, suggesting reduced hemolytic potency with increasing AAC_K values, and conversely, a negative correlation indicating increased hemolytic potency with certain physico-chemical properties like composition of hydrophobic residues (AAC_G, DPC1_IG, etc.). These insights underscore the diverse molecular characteristics influencing peptide hemolytic activity, informing potential applications in biomedical research and therapeutic development.

Our analysis underscores the significant role of hydrophobicity in influencing hemolytic activity. These findings are consistent with established literature, revealing that peptides with higher hydrophobicity tend to exhibit greater hemolytic potential compared to less hydrophobic counterparts. The preference for hydrophobic amino acids, such as Cysteine, Phenylalanine, Glycine, and Leucine, in hemolytic peptides can be attributed to their unique chemical properties and structural functions. These residues, composed mainly of carbon and hydrogen, possess minimal dipole moments and are naturally repelled from water^69^. This hydrophobic nature drives their affinity for lipid bilayers, the primary components of cell membranes, leading to membrane insertion and disruption, ultimately causing cell lysis.

Additionally, the amphipathic nature of many hemolytic peptides, with hydrophobic and hydrophilic regions, enhances their membrane-disrupting abilities^70,71^. Notably, peptides containing cysteine within β-sheets have been linked to increased hemolytic activity^72–74^. Furthermore, interactions between hydrophobic residues and specific lipid components or membrane proteins may enable selective targeting of cell types, enhancing hemolysis efficiency while minimizing interactions with non-target cells. Overall, the prevalence of hydrophobic amino acids in hemolytic peptides underscores their crucial role in membrane interactions, membrane integrity disruption, and hemolysis initiation.

## 5. Conclusion

HemoPI2.0 stands out as an innovative computational method capable of accurately quantifying and categorizing peptide hemolytic tendencies in peptides solely from their primary sequences. Developed exclusively on experimentally validated hemolytic peptides, this model fulfills a critical need for a robust computational tool that can swiftly and dependably predict peptide hemolytic potency. Its accessibility as a free, user-friendly, and publicly available web service further enhances its utility for diverse applications, spanning from fundamental research to drug discovery endeavors. A comprehensive analysis of various features sheds light on the factors influencing the hemolytic potency of peptides, offering crucial insights for designing peptide-based drugs with high therapeutic index.

## List of Abbreviations

| Abbreviation | Definition |
| --- | --- |
| RBCs | Red blood cells |
| HC<sub>50</sub> | Hemolytic concentration that lyse 50% of RBCs |
| FDA | Food and Drug Administration |
| ADMET | Absorption, distribution, metabolism, excretion, and toxicity |
| ML | Machine learning |
| LLM | Large language model |
| PLMs | Protein language models |
| AAC | Amino acid composition |
| DPC | Di-peptide composition |
| ESM | Evolutionary scale modeling |
| TSL | Two sample logo |
| Sp | Specificity |
| Sn | Sensitivity |
| Acc | Accuracy |
| MCC | Matthews correlation coefficient |
| AUC | Area under receiver operating characteristic |
| R | Pearson Correlation Coefficient |
| R<sup>2</sup> | Coefficient of Determination |
| MAE | Mean Absolute Error |
| MSE | Mean Squared Error |
| RF | Random forest |
| RFR | Random forest regressor |

## Funding source

The current work has been supported by the Department of Biotechnology (DBT) grant BT/PR40158/BTIS/137/24/2021.

## Conflict of interest

The authors declare no competing financial or non-financial interests.

## Authors’ contributions

ASR collected and processed the datasets. ASR and NK implemented the algorithms and developed the prediction models. GPSR analyzed the results. ASR and SC created the back end of the web server, and SC and NK created the front-end user interface. NKM did the benchmarking. ASR and GPSR penned the manuscript. GPSR conceived and coordinated the project. All authors have read and approved the final manuscript.

## Acknowledgments

The authors express their gratitude to the University Grants Commission (UGC), Council of Scientific and Industrial Research (CSIR), Department of Science and Technology (DST-INSPIRE), and Department of Biotechnology (DBT) for their generous fellowships and financial support. They also thank the Department of Computational Biology, IIITD, New Delhi, for the excellent infrastructure and facilities. The authors would like to acknowledge the Department of Biotechnology (DBT) for the infrastructure grant awarded to the institute. Furthermore, they would like to acknowledge BioRender.com for creating the figures utilized in this work.

## Data Availability Statement

The datasets generated for this study can be accessed on the ‘HemoPI2.0’ web server at https://webs.iiitd.edu.in/raghava/hemopi2/download.php. The source code for this study is publicly available on GitHub and can be found at https://github.com/raghavagps/HemoPI2.

## Notes

### Competing Interest Statement

The authors have declared no competing interest.

