## Supplementary Data for "Prediction of Hemolytic Peptides and their Hemolytic Concentration (HC_50_)"

**Mailing Address of Authors**

Anand Singh Rathore (ASR):

ORCID ID: https://orcid.org/0009-0004-8907-7174

Nishant Kumar (NK):

ORCID ID: https://orcid.org/0000-0001-7781-9602

Shubham Choudhury (SC):

ORCID ID: https://orcid.org/0000-0002-4509-4683

Naman Kumar Mehta (NKM):

ORCID ID: https://orcid.org/0009-0009-0244-2826

Gajendra P. S. Raghava (GPSR):

ORCID ID: https://orcid.org/0000-0002-8902-2876

*Corresponding Author

Prof. Gajendra P. S. Raghava

Head and Professor

Department of Computational Biology

Indraprastha Institute of Information Technology, Delhi

Okhla Industrial Estate, Phase III, (Near Govind Puri Metro Station)

New Delhi, India – 110020

Office: A-302 (R&D Block)

Website: http://webs.iiitd.edu.in/raghava/

**Author's Biography**

1. Anand Singh Rathore is pursuing a Ph.D. in Computational Biology at the Department of Computational Biology, Indraprastha Institute of Information Technology, New Delhi, India.

2. Nishant Kumar is pursuing a Ph.D. in Computational Biology at the Department of Computational Biology, Indraprastha Institute of Information Technology, New Delhi, India.

3. Shubham Choudhury is pursuing a Ph.D. in Computational Biology at the Department of Computational Biology, Indraprastha Institute of Information Technology, New Delhi, India.

4. Naman Kumar Mehta is pursuing a Ph.D. in Computational Biology at the Department of Computational Biology, Indraprastha Institute of Information Technology, New Delhi, India.

5. Gajendra P. S. Raghava is currently working as a Professor and Head of the Department of Computational Biology, Indraprastha Institute of Information Technology, New Delhi, India.

#### List of Content

| Table | Content | Page No. |
|---|---|---|
| Supplementary Table S1 | List of hemolytic peptides along with their corresponding hemolytic concentrations measured in µM. | 5 |
| Supplementary Table S2 | The entire set of motifs found in hemolytic and non-hemolytic peptides. | 21 |
| Supplementary Table S3 | A detailed correlation analysis of each feature with the $HC_{50}$ value. | 26 |
| Supplementary Table S4 | The detailed evaluation metrics of classification models used to predict hemolytic peptides. | 33 |
| Supplementary Table S5 | The detailed evaluation metrics of hybrid models. | 40 |
| Supplementary Table S6 | The detailed evaluation metrics of regression models used to predict $HC_{50}$ value of peptides. | 41 |
| List of Abbreviations | - | 49 |

### Supplementary Table S1: List of hemolytic peptides along with their corresponding hemolytic concentrations measured in µM.

**This data can be downloaded from:**

**https://webs.iiitd.edu.in/raghava/hemopi2/download.html**

| Sequence | $HC_{50}$ (µM) | Sequence | $HC_{50}$ (µM) | Sequence | $HC_{50}$ (µM) |
|---|---|---|---|---|---|
| GWLPTFGKILRKAMQLGPKLIQPI | 128 | GLPALISWIKRKRL | 596.7 | LLGMIPVAIKAISALSKL | 54.8 |
| GIMSSLMKKLKAHIAK | 400 | IVPFLLGMVPKLVCLITKKC | 7 | GMASKAGSVAGKIAKFALGAL | 500 |
| GFKDLLKGAAKALVKTVLK | 800 | KLKLKLKLKLKLKLKLKLKLKLKLKL | 0.3 | VRRFPFFFPFLRR | 552.5 |
| INWSKIFEKVKNLV | 101 | ILPILSLIGGLL | 180 | FLGALWNVFKSVF | 6.3 |
| KRLFKKLKFSLRKY | 20.1 | CVKVRVKVGSGVKVRVKVC | 590 | FSGGNCRGFRRRCFCTK | 133.7 |
| GIGKFLHTLKTFGKKWVGEIMNS | 35 | SLWENFKNAGKKFILNILDKIR | 128 | FLPAIAGILSQLF | 20 |
| VNWKKVLGKIIKVAK | 200 | GLFDVIKKVLKKIGGL | 22 | AKRHHGYKRKFHAKRHHGYKRKFH | 122.5 |
| FIHHIIGGLFSAGKAIHRLIRRRRR | 19 | FLQHIIGALSHFF | 50 | GLLSVLKGVLKTTGKHIFKNVGGSLLDQAKC KISGQC | 80 |
| YKAWRWAWRWK | 330.3 | FIHHIIGGLFSAGKAEHRLIRRRRR | 50.1 | GIMDTLKNLAKTAGKGALQSLLNKASCKLS GQC | 100 |
| ILCWKWCWWPWRR | 36.4 | GLMDTIKGVAKTVAASWLDKLKCKITG C | 101 | FAKKLAKLAKKLAKAL | 565.9 |
| YKLLKLLLPKLKGLLFKL | 11.1 | FLPLIGRVLSGIL | 120 | YKQCHKKGGKKGSG | 300 |
| GLLSGILGAGKHIVCGLSGLC | 295 | ILPILGNLLNGLL | 300 | GKWMSLLKHIWK | 87 |
| RFCVYAYVRVRGVLVRYRRCF | 60.5 | HIFKKVKTYWKKLFRILGRF | 15.9 | ILCWKWPWWCWRR | 64.3 |
| VSWKKSLGKIIKVVK | 30 | FIGTLIPLALGALTKLFK | 10 | GLLKLIKTLL | 11.4 |
| LKLKLKLKLK | 204.8 | MWSKILGHLIR | 200 | GMASLLAKVLPHVVKLIK | 105 |
| FAKLLAKLAKKL | 164.4 | IPCGESCVWIPCISGMFGCSCKDKVCYS | 13.3 | FLGAIAGVAAKFLPKVFCFITKKC | 8 |
| FLPVIAGLAAKVLPKLFCAITKKC | 5 | ILGKLLKTAAKLLSNL | 38 | KKVLKAAA | 101 |
| SMLSVLKNLGKVGLGFVKCKINKQC | 120 | FLPIVAKLLSGLLGRKKRRQRRR | 44.65 | ALWKTLLKKVLKA | 56.2 |
| RGLRRLGKKIAHGVKKYGPTVLRIIRIA G | 60.5 | SLFSIFKTAAKFVGKNLLKQAGKAGLE TLACKAKNEC | 75 | LLGMIPLAISAISALSKL | 40 |
| KFAKKFAKKFAKKFAKKFAKKFAK | 87.4 | FAKLFAKLAKKFAL | 616.9 | GRFKRFRKKFKKLFKKLS | 200 |
| LKIPGFVRDTLKKVAKEIFSAVTGAVTQS | 256 | RLIKRLKTFVRKTWKWVGHF | 94.2 | KRFKKFFMKLKKSVKKRVMKFFKKPMVIGV TFPF | 50 |
| GVIKAAKKVVKVLKNLF | 136.2 | GLPVCGETCTLGTCYTQGCTCSWPICK RN | 55.3 | AQWFAIQHISLNPPRSTIAMRAINNYRWR | 10.3 |
| SAVGRHLRRFLLRKHRKH | 452.3 7 | GWKKWFTKGERLSQRHFA | 113.3 | FAKKALKALKKL | 722.5 |
| ILGKLLSTAAGLLSKL | 50 | ILPFKFPFFFPFRR | 115.1 | HFLLTKVNLAKKIL | 500 |
| GIMSLFKGVLKTAGKHVAGSLVDQLKC KITGGC | 100 | IFWLFRGKADVAL | 64 | LLLFLLKKRKKRKY | 29.1 |
| INLKILARLAKKIL | 124.6 | KFKKFKKFKKFKKFKKFK | 205.3 | DIFGAIWPLALGALKNLIK | 5 |

| | | | | | |
|---|---|---|---|---|---|
| IKKILSKIKKLLK | 500 | HRIRQLKTTIKKFWEIWPKI | 200 | ALWKTMLKKLGT | 101 |
| KKKKKKKKKGGGLLALLALLA | 25 | KLKKLLKKWLKLLKKLLK | 14 | RIRWILRYWRWS | 100 |
| IILLLLKKFWKKW | 25 | KIAKGALKALKIAKVALKAL | 29.7 | AIGSILGALAKGLPTLISWIKNR | 8 |
| FAKKLAKKAKLAKKL | 584.4 | KLWKKIEKLIKKLLTSIR | 235.9 | GLLKKLLKKLLKKL | 30 |
| GLMDTVKNAAKNLAGQLLDRLKCKIT GC | 135 | KIAGKIAKIAGKIA | 181.9 | AGLQFPVGRVHRLLRK | 216.8 |
| FLPLVTMLLGKLF | 30 | AKRHHGYKRKFHAKRHHGYKRKFHA KRHHGYKRKFH | 123.1 | GKLIKKFGRKAISYAVKKARGKH | 36 |
| WWPWRR | 197.6 | GLLSLSLLGKLL | 50 | ALWTTMLKKLGKMALHAGKAALGAAADTI | 101 |
| RRWLWRLRWLWR | 17.8 | GIPCAESCVWIPCTITALMGCSCKNNVC YNN | 25 | LFNNYITAALKLLEKLYKV | 64 |
| LKKLLKKLLKKLLKKLLKKL | 3.7 | FLPAIAGVAAKFLPKIFCAITKKC | 85 | FPFLLSLIPSAISAIKRL | 104.5 |
| DHYICAKKGGTCNFSPCPLFNRIEGTCY SGKAKCCIR | 31.4 | GLLSGILGAGKNIVCGLSGLC | 295 | ALWKDLLKNVGKAAGKAVLNKVTDMVNQ | 114.7 |
| WRRRRRRRR | 128 | IKPFKKPFKPFRR | 116.7 | RPFTRAQWFAIQHISPRTIAMRAINNYRWR | 178.5 |
| CGKKWGWWKCKL | 548.4 | VRRFPWWAPFLRR | 65 | KSCFRVCYRGICYRRCRG | 32 |
| NVWKKILGKIIKVVK | 200 | RRRWWWWA | 200 | GWKKWLRKGAKHLGQAAIKGLAS | 102.3 |
| GLGSVLGKALKIGANLL | 150 | FLPLLLAGLPKFLCLVFKKC | 29.1 | FAKKLAKKLKKLAKKLIGAVLKV | 9.9 |
| RLYRRLYRRLYRRLYR | 338 | RALRKALKAWRKLAKKLQ | 300 | AAHHIIGGLFSAGKAIHRLIRRRRR | 26.2 |
| RIRFPWPWRWPWWPRFRG | 150 | GLWSKIKKAAKTAGKAALGFVNKMV | 107 | ILPCKWCWWPWRR | 49.4 |
| KLIHRLKTVFKKVWHFLGHL | 40.7 | INQKKIASIGKEV | 100 | GMAKLLAKVLPHVVKLIK | 150 |
| IILLLLKKFLKKW | 12.5 | AGTKEWLNKAKDFIKEKGLGMLRAAA NAALN | 150 | KWMKLLKKILK | 400 |
| GLFDVIKKLLKKIKGL | 6 | AVGIGALFLGFLGAAGSTMGARS | 40 | FFRLLFHGVHHVGKIKPRA | 112.1 |
| GYCFTACYRRNGVRICYRRCN | 40 | GIGHFLHKVKSFGKSWIGEIMNS | 82 | GIGKFIHSAKRFGRAWVGEIMNS | 303 |
| FAKLLAKFLKKAL | 659.6 | KALAKALAKLWKALAKAA | 10 | LGLFKKLLRLIKKGFKK | 98.6 |
| GLLSRIKTLL | 500 | KFLGTLVNLAKKIL | 500 | GVVKRIKTVV | 500 |
| RFARRFARRFARRFARRFARRFAR | 55 | LPFFLLSLIPSAISAIKKI | 9.1 | KKTWWKTWWTKWSQPKKKRKV | 101 |
| GEFLKCGESCVQGECYTPGCSCDWPICK KN | 101 | GIWDTIKSMGKVFAGLILQNL | 18 | GKWMHLLKHILK | 109 |
| NIWKKIFEKVKNLV | 101 | GLFGKILGVGKKVLCGLSGMC | 40 | ASVVKWLTKWVAKLLK | 64 |
| GWRKWIKKATHVGKHIGKAALDAYI | 89.1 | HFLKTLVNLAKKIL | 42 | FLPILAGLAANILPKVFCSITKKC | 7 |
| MPKEKVFLKIEKMGRNIRN | 424.1 | LLKKLLKKLLKKLLKK | 19 | AFGMALKLLKKVL | 241 |
| GEILCNLCTGLINTLENLLTTKRKRQQ | 10 | GILKTIKSIASKLKRKAK | 923 | GIWKTIKSMGKVFAGKIKQNL | 200 |
| FPLTCPTKWWKG | 335.8 | GLSSLGKLL | 300 | IGDLVKWIIDTVNKFTKK | 30 |
| LRLRLRLRLRLRLRLRLR | 205 | FPVTWPTKWWEG | 320.7 | GILSSLWKKLKKWIAK | 57 |
| KIAKGALKALKIAKGALKAL | 99 | KFAKKFKWFAKAAFKFFKK | 80.5 | VNWKKILPKIIKVAK | 200 |
| FLQKIIGALGKLF | 66 | GKLGPLLKIAAKVGSNLL | 800 | KWCFRVCYRGICYRKCR | 85.5 |
| LKKLLKKL | 1000 | ASVVKKLTKGVAKLLK | 128 | GLLDLLKGAGKGLLTHLASQI | 116.7 |
| GKMKEYFKKFGASFKRRFANLKKRL | 36 | GVVVRVPRVVVRWVRR | 128 | GIMRVFKGVLKTAGKSVAKNVAGSFLDRLKC KISGGC | 52 |
| RIRIRWIIR | 191.3 | ALWKTLLKKV | 50 | GLWDTIKQAGKKFFLNVLDKIRCKVAGGCRT | 18.5 |
| GKWSKILGHLIR | 200 | LALKSGGWLRLFGLKDKKH | 92 | KRRVRWIIW | 186.8 |
| GLPALILWIKRKRQQ | 336 | GFGSFLGSLFKTGLKIIPKLLPSIQQ | 30 | FAKLLAKLAKKIL | 674.8 |
| KRLFKELKFSLRKY | 34.1 | FLSLIPHIVSGVASLAKHF | 25 | GWFDILKHVFRLLK | 4.7 |
| GLLSVFKGVLKTAGKNVAKNVAGSLLD QLKCKISGGC | 73 | ALHKTMLKKLGTMAL | 101 | KALKKLLAKWLAAAKALL | 2 |
| SLFGTFAKMALKGASKLIPHLLPSRQQ | 64 | LCYCRRRFCFCV | 51.1 | HFLGTKVNLAKKIL | 500 |
| KKAKKKAKKKAKKKAKKKAKKKAKK KAK | 156.1 | IIKKIIKKIIKKI | 256 | GIKDLLKGAAKALVKTVLK | 800 |

| | | | | | |
|---|---|---|---|---|---|
| ILPIIGKILSTIFGK | 24 | GILDSFKGVAKGVAKDLAGKLLDKLKCKITGC | 101 | FWGALAKGALKLIPSLVSSFT | 6.4 |
| MPKWKVFKKIEKVGRNIRNGIVKAGPAIAVLGEAKALG | 244.8 | GKIIKLKASLKLL | 344.8 | FLPIVGRLISGIL | 101 |
| RFLKHLHTYYERAWHVIGHL | 200 | GVPCGESCVFIPCITGVIGCSCSSNVCYLN | 9.3 | QVVKKIRTWYHKAWHVLGKV | 200 |
| FAKKLAKLAKKLAKLAL | 210.5 | KFFRKLKKSVKKRAKEFFKKPRVIGVSIPF | 87.4 | LNWGAILKHIIK | 119 |
| FLQHIIGALTHIF | 100 | WCRRYRVLVRGVLVRYRRCW | 66 | WWRWRW | 535.9 |
| GIWSSLLKKLKKIIAK | 165 | FIKRIARLLRKIF | 53.1 | GKLFKKILKIL | 307.9 |
| GLMSVTKGVLKTAGKHIFKNVGGSLLDQAKCKISGQC | 125 | GIIKKIIKKIIKK | 512 | FWGALAKGALKLIVGSLFSSFSKKD | 8.8 |
| GLPVCGETCFGGTCNTPGCICDPWPVCTRN | 6.6 | ILPWKWPCWPCRR | 21.2 | RWCVYAYVRVRGVLVRYRRCW | 66.3 |
| FLQGIWDTVGKWL | 37.3 | VNWKKILGKIIKVVK | 200 | FAKKLAKKLKKLAKKLAKLALALKALALKAL | 17.8 |
| KWKLFKKIGAVLKVL | 18.7 | GFKRIVQRIKDFLRNLV | 160 | GMWKKILGKLIR | 200 |
| KFFKKFFKKFFKKFFKKFFKKFFKKFFK | 3.1 | GFSSIFRGVAKFASKGLGKDLAKLGVDLVA | 200 | KWKVFKKAEKMIRNIRNKIVK | 250 |
| KQKLAKLKAKLQKLKQKLAKL | 600 | GYPICGESCVGGICNIPGCSCSWPVCTTN | 225.9 | FAKLLFKALKKAL | 659.6 |
| KWKKLLKKLLPLLKKLLK | 50 | FPVTWRWWTWWKG | 268.7 | ALYKTMLKKLGTMAL | 101 |
| WWWLRRRW | 150 | KLKLKLKL | 253.7 | GFRKIVQKIRDFLRNLV | 80 |
| IKKIIKKI | 1000 | FAKLLAKLAKAKA | 715.3 | GWGSFFKKAAHVGKHVGKAALTHYL | 93.6 |
| GIGAVLGVLTTGLPALISWIKRKRQQ | 1.2 | ISLNPPRSTIAMRAINNYRWRSKNQNTFLR | 10.5 | ALWMTLLKKVLKAAAK | 23.1 |
| GIMSSLMKKLAKIIKK | 400 | GIGAVLKVLTTGLPALISWIKRKRQQ | 0.6 | LLKKLLKKM | 876.4 |
| FLSTLLNVASNVVPTLICKITKKC | 19 | LIGLVSKGTSVLVKTVSKKVLKQ | 15 | CLLKKLLKKLLKKC | 48.3 |
| LLKKLLKKLLKK | 335.3 | FAKKLAKALL | 886.5 | INWKKIFESVKNLV | 101 |
| KKFPWWWPFKK | 200 | KQIKIWFQNKKMKWKK | 128 | KGIVGMLGKLF | 618 |
| FLKGIVGMLGKLL | 45 | GFLSILKKVL | 101 | FLPIIGQLLSGLL | 300 |
| SLLSLIRKLIT | 1000 | LNLKALLAVAKKIL | 40.11 | FPVGRVHRLLRK | 271 |
| GLWNSIKIAGKKLFVNVLDKIR | 254.11 | LLIILRRRWRRQARARSR | 94 | IKKWLSKIKKLLK | 500 |
| KNWKKILKKIIKVVK | 200 | GFLGPLLKLGLKGVAKVIPHLIPSRQQ | 50 | HFLLTLVNLAKKIK | 500 |
| PFKIDIHLGGY | 50.4 | FDIMGLIKKVAGAL | 200 | FFLLSLIPSAISAIKKI | 33.3 |
| FLPVIAGAANFLPKLFCAISKKC | 14 | FFHHIFRGIVHVGKKIHRLVTG | 46 | GFMKYIGPLIPHAVKAIKKLI | 25 |
| NLLNKALGTVNGLLGRS | 256 | KVLKAAA | 101 | QYLRRVRTWLRRAWHILGKV | 20.8 |
| HFLLTLVKLAKKIL | 38 | QSHLSLCRWCCNCCHNKGCGFCCKF | 34.71 | GIGRFLHSARRFGRAFVGEIMNS | 249.4 |
| AIPCGESCVWIPCISTVIGCSCSNKVCYR | 25.5 | FLPVILPVIGKLLSGIL | 50 | FALALKALKKAL | 457.3 |
| RWGRWLRKIRRWRPKC | 14.1 | AQWFAIQHISLNPPRSTIAMRAINNYRWRSKNQNTFLR | 10.5 | SMLAVLKNLGKVGLGFVACKINKQC | 200 |
| GLFDVIKKVASVIGGL | 19.8 | KFGKIVGKVLKQLKKVSAVAKVAMKKG | 275.5 | GWKKWFNRAKKVGKTVGGLAVDHYL | 88.8 |
| WWWRRRIW | 200 | FAKLLAKLAKLKL | 404.9 | FLPGILKVAANVVPGVI | 512 |
| GAFGDLLKGVAKEAGMKLLNMAQCKLSGKC | 520 | FKAFKAFKAFKAFKAFKAFKAFKAFKAFKA | 143.8 | INSLKLGKKILGAL | 101 |
| GIWKKWIKKWLKVLKNLF | 23.5 | GSKRWRKFEKRVKKIFEETKEALPVVQGVVGVATAVGRK | 112.7 | VRRFPYYYPFLRR | 538.2 |
| FIHHIIGGLFSVGKHIHSLIHRRRR | 6.3 | IFGTILGFLKGL | 64 | KRIVQRIKDWLR | 800 |
| HFLGTLVNLAKKKL | 500 | GMWSKILGKLIR | 143 | KRIRKRIKKWLR | 800 |
| FFRNLWKGAKAAFRAGHAAWRA | 34.6 | WRWLRRIW | 100 | GLGKFLHSAKRFGKAFVGEAMNS | 1000 |
| RFARRFARRFARRFARRFARRFARRFAR | 5.4 | GLFGKLIKKKGRKAISYAVKKARGKH | 36 | IDWKKVDWKKVSKKTSKVMLKASKFL | 20 |
| FLPLIASVAANLVPKIFCKITKKC | 75 | LLKKLLKK | 495.5 | IWRIFRRIF | 101.4 |
| RFGRFLRKIRRFRPK | 48.1 | GLKKIFKKGLGSLVKGIAAHVAS | 400 | KYIRHLKTWFKKVFKLIGEV | 19.8 |
| LLGDEFRKSKEKIGKEFKRIVQRIKDFLRNLVPRTES | 43.5 | RWCRRYRVRVRGVLVYAYVCW | 80 | KWKVFKKIEKMGRNIRNGIVKAGPAIAVLGEAKAL | 250 |

| | | | | | |
|---|---|---|---|---|---|
| RGGRLCYCRRRFCICV | 40.8 | VNFKKLLGKLLKVVK | 200 | RRRRWWWL | 200 |
| GLNALKKVFQGIHKAIKLINNHVQ | 240 | LLPALISWIKRKRQQ | 533.3 | FKRFKGSVKHGKGHLVHHIGVAL | 400 |
| KIFKKFKTIIKKVWRIFGRF | 15.9 | LKIPGFVKDTLKKVAKGIFSAVAGAMTP S | 300 | GLLKRIKKLLKKIKKLL | 9 |
| GSKRWRKFEKRVKKIFEETKEALPVIQG VATIVGAVGR | 74.5 | FLGAIAAALPHVINAVTNAL | 120 | ALWKTLLKKVLKAAAKAALKAVLVGANA | 0.5 |
| RWCFRVCYRGICYRKCRG | 64.2 | SMLSVLKNLGKVGLGFVA | 200 | CLNLKALLAVAKKILC | 9.19 |
| GGLRSLGRKILRAWKKYGPIIVPIIRIG | 19.2 | FFGKVLKLIRKIF | 19.6 | ILGKIWKGIKSLF | 60.3 |
| FIFHIIKGLFHAGKMI | 210.6 | KIAKIAGKIAKIAGKIAKIAGKIA | 2.9 | AIPFIFIWRLLRKG | 15 |
| FWSTLLSIGKSLL | 10.3 | LFGMALKLLKKVL | 53 | GLPALISWIKRKRGG | 596.3 |
| GWYEIIKKIYKWLK | 5.1 | GLKALKKVFKGIHKAIKLINKHVQ | 180 | VNWKKVLKKIIKVAK | 200 |
| NLLGSLLKTGLKVGSNLL | 256 | FLGFLHHLF | 175.9 | GFMSKVANFAKKFAKGGVNAIMNQK | 64 |
| VKKFAWWWPFLKK | 65 | GLFTLIKGAAKLIGKTVAKEAGKTGLEL MACKITNQC | 100 | KIFKKFKDFLKKIFQYLGKV | 14.9 |
| GFGKALKLLKKVL | 800 | GYCFTACYRRNGARICYRRCN | 100 | FAKLAKKLL | 94.6 |
| GLLSALKALGKAL | 50 | LKKLLKKLLKKLLKKLLKKLLKKLLKK LLKKL | 2.1 | VNSKKISGKSIKVSK | 100 |
| FLPFLKSILGKIL | 50 | FLPMLAGLAASMVPKLVCLITKKC | 5 | LKLKLKLKLKLKLKLKLKLKLKLK | 171.8 |
| AKKVFKRLPKLFSKIWNWK | 800 | IIKKFLHSIWKFGKAFVGEIMNI | 16 | GLPICGETCVGGTCNTPGCSCSWPVCTRN | 6.96 |
| LPKVMAHMK | 101 | VNWKKILGKIIKVAK | 200 | KKIASIGKEVLKAL | 101 |
| VNWKKILPKIIKVVK | 200 | ILGKLLSTAAKLLSNL | 24 | GIGKFLHSAKKFGKAWVGEIMNS | 509 |
| GMWSKILGHLIK | 132 | FLPILAGLAAKIVPKLFCLATKKC | 5 | INWLKLGKKLLSAL | 44 |
| INWLRLGRRILGAL | 25 | GFMKYIGPLIPHAVKAISKLI | 120 | RPFIRAQWFAIQHISPRTIAMRAINNYRWR | 150 |
| GWRTLLKKAEVKTVGKLALKHYL | 95.6 | ILGTILGLLKGL | 64 | GLNALKKVFQGIHEAFKLINNHVQ | 250 |
| GKWKKILGKLIR | 200 | GKWMSKLKHILK | 200 | LKWLKWG | 150 |
| FLPIVTNLLSGLL | 116 | ALWMTLLKKVLKAAAKAALN | 5 | IRAWAKVLKGVGKILKGVGRW | 134.8 |
| FLGSLFSIGSKLLPGVIKLFQRKKQ | 28 | ALWKTLLKKVLKAAAK | 57.2 | FCYCRRRFCVCVGR | 45.3 |
| CAKAKAKAGSGAKAKAKAC | 1000 | RGGRLCYCRPRFCVCVGR | 38.1 | FIHHIIGGLFSAGKAIHRLIRRR | 7 |
| FAKKFAKKFAKKFAKKFAKKFAKK | 174.7 | IKKIIKKIIK | 1000 | GAGCNIFGGNDYRCHRHCKSIRGYKGGYCK LGGICKCY | 400 |
| FLAGLIGGLAKMLGK | 40 | VNRKKILGKSIKVVK | 100 | RGGRLCYCRRRFCVCF | 40.1 |
| WRRRYRRWRRRRRWRRRPRR | 200 | GLMSLFRGVLKTAGKHIFKNVGGSLLD QAKCKITGEC | 65 | GFFGKMKEYFKKFGASFKRRFANLKKRL | 12 |
| GAGALAKFLAKKVAKTVAKQAAKQGA KYVVNKQME | 40 | FWGAAAKMLGKALPGLISMFQKN | 48.3 | LRRLWLRANRLLRRLWLRANRL | 66 |
| GFGSLFKFLAKKVAKTVAKQAAKQGAK YVANKHME | 14.5 | LRIIRRILRRLI | 125 | RRWCVYAYVRVRGVRVYAYVCW | 80 |
| SLSRFLSFLKIVYPPAF | 99.5 | SFLTTVKKLVTNLAAL | 178 | GILSSLWKKPKKIIAK | 400 |
| CVKVQVKVGSGVKVQVKVC | 1000 | FFFLRRIF | 200 | RFRRLRKKTRKRLKKI | 256 |
| LCYCRRRFCVCV | 52.7 | GVSKILHSAGKFGKAFLGEIMKS | 200 | GLPVCGETCVGGTCNTPGCTCSWPVCTRN | 29.1 |
| ALWKTMLKKLGTMA | 101 | FALALKALKKLLKKLKKLAKKAL | 3.8 | KRRLILRILRLAIRALVKKR | 20 |
| WLRRIKAWLRRIKA | 256 | FVKLKKIANIINSIFKK | 1000 | KLKLLLLLKLK | 300 |
| ISQSDAILSAIWSGIKSLF | 129 | KLKLLLLKLK | 146 | FAKKFAKKFKKFAKKFAKFAFAF | 18.8 |
| KRIRKRIKKWKR | 800 | GIWDTIKSMGKVFAGKIKQNL | 200 | RQYMRQIEQALRYGYRISRR | 200 |
| GMWSKILKHLIR | 45 | KLKLLLKLGLKLL | 50 | KKYRYHLKPFCKK | 128 |
| KWCFRVCSRGSCYRRCRG | 64 | GFFGKRKEYFKKFGASFKRRFANLKKR L | 36 | FLSLIPKIISAISALIKHF | 1.074 |
| IKLSKETKDNLKKVLKGAIKGAIAVAKM V | 203.9 | GLLKRIATLL | 117.5 | GLPALISWIKRKRQQ | 549.7 |
| KWKLWKKIEKWGQGIGAVLKWLTTWL | 93.1 | FALAKLAKKAKAKLKKALKAL | 39.5 | VRRFPYWWPFLRR | 65 |
| GFRDVLKGAAKAFVKTVAGHIAN | 200 | IKKIVSKIKKLLK | 1000 | GRKKRRQRRRGALWKSLLKNVGKA | 512 |

| | | | | | |
|---|---|---|---|---|---|
| ILKWKWKWWKWRR | 395.1 | VRRFPWYWPFLRR | 65 | IAGKIAKIAGKIAKIAGKIA | 10.2 |
| FKIKASKVLDKFGKIVGKVLKQLKKVS AVAKV | 12.7 | GVIIKTLKGAAKTVAAKLL | 1000 | NIWKKIASIAKEVLKAL | 32 |
| GVFLDALKKFAKGGMNAVLNPK | 150 | GLLSLSLGKLL | 300 | FLGAILKIGHALAKTVLPMVTNAFKPKQ | 48.9 |
| ILKWKWKWWPWRR | 280.8 | WRRFWHR | 500 | FLSLIPHIVSGVASIAKHF | 39 |
| GKLGPLLKIAAKVGSKLL | 800 | IWRIFRRIFRIF | 19.6 | KNLRRIIRKGIHIIKKYF | 50 |
| FLKWALKWAAK | 64.03 | GTGLPMSERRKIMLMMR | 200 | VCKRWKKWKRKWKKWCV | 133.3 |
| GLLDTLKGAAKNVVGSLASKVMEKL | 400 | KLYKKFRHILKKVWHYVGKI | 28.8 | GIAKFGKAAAHFGKKWVGELMNS | 700 |
| FFGWLIRGAIHAGKAIHGLIHRRRH | 1 | SKVLRHLRRFLHRAHRKL | 460.7 8 | GLLKKIKKLL | 500 |
| QLLKKVRTWYRKAWHLYGKV | 200 | LKLGTLVNLAKKIL | 500 | GIWKTIKSMGKVFAGKILQNL | 95 |
| GLLSLLSLLGKLA | 3 | GLFDVIRKVASVIGGL | 19.5 | KRFKKFFKKLKNSVKKRAKKFFKKPKVIGVT FPF | 0.37 |
| GEFAQFEQEAQSLEQELSQLEQQLESL | 250 | FLGALWRVAKSVF | 12.5 | KWAVRIIRKFIKGFIS | 128 |
| GECIWDAIFHGAKHFLHRLVNP | 19.3 | RLYRRLYRK | 1000 | FFPIVGKLLSGLF | 150 |
| FPVTWKWWKWWKG | 500 | RKKRRQRRRLNLKALLAVAKKIL | 77.94 | FIHHIIGGAASAGKAIHRLIRRRRR | 35 |
| FAKGIAGMAGKLF | 200 | GRFKRFRKKFKKLFKKLSPVIPLLHLG | 250 | GLLSSLLLLGKLL | 20 |
| GFISTVKNLATNVAGTVIDTIKCKVTGG C | 300 | FAKKLAKKLAKLAL | 446.4 | IAGKIAKIAGKIAKIAGKIAKIAGKIA | 0.4 |
| ALWKDLLKNVGIAAGKAALNKVTDMV NQ | 436.7 | VKKFPWWWPFLKK | 551 | IPCGESCVWIPCITAIAGCSCKNKVCYT | 11.6 |
| RFRRLRKKIRKRLKKI | 256 | LWKIGKKIWRVLWNWR | 7.3 | LLKKLLKKLLKL | 338.7 |
| FWGFLGKLAMKAVPSLIGGNKK | 103.8 | LKMLGMLFHNIRNILKTV | 128 | RSKKWRKIEKRVKKIFEKTKEALPVIQGVATI VGAVGR | 43.2 |
| FKRIVQRIKDFLR | 256 | FAKLLAKAFKKAL | 678.4 | FLGALWNVAKSYF | 12.5 |
| GIKEFAHSLGKFGKAFVGGILNQ | 200 | GLFDIWKWWRWRR | 129.5 | LFGFLIKLIPSLFGALSNIGRNRNQ | 1.9 |
| ILLKKLLKKI | 809.7 | INWKSIFEKVKNLV | 101 | WKLFKKILKYL | 42 |
| VNWKKILG | 200 | KLLGPLLKIAAKVGSNLL | 400 | GLPTCGETCTLGTCYVPDCSCSWPICMKN | 13.1 |
| GKWMSLLKQILK | 110 | CWKWKWKWGSGWKWKWKWC | 45 | WKKIASIGKEVLKAL | 101 |
| RRWQWR | 101 | GLLSVLKGVLKTAGKHIFKNVGGSLLD QAKCKISGEC | 80 | AKVVKKLTKGVAGLLK | 128 |
| KRIVQRIKKWLR | 800 | RQIRIWFQNRRMRWRR | 128 | LWKIWKKIWRVWKNWR | 6.9 |
| FLPAVLKVAAHILPTAICAISRRC | 200 | FAKKLAKKLAKAAL | 655.3 | GLFDIWKKLRWRR | 139 |
| GRFKRFRKKLKRLWHKVGPFVGPILHY | 100 | LKLKLKLKL | 129.2 | SAVLRHLRRFLLRKHRKH | 448.9 7 |
| FLPKWALKWAAK | 328.3 | GLKALKKVFKGIHKAIKLINNHVQ | 330 | KFAKKFAK | 258.8 |
| CKGKGAKCSRLMYDCCTGSCRSGKC | 1000 | GIGKFLHSAKKFGKAFVGEIMNS | 17 | VRRFPWWWAFLRR | 186.2 |
| LRWLRWG | 150 | KWRRWVRWI | 289 | FKIKPGKVLDKFGKIVGKVLKQLKKVSAVAK V | 90.5 |
| FFFIRRIARLLRRIF | 5.6 | FAKKLAKKLAKAL | 687.3 | KWCSRVCYRGICSRRCRG | 32 |
| DSHAKRHHGYKRKFHEKHHSHRGY | 49.1 | IKKIWSKIKKLLK | 500 | GIRKWFKKAAHVGKEVGKVALNACL | 94.1 |
| IKIPPIVKDTLKKVAKGVLSTVADALSKS | 512 | GLLTRIKTLL | 500 | GLGSLLGKALKFGLKAAGKFMGGEPQQ | 200 |
| DAACAAKCLWR | 162.2 | GWLDVAKKKGKAAFNVAKNFI | 270 | YRMWRWAWRWR | 230.5 |
| IDWKKIFEKVKDLV | 101 | GLVGTLLGHIGKAILS | 166.3 | KWCFRVCYRGICYRKCRG | 128 |
| FFPLIFGALSSILPKIL | 31 | GLWKKIKNVAKAAGKAAKGAL | 834.6 | GMASKAGSVLGKVAKVALKAAL | 500 |
| QVFKKFRPFYRRPWELFGKL | 200 | IWLTALKFLGKHAAKHLAKQQLSKL | 125 | KFKKVIWKSFL | 50 |
| NFLDTLINLAKKFI | 80 | GMWKKILGHLIR | 93 | SILDKIKNVALGVARGAGTGILKALLCKLDKS C | 480 |
| WWWLRRIR | 200 | RWQWRWQWR | 101 | RLYRRLYRRLYRRLYRKKK | 1000 |
| FLPKLLAGLPSFLCLVFKKC | 31 | GLGSLLGKAFKFGLKTVGKMMGGAPR EE | 105 | GLLDLLKLLLKAAG | 63.2 |
| GKGALKKFLAKKVAKTVAKQAAKQGA KYVVNKQME | 40 | VRRFPWYYPFLRR | 65 | VRRFAWWWAFLRR | 32 |

| | | | | | |
|---|---|---|---|---|---|
| LLKKLLKLLLKL | 9.3 | FLREFHKWIERVVGWLGKVF | 128 | GLPLLISWIKRKRQQ | 537.3 |
| LNWKAILKHIIK | 110 | GILSSIKGVAKGVAKNVAAQLLDTLKC KITGC | 101 | IKRQYKRFFKLFKWFLKK | 25 |
| GLNALKKVFQGIHEAIKLINNHVQ | 360 | FFHHIFRPIVHVGKTIHRLVTG | 200.3 | GFSSIFRGVAKFASKGLGKDLAKLGVDLVAS KISKQS | 200 |
| VRRFPWAAPFLRR | 65 | FWGKLWEGVKNAI | 101 | GLKEIFKAGLGSLVKGIAAHVAS | 145 |
| FFPGIIKVAGAILPTAICAITKRC | 100 | HFLGTLVKLAKKIL | 38 | ALWKTLLKKVLKAAA | 19.7 |
| SILPTIVSFLSKVF | 12.5 | CVKVKVKVGSGVKVKVKVC | 880 | GLVGTLLGHIGKAILG | 404.8 |
| ILKKLLSTAAGLLSNL | 105 | SFLTTVKKLVTNLAALAGTVIDTIKCKV TGGCRT | 16.11 | KWKLFKKALKKLKKALKKAL | 32 |
| ASVVKKLWKGVWKLLK | 128 | FAKKLKKLAKLAKKL | 513.4 | VCRTGRSRWRDVCRNFMRRYQSR | 300 |
| FWGALAKGALKLIGSLFSSFSKKD | 1.4 | GIWKKWIKKWLNVLKNLF | 24.7 | GLLKWIKTLL | 20 |
| QIIKKIRTWYRKAWHVLGKV | 200 | GLFDIIKKIAESF | 128 | TLLKKVLKAAAKAALNAVLVGANA | 16.5 |
| IKKIWSKIKKWWK | 500 | RFGRFLRKILRFLLK | 12.6 | GKYTCGETCFKGKCYTPGCTCSYPICKKD | 64 |
| ITKQITKQLNRQTKIQSK | 250 | WCYCRRRFCVCVGR | 44.3 | GIGKHVGKALKGLKGLLKGLGES | 800 |
| GIFPIFAKLLGKVIKVASSLISKGRTE | 6.2 | FLKGIKGMLGKLF | 590 | IKLSPETKDNLKKVLKGAIKGAIAVAKMV | 225 |
| GFMKYIGPLIPHAVKAISDLI | 65 | RLYRKVYG | 1000 | FPFLLSLIPSAISALKKL | 30.3 |
| GIPCAESCVWIPCTVTALVGCSCSDKVC YN | 8.7 | FFHHIFRGIVHVGKTIHRLVTG | 11 | YCYCRRRFCVCVGR | 44.9 |
| ILPWKWEWWPWRR | 11 | GKWKSLLKHILK | 200 | ALWKNMLKGIGKLAGKAALGAVKKLVGAES | 50 |
| LLRRLLRR | 446 | WKRIVRRIKRWLR | 256 | GFWGKLWEGVKNAIKKK | 1000 |
| GKWSKILGKLIR | 200 | FLPFLLSALPKVFCFFSKKC | 3 | GLLGKLLKIAAKVGKKLL | 25 |
| GKWMTLLKHILK | 86 | GLLRASSVWGRKYYVDLAGCAKA | 500 | FAKLLAKALKKFL | 461.7 |
| RNFFKRIRRAGKRIRKAIISAAPAVETLA QAQKIIKGGD | 735.4 | IKKILSKIKKLL | 550 | HKWMSLLKHILK | 200 |
| GFGTILKALAKIAGKVVKKLATKPGATY MLKENLK | 2.6 | GVIPCGESCVFIPCISTLLGCSCKNKVCY RN | 550 | GLLKVIKTLL | 53.4 |
| SWLSKTAKKLENSAKKRISEGIAIAQGP R | 31.6 | KLSLLLSLGLKLL | 25 | VLGLALIVGGALLIKKKQAKS | 213.5 |
| GIWDTIKSMGKVFAGAILQNL | 20 | FLKGIVGKLGKLF | 530 | SLLGTVKDLLIGAGKSAAQSVLKGLSCKLSK DC | 300 |
| WKKWWKKWWKKW | 483 | RGGCLCYCRRRFCVCVCR | 798.6 | GFWTTAAEGLKKFAKAGLASILNPK | 105 |
| GDACGETCFTGICFTAGCSCNPWPTCTR N | 51.9 | ALWKTLLKHVGKAAGKAALNAVTDM VNQ | 128 | FLGALWNVLKSVF | 12.5 |
| LNWGAFLKHFFK | 81 | KLLKRIKKLL | 288.7 | GVWSTVLGGLKKFAKGGLEAIVNPK | 128 |
| ASVVNKLTGGVAGLLK | 128 | AARRAARRAARR | 355.4 | RFLKHFKTVYKRYWKVLGRL | 142.9 |
| GFCRCLCRRGVCRCLCTK | 1000 | SMLSVLKNLGKVGLGFVACKINKKC | 80 | VRRFPWWWPFLRR | 200 |
| FAKLLALALKLKL | 238.6 | RKLRRLKRKIAHKVKKY | 101 | FIITGLVRGLTKLF | 124.8 |
| KMWSKILGHLIR | 200 | GLWNTIKEAGKKFAINVLDKIRCGIAGG CKT | 75 | GLSLLLSLLGKLL | 3.1 |
| RFRRLRKKKRKRLKKI | 256 | GLLSALSLLGKLL | 6 | GLPVCGETCVGGTCNTPGCSCSWPVCFRN | 27.06 |
| AKKFAKKFAKKFAKKFAKKFAKKF | 13.6 | GLLKKIKWLL | 152.5 3 | LCYCRRRFCWCV | 49.8 |
| GLPICGETCVGGTCNTPGCFCTWPVCTR N | 11.53 | SLWENFKNAGKKFILNILDK | 128 | RVKRVWKLVIRLVKALYKLYRAIKKK | 1 |
| GKLLSLLSLLGLL | 20 | KRRLIARILRLAARALVKKR | 40 | GFSSIFRGVAKFASKGLGKKLAKLGVKLVAC KISKQC | 11 |
| FLSALWGVAKSLF | 30.7 | FLQHIIGALGKLF | 40 | IFFRRNKKMAVKVAINGFGRIGRLAFRQMF | 1000 |
| SYERKINRHFKTLKKNLKKK | 249.4 | FLPILASLAAKLGPKLFCLVTKKC | 4 | FLPVIASVAAKVLPKVFCFITKKC | 7 |
| GLLKRIKSLL | 500 | DLFKQLQRLFLGILYCLYKIW | 130.9 | RGGRLCYCRRRFCVCL | 40.8 |
| RLYRKVYGRLYRKVYGK | 1000 | INWSSIFEKVKNLV | 101 | GKWMKLLKHILK | 87.4 |
| GLLKWIKKLL | 11.58 | GEYCGESCYLIPCFTPGCYCVSRQCVN KN | 25 | SLWENFKNAGKQFILNILDKIRCRVAGGCRT | 220 |
| RRWQWRLCYCRRRFCVCVG | 100.6 | VGKVLKQLKKVSAVAKVAMKKGAALL K | 356.5 | GAVVDILKGAGKNLLSLALNKLSEKV | 186.9 |
| WCPPMIPLCSRF | 500 | GIFPIFAKLLGKVIKVASSLISKGRTK | 4 | GLPVCGETCFTGSCYTPGCSCNWPVCNRN | 4.29 |

| | | | | | |
|---|---|---|---|---|---|
| GLLSLLSLLGKAL | 4 | GLKALKKVFQGIHEAIKLINNHVQ | 340 | HFLGTLVNKAKKIL | 500 |
| GLFGKLIKKFLRKAISYAVKKARGKH | 3 | IDWKKVDWKKVSKKTCKVMLKACKF L | 20 | ILGKLLSTAAKLLSKL | 16 |
| NFLGTLVNLAKKIL | 64 | KLKLKLKLKLKLKLKLKLKL | 0.4 | INWKKIASIGKEVLKAI | 101 |
| GFKKLLKGAAKALVKTVLF | 24 | FAKKLAKKLKKLAKKLAK | 487.1 | IKLSPETKKNLKKVLKGAIKGAIAVAKMV | 173 |
| RGLRRLGRKIAHGVKKGGPTVLRIIRIA G | 101 | QRIRQLHTWIKKAWHIWPKI | 200 | IKLSKKTKDNLKKVLKGAIKGAIAVAKMV | 136.3 |
| VYWKKILGKIIKVVK | 190 | FLPIIGKLLSGLL | 95 | FLPFVGNLLNGLL | 150 |
| GLLKRIKWLL | 77.5 | GKWMSWLKHILK | 87 | HFLGTLVNLAKKIK | 500 |
| LKKLLKKLLKKLLKKLLKKLLKKL | 3.8 | LLAGLAANFLPKIFCKITRKC | 32.8 | GLMSLFKGVLKTAGKHIFKNVGGSLLDQAK CKITGEC | 55 |
| YKLLKLLLPKLKPLLFKL | 24.9 | FLSTLLNVASKVVPTLFCKITKKC | 37.5 | FLGALLGPLMNLLQ | 105 |
| GFLSILKKVLKKVMAHMK | 27.3 | ILKWKWPWWPWRR | 203.8 | GMWPKILGHLIR | 200 |
| GIPCGESCVWIPCISAAIGCSCKSKVCYR N | 11 | SKHWLWLW | 17.3 | KWCFCVCYRGICYCRCRG | 590 |
| FAKALKALLKALKAL | 3.7 | ILPWKKPWEPWRR | 110.1 | FAKLLAKLAKKAA | 715.3 |
| ILGKLLSWAAGLLSNL | 13 | WLWKKIKNVAKAAGKAAKGAL | 1000 | LVKKLLKLAMGFG | 200 |
| FIHHIIGGLFSAGKAAHRLIRRRRR | 8.5 | NLWAGILKHIIK | 200 | ASVVKKWTKGWAKLLK | 128 |
| LLGSLLKLLPKLL | 49.2 | GLLKRIKNLL | 319.4 | RGGRLCYCRRRFCVCW | 39.4 |
| CLLKKLLKK | 899.3 | RGTCYNRVGLIIRNFSKLKGKKV | 261.1 8 | FLPAIAGLAAKFLPKIFCAITKKC | 16 |
| GIMDSVKGLAKNLAGKLLDSLKCKITG C | 200 | FLPGLIAGIAKML | 73 | ILPYKYPYYPYRR | 110.1 |
| RWWRWWR | 731.9 | AKKVFKRLEKLFSKIWNWK | 397.7 | FALALKALKKALKKLKKALKKAL | 39.5 |
| GLPALISWIKRKR | 639.8 | GFGMALRLLRRVL | 72 | LKLKLKLKLKLKLKLKLKLKLK | 187.3 |
| IWLTALKFLGKNLGKHLAKQQLAKL | 32.35 | GIMSSLMKKLAKIIAK | 88 | FLSGIVGMLGKLF | 25 |
| RWNGRIIKGFYNLVKIWKDLKG | 88.9 | KNWKKILGKIIKVVK | 200 | FKIKPGKVLDKFGKIVGKVLKQLKKVS | 297.7 |
| AGWGSIFKHIFKAGKFIHGAIQAHND | 89 | VKWRWKWKWRWKWKWKV | 121.9 | HFKGTLVNLAKKIL | 500 |
| GLWSKIKNVAAAAGKAALGAL | 106.4 | GIMISLMKKLAAHIAK | 320 | FKAFKAFKAFKAFKAFKAFKAFKAFKA | 159.6 |
| GASCGETCFTGICFTAGCSCNPWPTCTR N | 51.9 | KRFWQLVPLAIKIYRAWKRR | 20 | LNWGAALKHAAK | 200 |
| FLPIIAGAAAKVVQKIFCAISKKC | 16 | GWKDWLNKAKDFIKEKGPEILRAAAN AAIN | 101 | FFRLLFHGVHHGGGYLNAA | 119.7 |
| FFHHIFRGAVHVGKTIHRLVTG | 2.4 | FAKKLAKLAKKALAL | 610.1 | ASVVKWWTKWWAKLLK | 64 |
| GILSSWLKKLKKIIAK | 400 | IIGAIAAALPHVINAIKNTF | 120 | FLGALWNVAKKVF | 25 |
| KNWKAILKHIIK | 200 | GLKKIFKAGLGSLVKGIKAHVAS | 400 | GLLSLASLLGKLL | 4 |
| EKRWRRLIFNYF | 200 | RGLRRLGRKIAHGVKKYGPTVLRIIRTA T | 87.6 | LKLKLKLKLKLKL | 1.2 |
| GALSLLSLLGKLL | 4 | FIHHIIGGLFSVGKHIHSLIHGH | 9.8 | QVLKKVRTWYRKAFHVIGKV | 200 |
| FAKKLAKLAKKL | 722.5 | KKKLAKLKLGAKLKLKGKLGA | 100 | FFHHVFRGIVHVGKTIHRLVTG | 40 |
| KKKFPWWWPFKKK | 200 | IIGGLFSAGKAIHRLIRRRRR | 61.4 | GLWSKIKEVGKEAAKAAAKAAGKAALGAVS EAV | 500 |
| GMASLLAKVLPKVVKLIK | 200 | MAADIISTIGDLVKLIINTVKKFQK | 4.4 | KWKLFKKKTKLFKKFAKKLAKKL | 2.8 |
| ASVVNKLTKGVAKLLK | 128 | VNWKKILGKIKKVVK | 200 | YKLLKLLLPKLKPLLPKL | 58 |
| HLIHRLHTYWHKPWHYLGKL | 200 | FAKLFAKAFKKAL | 663.1 | FLPIVAGLAANFLPKIVCKITKKC | 19 |
| IIPLPLKKFLKKL | 500 | FIHHIIGGLFSAGKARHRLIRRRRR | 33.1 | GLLALISWIKRKRQQ | 65.4 |
| LLIILKKKWKKQAKAKSK | 200 | SKVGRHGRRFGHRAHRKL | 500 | FLSLIPHIVSGVAALANHL | 53.98 |
| ALWMTLLKKVLKAAAKAALDAVLVGA NA | 1.2 | YKLLKWLLLKLKALLEKL | 2.8 | KLKLKLKLKLKLKLKLKLKLKL | 0.4 |
| FLGFLKNLF | 122.7 | LALALALALALALALALALA | 220.4 | RGLRRLGRKIAHGVKKYGPTVLRIIRIA | 86 |
| GFFALIPKIISSPLFKTLLSAVGSALSSSGG QE | 50 | IFKAIWSGIKSLF | 59.25 | LWKIWKKIWRVGWNWR | 21.3 |
| HIIKKIRTWYRKAWHVLGKV | 200 | FIGAIARLLSKIF | 11.6 | GIGAVLKVLTTGLCALISWIKRKRQQ | 2.5 |

| | | | | | |
|---|---|---|---|---|---|
| KFAKKFAKKFAKKFAKKFAKKFAKKFAKKFA K | 74.9 | CRWGRWLRKIRRWRPK | 5.5 | GLKKIFKAGLGSLVKGIAAHVAS | 400 |
| LNLKGLFKKVASLLT | 353.4 | KNLRRIIRKGIHIIKKYG | 101 | FLPLLLAGLPKLLCLFFKKC | 5 |
| RMMRIFWVIKLAR | 32 | FLPVLTGLTPSIVPKLVCLLTKKC | 150 | KRLFKKLLFSLRKY | 23.1 |
| GLWKKIKNVAAAAGKAALGAL | 444.8 | FASGIAGMAGKLF | 101 | LLGSLLSLGLKLL | 13 |
| RFRRLRKKFRKRLKKI | 256 | FWGALAKGALKLIGVSLFSSFSKKD | 9.3 | RPPQFTRAQWFAIQHISLN | 11.2 |
| GVIIDTLKGAAKTVAAELL | 1000 | VNWKKVLGKVVKVVK | 200 | KKFAKKFAKKFAKKFAKKFAKKFA | 174.7 |
| GIGKFIHSAKKFGKLFVGEIMNS | 260 | GIKKFLKSAKKFVKAFK | 50 | KKYRYHLKPFSKK | 128 |
| ALWDTLLKKVLKAAAKAALNAVLVGA NA | 2.3 | GVIKSVLKGVAKTVALGML | 280 | FLQGIIDTVGKWL | 82.3 |
| KLKLKLKLKLK | 185.8 | KLLKLLLKLLLKLLK | 0.8 | SLWENFK | 128 |
| FLPIIAGIAAKFLPKIFCTISKKC | 9 | FIGMIPGLIGGLISAIK | 30 | KIAKVALKALKIAKGALKAL | 336.8 |
| WKLFKKILKFL | 32 | CKINKQC | 200 | GNPLKLFLPSTWVHFFKFLR | 40 |
| GLSALISWIKRKRQQ | 222.8 | ILSAIWSGIKSLF | 17 | IILLLLKKFWKKL | 12.5 |
| FLKGIKGMLGKLL | 800 | FAKLLAKKLL | 854.7 | VALALKALKKALKKLKKALKKAL | 59.7 |
| LKLKLKLKLKLKLK | 293.3 | GLPVCGETCVGGTCNTPGCSCSWPVCT RN | 31 | VAKKLAKLAKKLAKLAL | 229.3 |
| FLPIVGKLLSGLF | 150 | WKRWVQRWKRFLR | 230 | GIIKKIIKKIIKKIIKKI | 40.7 |
| GYGCPFNQYQCHSHCSGIRGYKGGYCK GTFKQTCKCY | 24 | FPVTWRWWKWWKG | 264.8 | FVDLKKIANIINSIF | 230 |
| QIWHKIRKLWQIIKDGF | 22.8 | GYCFTACYARNGVRICYRRCN | 64 | TLLKKVLKAAAK | 50 |
| RGGRLCYCRRRFCVCV | 41.1 | ALWMTLLKKVLKAAAKAALKAVLVGA NA | 1.2 | GFGSLLGKALKIGTNLL | 145 |
| RGWRRWGRKWAHGWKKYG | 400 | INWKKIASIGKEV | 101 | GVLGTVKNLLIGAGKSAAQSVLKTLSCKLSN DC | 180 |
| VSRRRRRRGGRRRR | 37.2 | LLKKVLKAAA | 101 | CIGAVLKVLTTGLPALISWIKRKRQQC | 3 |
| TIAMRAINNYRWRSKNQNTFLR | 18.7 | INWLKLGKAIIDAL | 11.3 | LNWGAILKKIIK | 200 |
| RPFTRAQIFAIQHISPRTIAMRAINNYRW R | 150 | MAQDIISTIGDLVKWIIDTVNKFPKK | 75 | RRLFRRILRWL | 4.8 |
| KITLKLAIKAWKLALKAA | 10 | KRRKLAFRLFRLFLKLVLKK | 20 | ASVVKKWWKGWWKLLK | 64 |
| FFGHLFRGIINVGKHIHGLLSG | 50 | RRGCFRVCYRGFCFQRCR | 46 | AMTSRKRKFVWYVLSSSLKWLIKKAKKIGV QVCGFE | 16.47 |
| GLKLLLSLGLKLL | 300 | ILPILGNLLNSLL | 300 | GVVDILKGAAKDIAGHLASKVMNKL | 200 |
| GFSSIFRGVAKFASKGLGKDLAKLGVDL VACKISKQC | 150 | GFKDLLKGAKKALVKTVLF | 500 | GLFDIVKKVLRLLK | 24.7 |
| MPLSWFFPRTWGKR | 272.5 | GVIDAAKKVVNVLKNLF | 138.2 | AKVVNKLTGGVAGLLK | 128 |
| IKKIVSKIKKLL | 352 | GFWGKLWEGVKNAI | 22.5 | LNWGAWLKHWWK | 132 |
| YRMWRWRWRWR | 284.3 | ILGKLLSTAAGLLKNL | 55 | GLKALKKVFKGIHKAIKLINKHVK | 95 |
| ALFKTMLKKLGTMAL | 101 | GIPCGESCVFIPCITGAIGCSCKSKVCYR N | 7.1 | LNWGAIKKHIIK | 200 |
| KKRLKKIFKKPMVIGVTIPF | 200 | FIHHIIGGLFSAGKAIHRHHRRRRR | 24.8 | KLKLKLKLKLKL | 5.4 |
| INWKKIASIGKEVLKA | 101 | FFHHIFRGIVHVAKTIHRLVTG | 4 | ALWKTLLKKVLKAAAKAALNAVLVGANA | 2 |
| VKRFKKFFRKLKKSV | 165.2 | CKKIGAVLKVLTTGLPALISWIKRKRQQ KKC | 2.5 | IILLLLKKFLKKL | 6.25 |
| FLSIIAKVLGSLF | 30 | FLPIITNLLGKLL | 35 | ILGKLLSTAWGLLSNL | 11 |
| FPVTFRFFKFFKG | 295.3 | ASVVKKWWWKWWWKLLK | 8 | LKKLLKKLLKKLLKKLLKKLLKKLLKKL | 2.7 |
| RVKRVWPLVIRTVIAGYNLYRAIKKK | 6 | FSFLSRIF | 200 | VFIDILDKMENAIHKAAQAGIG | 42.5 |
| GKWWSLLKHILK | 33 | SKVWRHWRRFWHRAHRKK | 500 | ALWKDILKNLLKAALNEINQIVQ | 3.44 |
| FLRSLLRGAKAIYRGARAGWRG | 22.8 | KFKKFKKFKKFKKFKKFKKFK | 176.1 | GLWSKIKEAAKAAGKAALNAVTGLVNQGDQ PS | 200 |
| GLLKKIKTLL | 1000 | IWLTALKFLGKNLGK | 381.4 | FIGALLRPALKLLAGK | 256 |
| ALWKTMLKKLGTMAL | 101 | GIKDLLKGAAKALVKTVLF | 142 | KLALKLALKALKAALKLA | 10 |
| TKPKGTKPKGTKPKGTKPKG | 300 | FPVTWPTKWWKG | 300 | GLPVCGETCVGGTCYTPGCTCSWPVCTRN | 6.9 |

| | | | | | |
|---|---|---|---|---|---|
| LLKALKKLLKKLL | 30 | RWFRIQLQIRRWRNRR | 38.6 | INWKKIASIGKEVLK | 101 |
| NWKKIASIGKEVLKAL | 101 | KIALKALKALKALGKALKAL | 9.9 | IKLSPKTKKNLKKVLKGAIKGAIAVAKMV | 84.6 |
| GLLSLLSLLGKLL | 10.3 | KWCFRVCYRGICYRRCRG | 107.1 | FALALKALKKLAKKLKKLAKKAL | 6.4 |
| WKRWVQRWKRWLR | 167.5 | GIGKFLHSAGKFGKAFLGEVMKS | 200 | KIFGAIWPLALGALKNLIK | 4 |
| KIKIPWGKVKDFLVGGMKAV | 321.4 | GILSSLLKKWKKIIAK | 400 | FAKLLAKALKKFAL | 252 |
| GIMSSLMKKLKKIIAK | 400 | GLLDFFKGAGKELLTHLASQI | 109.5 | RGGRLCYCRRRFCVCA | 41.7 |
| SMLSVLKNLGKVGLGFVACKINKQC | 140 | GLSLLLSLGLKLL | 300 | KILRGVCKKIMRTFLRRISKDILTGKK | 279.4 |
| LIGLVSKGTCVLVKTVCKKVLKQ | 15 | AGGKRIVQRIKDFLRGAGGKRIVQRIK DFLRG | 40 | KFAKKFAKKFAKKFAKKFAK | 104.7 |
| KFAKKFAKKFAKKFAK | 130.6 | FLSLIPKIATGIAALAKHL | 23 | GIPCGESCVWIPCITSAIGCSCKSKVCYRN | 8.91 |
| LLGMIPVAITAISALSKL | 3.3 | KNWGKILKHIIK | 200 | GLLSLLSLAGKLL | 14 |
| GLFGKLQKKFGRKAISYAVKKARGKH | 36 | RRRRRR | 1000 | KWCFKVCYKGICYKKCKG | 32 |
| GLLKRIKFLL | 79.9 | FWGFLGKLAMKAVPSLIGGNKSSSK | 167.4 | ILKWKWPWWKWRR | 401.2 |
| RRWLWLRRWLWR | 29.3 | FPFLKLSLKIPKSAIKSAIKRL | 1000 | FVDLKKIANIINSIFGK | 99.08 |
| KWKVFKKIEKMIRNIRNKIAK | 250 | INWSSIFESVSNLV | 101 | VKRWKKWWRKLKKSV | 153.8 |
| NKWKKILGKIIKVVK | 200 | GALSALKALGKLL | 50 | FLPLLLSALPSFLCLVFKKC | 5 |
| TRSSRAGLQFPVGRVHRLLRK | 164.5 | RFGRFLRKILRFLKK | 109.3 | ENFFKRIRRAGKRIRDAIISAAPAVETLAQAQ KIIKGGD | 1000 |
| GILDTIKNAAKTVAVGLLEKIKCKMTGC | 150 | FAKLLAKLAKKLL | 13.2 | ALWKTLLKNVGKAAGKAVLNAVTDMVNQ | 70.7 |
| FFPLALLCKVFKKC | 300 | SMLSVLKNLGKVGLGFVACKIKKQC | 90 | VKRFKKFFRKFKKFV | 155.5 |
| FLGELWNVWKSVF | 25 | LRFIKKILKKLI | 125 | FLPILAGLAAKLVPKVFCSITKKC | 5 |
| CLLKKLLKKC | 823 | AGTKDWLNKAKDFIKEKGLGILSAAAN AAIN | 200 | YKLLKLLLPKLKPLLIKL | 31.8 |
| FLPFVGNLLKGLL | 150 | GLFEVIKKVASVIGGL | 19.7 | RFARRFARRFARRFARRFARRFARRFARRFAR | 4 |
| VKKFPWWWAFLKK | 65 | KKYLLPIVGNLLKSLL | 31.9 | HWRRFWHR | 500 |
| EAQNLEKEIAALEQAIQGLEKEIPALAQ QIQALEL | 250 | FAKLLAKLAKAKL | 659.7 | LKLSPKTKKTLKKVLKGAIKGAIAIASMA | 192.7 |
| GLLKRIKVLL | 263.2 | RWRWRWRW | 100 | VLPKVMAHMK | 101 |
| HFLLTLKNLAKKIL | 330 | GLWKKIKNVAKAAGKAALGAL | 19.39 | GIMSSLMKKLAKHIAK | 400 |
| KIAKIAGKIAKIAGKIA | 148.9 | VNWKKSLGKSIKVVK | 31.3 | FPVTWPTKWLKG | 336.7 |
| TLYKKFRHFLKKPWHVVGKI | 200 | KWCVYAYVKVKGVLVKYKKCW | 63 | FLPLVTGLLSGLL | 300 |
| KRIWQRIKDFLR | 340 | GLLKKLLKIAAKVGKKLL | 25 | FLWALFKVASK | 76.5 |
| VRRFPAWAPFLRR | 65 | ACYCRRRFCVCVGR | 47.3 | GKWMSLLKHWLK | 106 |
| WLRRIKAWLRRKRK | 256 | SNWKKIASIGKEVLKAL | 101 | INWKKLLDAAKQIL | 26 |
| IKKIWSKIKKWLK | 500 | FILYGLIRFGRLLRK | 10 | FLPLLASLFSRLF | 12.5 |
| IISTIGKLVKWIIDTV | 15 | GKWMKLLKKILK | 339 | LKLKLKLKLKLKLKLKLK | 228.6 |
| GFMKYIKPLIPHAVKAIKKLI | 200 | GLPALISWIKRKRQQG | 53.3 | KWCFRVCYSGICYSRCSG | 15.3 |
| FLPIIAGAAAKVVEKIFCAISKKC | 13 | GFCGYACYYRNGVRICVRYCN | 128 | VKRWKKWFRKWKKWV | 144.5 |
| GRFKRFRKKFKKLFKKLSPVIPLLHL | 75 | KLFKKLFKKLFK | 395 | KRFKKFFKKLK | 5.98 |
| IKNAKVCVYAVCVSHK | 128 | GILNTIKSIASKLKRKAK | 640 | VQLRIRVAVIRA | 292.5 |
| GVLDTFKDVAIGVAKGAGTGVLKALLC KLDKSC | 480 | FLKWALKVAAK | 512 | GVFGLLAKAALKGASKLIPHLLPSRQQ | 64 |
| KKLIKKILKIL | 299.4 | FLSSIGKILGNLL | 120 | WKRIVRWIKRWLR | 14 |
| GLGSLLGKAFKFGLKTVGKMMGGAPR EQ | 70 | FAKKLAKKLKKLAKLALAK | 18.7 | QIYKKYRTWYRKAWKVLGKV | 200 |
| INWKKIFEKVSNLV | 101 | KLLKRIKTLL | 249.4 | FKKLKKIANIINSIFKK | 1000 |
| FAKKLAKKLKKLAKKLAKKWKL | 42.2 | INWKALLDAAKKVL | 34 | KWARLWRWFRITRWLWYIK | 5 |

| | | | | | |
|---|---|---|---|---|---|
| ILPILAPLIGGLL | 450 | GIPCGESCVWIPCISAALGCSCKNKVCYRN | 1000 | ALWMTLLKKVLKAAAKAALNAVLVGANA | 1.4 |
| ILGIITSLLKSL | 31 | FLPLKKLRFGLL | 134.1 | PLLKKLLKKP | 415.6 |
| SKVWRHWRRFWHRAHRLH | 500 | GLSLLLKLGLKLL | 25 | KLKKLLKKLLKL | 335.3 |
| CVRRFPWWWPFLRRC | 3.75 | LRLRLRLRLRLRLR | 263 | ILPWKWCWWCWRR | 32.1 |
| FLRFIGSVIHGIGHLVHHIGVAL | 64 | ALWDTLLKKVLKAAAKAALDAVLVGANA | 5 | VRRFPWWYPFLRR | 65 |
| ALWKKILKNAGKAALNKINQIVQ | 14.98 | WKIFKLKLRMLW | 6.25 | DAACAAHCLWR | 161 |
| YKLLKKLLKKLKKLLKKL | 2.7 | ASVVKKLTKGVAGLLK | 128 | FKRIVQLLKKLLR | 244.6 |
| FALGAVTKVLPKLFCLITRKC | 53.12 | INWKKIASIGKEVLKAL | 80 | LFGFLIPLLPHIIGAIPQVIGAIR | 0.19 |
| HFLGTLVNLAKKIL | 68.9 | INWLKLGKMVIDAL | 24.18 | KNWGAILKHIIK | 200 |
| VRRYPWWWPYLRR | 112.3 | GLLSSLGKLL | 300 | KIWFQNRRMKWKK | 128 |
| GFGALFKFLAKKVAKTVAKQAAKQGAKYVVNKQME | 20.5 | KWCFRVCYRGICYSRCRG | 64 | RQIKIWFQNRRLKWKK | 128 |
| GILSSFKGVAKGVAKNLAGKLLDELKCKITGC | 290 | PRRRRSSSRP | 79.8 | FFHHIFRAIVHVGKTIHRLVTG | 4 |
| GIGAVLLVLTTGLPALISWIKRKRQQ | 76.7 | SKWMSLLKHILK | 87 | GLWKSLFKNVGKAAGKAALNAVTDMVNQ | 173.1 |
| GFGCPFNQGQCHKHCQSIRRRGGYCDGFLKTRCVCYR | 66.85 | VWPLVIRTVIAGYNLYRAIKKK | 11 | GFWSSVWDGAKNVGTAIIKNAKVCVYAVCVSHK | 64 |
| VNWKKILGKSIKVSK | 69 | RWGKWFKKATHVGKHVGKAALTAYL | 89 | NILNTIINLAKKIL | 300 |
| GILSSLLKKLKKIIAK | 284 | GLLSLLSLGLKLL | 7 | AKRHHGYKRKFH | 120.8 |
| GWGSIFKHGRHAAKHIGHAAVNHYL | 91.8 | GVGKFLHSAKKFGQALVSEIMKS | 200 | FLGSLLGLVGKVVPTLFCKISKKC | 78.5 |
| INWKKMAATALKMI | 37 | FFHHIFRGIVHVGKKIHRLVKG | 200 | RGLRRLGRKIAHGVKKYGPTVLRIIRIAGGGGGSC | 146 |
| LLGMIKVAITAISALSKL | 50.2 | RSTEDIIKSISGGGFLNAMNA | 116 | GLLKKIKRLL | 165.7 |
| GLLKIIKTLL | 25.5 | FKRIVQRILDFLR | 260.1 | TLISWIKNKRKQRPRVSRRRRRRGGRRRR | 264.4 |
| SMLGVLKNLGKVGLGFVACKINKQC | 200 | KFKKFKKFKKFKKFKKFKKFKKFKKFKKFK | 123.5 | FAKALAKLAKKLL | 208.3 |
| RGLRRLGRKIAHGVKKY | 101 | GMWSKILPHLIR | 200 | GIWKKWIKKWLKKLLKKLWKKG | 28.7 |
| GIIKKIIKKIIKKI | 1000 | VRRFPYYWPFLRR | 65 | GLKKIFKAGLGSLKKGIAAHVAS | 400 |
| FAKLLAKALKKAL | 347.2 | GLASTIGSLLGKFAKGGAQAFLQPK | 160 | GILSSLWKKLKKIIAK | 197 |
| LKLSPKTKDTLKKVLKGAIKGAIAIASMA | 161 | KALWKTMLKKLGTMAL | 101 | KFKKLFKKLSPVFKRIVQRIKDFLR | 200 |
| FFHHIFRGIVHVGKTVHRLVTG | 40 | GIMSSLMKKLKKIIKK | 400 | FLPLIAGVAANFLPKIFCLISKKC | 5 |
| GLWDVIKKVASVIGGL | 10.2 | IDWKKLLDAAKQIL | 0.45 | GKPTCGETCFKGKCYTPGCTCSYPLCKKD | 64 |
| IFGAIWSGIKSLF | 48 | FAKLWAKLAFGKGIGKVGKKLL | 375.3 | IKLSPKTKDNLKKVLKGAIKGAIAVAKMV | 185 |
| GLLKFIKTLL | 65.8 | FILNILDK | 128 | GIMNTVKDVATGVATHLLNMVKCKITGC | 90 |
| GVIIKKLKGAAKKVAAKLL | 1000 | FLGALLKIGAKVLPSVLCGIFKKC | 16.62 | GLASLLSLLGKLL | 3 |
| ILPIRSLIKKLL | 103.4 | IFGAILPLALGALKNLIK | 18 | FLGALFKVASKLVPAAICSISKKC | 96 |
| GKWMSALKHILK | 200 | FLPLIASLAANFVPKIFCKITKKC | 75 | LNWGAILKHIKK | 200 |
| FFPIIAGMAAKLIPSLFCKITKKC | 7 | FFHHIFRPIVHVPKTIHRLVTG | 200.3 | FLGSIVGALASALPSLISKIRN | 12 |
| AKVVNKLTGKVAGLLK | 128 | GIGGALLSFGKSALKGLAKGLAEHF | 25.9 | IWRIFRRIFRIFIRF | 15 |
| FLPIIAGMAAKVICAITKKC | 25 | SAVGRHLRRFGLRKHRKH | 452.55 | ILGTILGLLKSL | 4 |
| CVKVSVKVGSGVKVSVKVC | 589 | GIFDVLKNLAKGVITSLAS | 98 | HFLGTLKNLAKKIL | 330 |
| WKRWVRRWKRWLR | 138 | FPVTWPTKWWKS | 314.9 | FFHHIFRPIVHVAKTIHRLVTG | 32 |
| SAVWRWRRFWLRKRK | 500 | GLKDWWNKHKDKIVKVVKEMGKAGINAA | 11.53 | GTPCGESCVYIPCISGVIGCSCTDKVCYLN | 27.4 |
| FKRIVQIIKKFLR | 210.6 | YKLLKLLLPKLKGLLPKL | 59.1 | LLGDFFRKSKEKIGKEFKRIVQRIKDFLRNLVPRTES | 9.7 |
| KKLFKKILKFL | 284.9 | KLKLKLKLKLKLKLKLKLKLKL | 0.3 | SMLKVLKNLGKVGLGFVACKINKQC | 120 |
| IAKIAGKIAKIAGKIA | 160.9 | LLRRLLRRLLRRLLRR | 5.5 | ILPWKWKWWKWRR | 401.2 |

| | | | | | |
|---|---|---|---|---|---|
| FIHHIIGGLFSIGKIIHRLIRRRRR | 0.4 | RWFKIQLQIRRWKNKK | 64.4 | GMASTAGSVLGKLAKAVAIGAL | 200 |
| IAGKIAKIAGKIAKIAGKIAK | 32.1 | GILKTIKSIASKVANTVQKLKRKAKNAV | 824 | FLRFAGSVIHGAGHLVHHIGVAL | 334 |
| VRRFPAWWPFLRR | 65 | GIGKFLHSAKKFGKAFV | 101 | GVLDTLKNVAIGVAKGAGTGVLKALLCQLDKSC | 50 |
| GLLDFVTGVGKDIFAQLIKQI | 14.3 | GKWKKLLKKILK | 400 | RVKRVWPLVIRTVIALYNLYRAIKKK | 3 |
| KFHHIFRGIVHVGKTIHRLVTG | 100 | FLPGILKVAAK | 381.4 | VNWKKILGKI | 200 |
| GWLDVAKKIGKAAFNVAKNFI | 26 | GLNAFKKVFQGIHEAIKLINNHVQ | 440 | INWKKIFEKVKDLV | 100 |
| RGLRRLGRKIAHGVKKYGPTVKRIKRKA | 101 | SMLSVLRNLGKVGLGFVACKINKQC | 200 | RWRWRW | 210 |
| KKLFKKILKYL | 101 | RAVIYKIPYNAIASRWIIAPKKC | 200 | CGRRWWGWRCRL | 602.4 |
| GVLKRIKTLV | 500 | GLNALKKVFQGIHEAIKLINKHVQ | 270 | KWAKKWKWFAKAAWKWYKK | 16.2 |
| WRAWAKIYHGVGKLLKGVGRW | 191.3 | GKIIKVVK | 200 | KLLAKAALKWLLKALKAA | 1.7 |
| AFLYRLTRQIRPWWRWLYKW | 33.7 | WKKILSKIKKLLK | 500 | GILSKLLKKLKKIIAK | 400 |
| GKWKKILGHLIR | 200 | FLSLIPKIAGGIAALVKNL | 32 | SFLTTVKKLVTNLAALAGTVIDTIKCKVTGGC | 63.9 |
| KLKLLLKLK | 146 | RVCSWIPLPICH | 260.2 | KILKLLKKFLKKL | 500 |
| QIIKKVRTWIKKAWHLIGKI | 101 | GVIKAAKKVVKVLKKLF | 135.2 | SMLSVLKNLGKVGLGFVAAKIAKQA | 200 |
| FAKFLAKFLKKAL | 38.7 | ILPWKWKWWPWRR | 182.4 | SLWENFKNAGKKFILNILDKIRCRVAGGCRT | 102.4 |
| GLAANFLPKIFCKITRKC | 71.3 | DLWNSIKDMAAAAGRAALNAVTGMVNQ | 692 | LKLKLKLKLKLKLKLK | 256.9 |
| FLYIVAKLLSGLL | 526.5 | VSAVAKVAMKKGAALLKKMGVKISPLK | 361.5 | GLPALISWIKRKRQ | 71 |
| HFLLKLVNLAKKIL | 100 | RLIKRIKTWYRKAWKVVGKF | 200 | SILSLFKMGAKALGKTLIKQAGKAGAEYVACKATNQC | 10 |
| FLQKIIGALGHLF | 69 | FDLLGLVKSVVSAL | 64 | GALSALKALGKAL | 101 |
| GINLKRKGNIMKKVKNIFHKIANADPMIWGYVMLSESK | 41.31 | FAKKLAKKLAKLL | 501 | GVPICGETCTLGTCYTAGCSCSWPVCTRN | 33.04 |
| FKLRAKIKVRLRAKIKL | 64.3 | INWIKIGKKIIASL | 185 | GLWDSIKNFGKTIALNVMDKIKCKIGGGCPP | 64 |
| RRWPWWPWRR | 200 | HFLLTLVNKAKKIL | 500 | RWRWRWRWRW | 76 |
| LNWGAKLKHIIK | 200 | FHAWAKLLKGVGRFFKGIGRW | 19.8 | HFLGTLVNLKKKIL | 500 |
| ILSAIWSGIKGLL | 138.4 | GVWTTILGGLKKFAKGGLEALTNPK | 60 | KWRRWIRWL | 286.1 |
| FALAAKALKKLAKKLKKLAKKAL | 119.1 | FAKKLAKLAKKLLAL | 208.2 | FKRIVQLIKDFLR | 67 |
| GLLGLLGSVVSHVVPAIVGHF | 25 | ITEVITILLNRLTDRLEK | 250 | GFLSILKKVLAKVMAHMK | 29.7 |
| ASVVKWLWKWVWKLLK | 64 | VRRFPWAWPFLRR | 65 | AKVVNKLTKKVAKLLK | 128 |
| GLKALKKVFQGIHKAIKLINNHVQ | 250 | GVGKFLHSAKKFGQALASEIMKS | 200 | GFLGPLLKLAAKGVAKVIPHLIPSRQQ | 90 |
| KWKLFKKIEKVGQNIRDGIIKAGPAVAWVGQATQIAK | 24.5 | LLIILRRRWRKQARARSK | 200 | LLGPVLGLVSNVLGGLL | 308.6 |
| AGWKEWLNKAKDFIKEKGLGMLSAAANAALN | 101 | KWKVFKKIEKMGRNIRNGIVKAGPKWKVFKKIEK | 101 | FLSLIPAAISAVSALANHF | 52.17 |
| FLSHIAGFLSNLF | 80 | RWCFRVCYRGICYRKCR | 54.8 | KLKLKLKLKLKLK | 5.6 |
| GIGKFLHAAKKFAKAFVAEIMNS | 69.9 | SISCGESCAMISFCFTEVIGCSCKNKVCYLN | 55.3 | LRFLKKILKHLF | 30 |
| LLPIVGNLLKSLL | 120 | GKAMSLLKHILK | 200 | GVIIDTLKGAAKTVAAELLRKAHCKLTNSC | 700 |
| RPFTRAQWFAIIHISPRTIAMRAINNYRWR | 101 | SMLSVLKNLGKVGLGFVASKINKQS | 200 | GILDTFKGVAKGVAKDLAVHMLENLKCKMTGC | 101 |
| AVDLAKIANKVLSSLF | 250 | GILSSFKGVAKGVAKDLAGKLLETLK | 200 | RRWRIVVIRVRR | 50 |
| LKLKLKLKLKLKLKLKLKLK | 205.9 | FKAFKAFKAFKAFKAFKAFKAFKA | 179.5 | FAKKLLAKALKL | 730.4 |
| ILPWKWPWWKWRR | 249.6 | GFKRLVQRLKDFLRNLV | 900 | LWGALLGLGSTLLSKL | 4.7 |
| RLYRRLYR | 1000 | FDVMGIIKKIASAL | 181.1 | MMRVMRRKTKVIWEKKDFIGLYSID | 95.4 |
| GKWVKLLKKILK | 400 | GWASSIGSILGKFAKGGAQAFLQPK | 256 | IWSFLIKAATKLLPSLFGG | 248.5 |
| GILKSLLKKLKKIIAK | 400 | LLIILRRRIRKQAHAHSK | 200 | INLKAIAALVKKV | 725.2 |
| RGLRRLGRKIAHGVKKYGATVLRIIRIA | 100 | KIAKVALKALKIAKVALKAL | 267.5 | FFHHIFRAIVHVPKTIHRLVTG | 75 |

| | | | | | |
|---|---|---|---|---|---|
| IFGAIWPLALGALKNLIK | 4 | RGLRRLGRKIAHGVKKYG | 400 | FLPLLAGVVANFLPQIICKIARKC | 38.1 |
| ILPAKAPAAPARR | 147.4 | KLALKAAAKAWKAAAKAA | 200 | GFKMALKLLKKVL | 199 |
| VNWKKILKKIIKVAK | 88.2 | KRIVKLIKKWLR | 800 | LNWGAILKHKIK | 200 |
| MKKLLLILFCLALALAGCKKAP | 171.6 | GILKTIKSIASKVANTVQKLKRKAKNAVA | 665 | LKLKLKLKLKLKLKL | 0.5 |
| GVWSTILGGLKKFAKGGLDAIVNPK | 64 | GMLKRIKTLL | 400 | HHIIGGLFSAGKAIHRLIRRRRR | 27.6 |
| MLLKKLLKKM | 785.5 | VRRFAWWWPFLRR | 526 | FLSLALAALPKFLCLVFKKC | 17 |
| GIKKFLHIIWKFIKAFVGEIMNS | 2.9 | GLLKRIKALL | 84.3 | IKLSKETKKNLKKVLKGAIKGAIAVAKMV | 126.4 |
| FWGALAKGALKLIVSLFSSFSKKD | 6.4 | RRRWWWWL | 60 | FLPILGNLLNGLL | 150 |
| KFKKFKKFKKFKKFKKFKKFKKFKKFK | 137.2 | IKKFKKPFKPFRR | 114.6 | GLNALKKVFQGIHEAIKLFNNHVQ | 430 |
| ALLKTMLKKLGTMAL | 101 | RLYRKVYGRLYRKVYGRLYRKVYGRLYRKVYGKKK | 610 | GLPTCGETCTLGKCNTPKCTCNWPICYKD | 64 |
| KQLKKVSAVAKVAMKKGAALLKKMGVK | 350.5 | INWLKLGKKILGAL | 45 | YGRKKRRQRRR | 300 |
| GIWDTIKSMGKVFAGKILQNL | 90 | KWCFRVCYSGICYRRCRG | 64 | FKVTWKTKWWKG | 600 |
| GILSSFKGVAKGVAKDLAGKLLETLKCKITGC | 160 | FFGSLLKLLPKLL | 49.2 | GLPTCGETCTLGKCNTPKCTCNWPICYKN | 64 |
| FLSKIWDGVKSLL | 39.3 | RWFKIQMQIRRWKNKK | 128 | KRIVKLILKWLR | 96 |
| HFLLTLVNLAKKKL | 500 | KLLKVIKKLL | 189.2 | KRRLFLFRLFRLFLRLFLKK | 8 |
| RGLRRLGRKIAHGVKKYGPTVLRIIRIAG | 23.1 | WRRFWRR | 500 | INWKKIASIGKEVL | 101 |
| FLKGIVGMLGKLF | 26 | FLSTALKVAANVVPTLFCKITKKC | 150 | LKKLLKKLLK | 1000 |
| GWKKWLRKGAKHLGQAAIK | 116.3 | KRRKLIKILKLIAKLIRKKR | 40 | KIGKALGKALKALGKALGKA | 14.9 |
| INWKKGKEVLKAL | 101 | GLMSVLKGVLKTAGKHIFKNVGGSLLDQAKCKISGQC | 125 | FIHHIIGGLFSAGKAIHRLIR | 21.2 |
| KWFKIQLQIKKWKNKK | 90.2 | GFLSILKKVLGKVMAHMK | 41.4 | CGRRWGWWRCRL | 602.4 |
| CGKKWWGWKCKL | 646 | GFLGSLLKTGLKVGSNLL | 150 | FFGAIAAALPHVISAIKNAL | 120 |
| GFFALIPKIISSPLFKTLLSAVGSALS | 35.7 | SIPCGESCVFIPCTVTALLGCSCKSKVCYKN | 405 | IISTIGDLVKWIIKTV | 40 |
| SMLSVLKNLKKVKLKFVACKINKQC | 200 | INWLKLGKKILGAI | 75 | NVWKKILGKIIKVAK | 200 |
| NVWKKVLGKIIKVAK | 200 | LNLKGLIKKVASLLN | 530.3 | ILGAILPLVSGLLSNKL | 115 |
| RVRRFPWWWPFLRRR | 60.9 | FKAFKAFKAFKAFKAFKAFKA | 204.9 | GKWMSLLKHILK | 78 |
| FLSLIPKAISAISALINHF | 9.266 | RGGRLCYCRRRFCVCVGR | 37.1 | YRWWRWARRW | 437.2 |
| GFGSLLGKALRLGANVL | 140 | LKLLLLKLLKLKLWK | 5 | KWKLFKKIFKRIVQRIKDFLRN | 76.2 |
| FLPLAVSLAANFLPKLFCKITKKC | 1.9 | KKPFKFPKKPFRR | 115.7 | KAAKKAAKKAAKKAAKKAAKKAAKKAAK | 178.3 |
| VRRFPYWYPFLRR | 65 | ICYCRRRFCVCVGR | 46.2 | FVDLKKIANILNSIF | 152.6 |
| FFPLVLGALGSILPKIF | 4.8 | AFALIAGALYRIFHRR | 187.2 | LDVKKIICVACKIKPNPACKKICPK | 200 |
| AKKVFKRLPKLPSKIWNWK | 800 | RFGRFLRKILRFLRK | 72.4 | IKKIIKKIIKKI | 1000 |
| KFKKFKKFKKFKKFKKFKKFK | 154.2 | ALWMTLLKKVLK | 100 | GIWKTIKSMGKVFAGAILQNL | 40 |
| GVVTDLLKTAGKLLGNLFGSLSG | 100 | KTKLFKKFAKKLAKKLKKLAKKL | 330 | FFGWLIKGAIHAGKAIHGLIHRRRH | 1 |
| LNWGAGLKHGGK | 200 | FFHHAFRGIVHVGKTIHRLVTG | 500 | FLSTIWNGIKSLF | 24.48 |
| AKVVNKLTKGVAKLLK | 128 | FVKELWDKVKKMGSAAWSAAKGAFA | 250 | RAGLQFPVGRVHRLLRK | 199.9 |
| INWKKLGKKILGAL | 140 | KIASIGKEVLKAL | 101 | GMWSKIPGHLIR | 200 |
| LNWGAVLKHVVK | 200 | SAVGRHGRRFGLRKHRKH | 300 | GLLGPLLKIAKKVGSNLL | 350 |
| KVKVKVKVPPTKVKVKVK | 1000 | FVGAIAAALPHVISAIKNAL | 120 | GFKDLLKGAAKALVKAVLF | 22 |
| ILCWKCPWWPWRR | 38.2 | KLALKLALKAWKAALKLA | 11 | ILGPVISTIGNALGGLLKNL | 31.4 |
| FLGAIAQALTSLLGKL | 22 | GFWGKLFKLGLHGIGLLHLHL | 26.9 | GIFSALAAGVKLLGNTLFKMAGKAGAEHLACKATNQC | 35 |
| LKFLKFG | 150 | KRIVQRIKDWLRKLCKKW | 83.5 | GLPVCGETCFGGTCNTPGCSCTWPICTRD | 11.5 |

| | | | | | |
|---|---|---|---|---|---|
| MQFITDLIKKAVDFFKGLFGNK | 1.2 | IDWSKIFEKVKNLV | 100 | KVMAHMK | 101 |
| GLKEVLHSTKKFAKGFITGLTGQ | 160 | ILGKLLKTAAGLLSNL | 28 | GFGMALKLLKKVL | 165 |
| GKWLKLLKKILK | 196 | FMGSALRIAAKVLPAALCQIFKKC | 77.6 | FPVTWGWWKWWKG | 279.5 |
| GLLKPLLKIAAKVGSNLL | 500 | ALWKSLLKNVGKA | 512 | FALALKKALKALKKAL | 114.2 |
| INWKKIFEKVKNLV | 101 | GLRSKIWLWVLLMIWQESNKFKKM | 175 | FIRRIARLLRRIF | 886.5 |
| KRFWPLVPVAINTVAAGINLYKAIRRK | 9 | GKVLDKFGKIVGKVLKQLKKVSAVAKV | 117.4 | IAKIAGKIAKIAGKIAKIAGKIAK | 5 |
| VNWKKVLAKIIKVVK | 200 | KRLFKKELLFSLRKY | 59.2 | GSKRWRKFEKRVKKVFEHTKEALPVIQGVATVVGAVGRR | 112.5 |
| GTKEWLNKAKDFIKEKGLGMLSAAANAALN | 150 | FKLAFKLAKKAFL | 645.2 | KWCFRVCYRGICYRRCR | 128 |
| LLSLALAALPKLFCLIFKKC | 180.5 | SWKSMAKKLKEYMEKLKQRA | 51.6 | ASVVKKLTKKVAGLLK | 128 |
| PKVMAHMK | 101 | SAVWRHWRRFWLRKHRKH | 500 | FFRRFFRRFFRR | 268.4 |
| FAKKLAKLAKKLAKLALAL | 37.6 | KIFKKFKDWFKKAFHVLGKV | 101 | LNKGAILKHIIK | 200 |
| IWNKIAKSIGKVLEKAL | 101 | FAVTWATKWWKG | 300 | GIAGLLRSFVRMLAKIMGG | 64.3 |
| ASVVNKLTKKVAKLLK | 128 | VNWKKILAKIIKVAK | 66.5 | RWRRFWRR | 500 |
| FAKKLAKKLKKLAKLALAL | 119.4 | KWCFRVCYRGICYRRCSG | 64 | GIWKKWIKKWLKKLKNLF | 107.5 |
| IDWLKLGKMVMDVL | 50 | FLPIIAGVAAKVLPKLFCAITKKC | 5 | FLPILGKLLSGIL | 8 |
| IKWKKLLRAAKRIL | 75 | FVKLKKILNIINSIFKK | 86.99 | KIWFQNKKMKWKK | 128 |
| GIPCAESCVYIPCTVTALLGCSCSNRVCYN | 61.7 | IKKILSKIKKWWK | 500 | GIMSSLMKKLKKHIAK | 400 |
| WKRIVRRIWRWLR | 12.5 | KLKKKLKKLLKK | 328.7 | GVIIKTL | 1000 |
| GKWMSFLKHILK | 69 | ALWKTMLKKLGTMALHAGKAALGAAADTISQGTQ | 29 | GMWSKILGHLIR | 82 |
| GLLKRIKTLL | 1000 | ILGKLLSTWAGLLSNL | 39 | GLLSAAKAAGKLL | 50 |
| FPVTWRTKWWKG | 300 | FRIRVRVAKKFGKAFVGEIM | 128 | VRRFPAAWPFLRR | 65 |
| VKRWKKWRWKWKKWV | 142 | FFGSVLKLIPKIL | 124.8 | GKLEVLHSTKKFAKGFITGLTGQ | 160 |
| GIMDTIKGAAKDLAGQLLDKLKCKITKC | 200 | GKWMSLWKHILK | 101 | GMWSKILGHL | 200 |
| SKVGRHLRRFGHRAHRKL | 500 | FLPVLAGLTPSIVPKLVCLLTKKC | 80 | FALALKALKKLKKALKKAL | 334.3 |
| SAVWRRWRRFWLRKRRKR | 500 | SVIWRKLFFIFIKRSGNWIKKVEKRQNLL | 10.51 | KLLKFIKTLL | 186 |
| INWLKLGKKMMSAI | 101 | GIGAVLTTGLPALISWIKRKRQQ | 6.5 | GWLDVAKKIGKAAFNVAKNFL | 37.4 |
| KAACAAHCLWR | 159.4 | WSPQEEDRIIEGGI | 123 | KKWWKWWKKWWRR | 45.3 |
| GKWMSLLKKILK | 126 | GLLPLLSLLGKLL | 2 | ILGKLLSTAAGLLSNL | 137.8 |
| GIPCGESCVWIPCISSAIGCSCKSKVCYRN | 5.1 | ENFFKEIERAGQRIRDAIISAAPAVETLAQAQKIIKGGD | 115 | RRRWWWWV | 130 |
| LLKKLLKKLLKKC | 252.2 | FLRALWNVAKSVF | 12.5 | GRRKRKWLRRIGKGVKIIGGAALDHL | 85.9 |
| FLIGMTQGLICLITRKC | 48 | RFRRLRKKWRKRLKKI | 256 | AAAKAALNAVLVGANA | 40 |
| FLPGILKVAANVVPGVICAITKKC | 145.8 | GLKLLLKLGLKLL | 25 | GIGKFIHSVKKWGKTFIGEIMNS | 56 |
| IAKIAGKIAKIAGKIAK | 148.9 | FIHHIIGGLFSAGKAIHSLIHRRRR | 17.3 | FFHHIFRGIVHVPKTIHRLVTG | 200.3 |
| GLMSVLKGVLKTAGKHIFKNVGGSLLDQAKCKITGQC | 100 | LRFLRRILRRLL | 30 | IIPLPLKKFAKKT | 500 |
| IDWKKIFEKVKNLV | 101 | FAKLLAKALKKL | 474.8 | LALKALLAVAKKIL | 256 |
| GMWSKILGHLI | 122 | KWKVFKKIEKNGRNIRNGIVKAGPAIAVLGEAKAL | 26.2 | GLLGPLLKIAAKVGSNLL | 140 |
| RWWLRRIW | 200 | GFLGPLLKLGLKGAAKLLPQLLPSRQQ | 50 | VNSKKISPKSIKVSK | 82 |
| GKFMSLLKHILK | 200 | WLLKKLLKKW | 262.8 | FLGLLFHGVHHVGKWIHGLIHGHH | 5.8 |
| HYIRHLKTWFHKPFKLIGKV | 200 | KFAFKFAFKFAFKFAFKFAFKFAFKFAF | 144.1 | VNWKKLLGKLLKVVK | 71.5 |
| GLMDTVKNAAKNLAGQLLDTIKCKMTGC | 100 | ASVVNKLTGKVAGLLK | 128 | LVIRTVIAGYNLYRAIKKK | 360 |
| FLHFLHHLF | 336.9 | WIRRIKKWIRRVHK | 256 | SLWENFKNAGK | 128 |

| | | | | | |
|---|---|---|---|---|---|
| KRFKKFFKKPK | 95.92 | GLLGPLLKIAAKVGKKLL | 60 | GLKKWFKKAVHVGKKVGKVALNAYL | 91.2 |
| FIHHAAGGLFSAGKAIHRLIRRRRR | 69.1 | GLLDTFKNLALNAAKSAGVSVLNSLSC KLSKTC | 140 | ILGKLLSTAWGLLSKL | 6 |
| FPVTWRWWRWWRG | 65 | GMASKAGSVLGKITKIALGAL | 200 | FLGALLKIGAKLLPSVVGLFKKKQQ | 61.7 |
| FLSMIPHIVSGVAALAKHL | 33 | AAGLAMLFLGILSAAGSTMGARA | 40 | FAKLWAKLAKKL | 693.5 |
| CFKFKFKFGSGFKFKFKFC | 35 | IAKIAGKIAKIAGKIAKIAGKIA | 3.1 | RKIWWWWL | 57 |
| GLNALKKVFQGFHEAIKLINNHVQ | 370 | GLLKFIKWLL | 0.6 | KVLKAAAKAALNAVLVGANA | 25 |
| FVDLKKIANIINSIFKK | 1000 | FFHHFARGIVHVGKTIHRLVTG | 150 | GLWRVIRKVASVIGGL | 15.5 |
| GLADYWRTAFRANFANLGPGIRCKSAR C | 64 | LRILRRLLRRLF | 125 | ILGLVISTIGNVLGGLLKNL | 61.5 |
| KIPPVVKDTLKKVAKGVLSAVAGALS | 512 | RGGRLCYCRGWICFCVGR | 38 | INWSSIFESVKNLV | 101 |
| HLLKKWRTWLRKAWHIVGKV | 58.5 | FFFLPSLIGGLVSAIK | 25 | GVVVRVGRVVVRWVRRRR | 90 |
| GIMSSLMKKLAAHIKK | 400 | FLPILASLAAKFGPKLFCLVTKKC | 10 | HFLGKLVNLAKKIL | 100 |
| VKRFKKFFRKFKKSV | 160.2 | GMFTNMLKGIGKLAGKAALGAVKTLA | 76.55 | KALAALLKKWAKLLAALK | 2.5 |
| RRFPWWWPFRR | 142.2 | FWGALAKGALKLIGVGSLFSSFSKKD | 11.5 | GIWKTIKSMGKVFAGAIKQNL | 200 |
| LDVKKIICVACKIRPNPACKKICPK | 200 | FLPLILPSIVTALSSFLKQG | 82 | FAKLLKLAAKKLL | 404.9 |
| FSVTWRWWKWWKG | 266.2 | GVFRRLRKVTRKVLKKIGKVLKWI | 128 | KRAKKFFKKPK | 107 |
| GLLKRIKKLL | 500 | FLPGLIKAAVGVGSTILCKITKKC | 100 | KWKLFKKISKFLHLAKKF | 100 |
| WWRLRRIW | 200 | LRIVKLILKWLR | 22 | GILKKILKKILKKL | 80 |
| GFKDLLKGAAKALVKTVKF | 500 | FLPKILKVAAK | 512 | KAACAAKCLWR | 160.5 |
| GFLGILFHGVHHGRKKALHMNSERRS | 85.8 | GLFDVVKGVLKGVGKNVAGSLLEQLK CKLSGGC | 65.4 | GVIIDTLKGAAKTVAAELLRKAH | 1000 |
| GIPCGESCVFIPCITAAIGCSCKSKVCYR N | 8.4 | IKKWWSKIKKLLK | 500 | KIAGKIAKIAGKIAKIAGKIAKIAGKIA | 0.4 |
| KLLKKAGKLLKKAGKLLKKAG | 101 | FLSLIPHIASGIASLVKNF | 18.6 | SMLSVLKKLGKVGLGFVACKINKQC | 110 |
| GLMDVFKGAAKNLLASALDKIRCKVTK C | 150 | GMMKRIKTMM | 500 | WKKIWSKIKKLLK | 500 |
| MLLKKLLKKLLKKM | 90 | GILSSLLKKLKKWIAK | 115 | FLPLLISALTSLFPKLGK | 15 |
| KFAKKFAKKFAK | 347.2 | FLSLIPKAISAVSALANHF | 197.7 | GLKEVAHSAKKFAKGFISGLTGS | 256 |
| GIGKFLKKAKKFGKAFVKMKK | 100 | IKKIWSKIKKLWK | 500 | SKVWRHWRRFWHRAHRKL | 500 |
| LKKVLKAAA | 101 | AGTKEWLNKAKDFIKEKGPGMLSAAA NAALN | 250 | GIGKFIHAAKKFGKLFIGEIMNS | 32 |
| GLSLLLSLLLGKL | 8 | GLWNSIKIAGKKLFVNVLDKIRCKVAG GC | 33.03 | GLLGPLLKIAAKVGKNLL | 110 |
| IWLTKALKFLGKNLGK | 506.5 | GVIIKTLKGAAKTV | 1000 | ALWKSILKNVGKAAGKAVLNAVTDMVNQ | 128 |
| HFLLTLVNLKKKIL | 500 | GWKDWFRKAKKVGKTVGGLALNHYL | 88.4 | GIMSSLMKKLAAIIAK | 93 |
| RGGRLCYCRRRFCVCY | 39.8 | FLSLLPHIASGIASLVSKF | 15.82 | CGKKPWWWKCKL | 629.7 |
| GFMDTAKNVAKNVAVTLIDKLRCKVTG GC | 135 | FLPIALKALGSIFPKIL | 107.2 | FLPMLAGLAANFLPKIVCKITKKC | 19 |
| GIGKFLHSAGKFGKAFIGEIMKS | 180 | FWSTIWNAAKSLI | 10.6 | GMASKLAKVLPHVVKLIK | 200 |
| KKLFKKILKVL | 334 | KWCFRVCYRGFCYRRCRK | 8.4 | NLLNDALGTVNGLLGRS | 256 |
| FLPLIAGVAASILPKIFCFITKKC | 5 | RQIKIWFQNRRMKWKK | 300 | FAKKLAKKLKKLAKKLAKLALAL | 19.7 |
| ALWKSLLKNVGKAAGKAALNAVTDMV NQ | 216.6 | GLLKRIKLLL | 68 | GLPVCGETCFGGTCNTPGCSCETWPVCSRN | 7.52 |
| KWASLWNWFNITNWLWYIK | 212 | FVKLKKILNIILSIFKK | 28.34 | VAKLLAKLAKKLL | 139.5 |
| IWLLLLKKFLKKL | 50 | VRRFPAAAPFLRR | 65 | IKIPAFVKDTLKKVAKGVISAVAGALTQ | 160 |
| SKVGRHLRRFLHRAHRKL | 493.9 2 | WKRRIKIWKKIR | 256 | GIKSALLGIAKNVGMSLLQKAQCKLSGSC | 49.04 |
| ALWKDILKNAGKAALNEINQIVQ | 138.1 | YWRWRW | 556.6 | AMTLRKRKFAWYVLSSSLKWLIKKAKKIGV QVCGFE | 2.67 |
| FLSLIPKIISAISALINHF | 2.766 | GLFDIWAWWRWRR | 133.5 | KLGFENFLVKALKTVMHVPTSPLL | 256 |
| KRAKKFFKKLK | 7.98 | FWGALAKGALKLIGPGSLFSSFSKKD | 46.6 | KIAKRIWKILRRRL | 66.7 |

| | | | | | |
|---|---|---|---|---|---|
| ILGKIWEGIKSLF | 28.5 | SMLSVLKNLKKVGLGFVACKINKQC | 200 | GWKSVFRKAKKVGKTVGGLALDHYL | 92 |
| FAKKLAKKLL | 843.9 | CGRRPWWWRCRL | 115.9 | GIKKWVKGVAKGVAKDLAKKIL | 200 |
| GFKDLLKKAAKALVKTVLF | 11 | WLRRIGKGVKIIGGAALDHL | 116.5 | GWGCNIFGGNDYRCHRHCKSIRGYKGGYCK LGGICKCY | 128.3 |
| VQWRIRVAVIRK | 657 | SMLSVLKNLGRVGLGFVACKINKQC | 200 | VKRWKKWKRKWKKWV | 145.8 |
| VKWRWKKWRWKWKWKV | 131.2 | LLRRLLRRLLRR | 301.4 | FFHHIFRGVVHVGKTIHRLVTG | 35 |
| WKKWWKKW | 1000 | RIWFQNRRMRWRR | 128 | KWKLFKKIEKVGQNIRDGIIKAGPAVAVVGQ ATQIAK | 101 |
| FLPLLAGLAANFLPKIFCKITRKC | 0.4 | IISTIGKLVKWIIKTV | 11 | KQKTLKKVWKLSEKVLIFASAFAKKAGAAE ATLVL | 61.09 |
| FLQHIIGALGHLF | 115 | LWKIWWKIWRVWKNWR | 1.7 | IKIPSFFRNILKKVGKEAVSLIAGALKQS | 28.1 |
| KIPKPLKKFAKKT | 500 | GLLGPLLKIAAKVGSKLL | 80 | FLAGLIGGLAKML | 40 |
| LRFLKKALKKLF | 125 | FLPIVAKLLSGLL | 39.6 | WWWLRRIW | 20 |
| ILSYLWNGIKSIF | 32.8 | KLKLKLKLK | 225.1 | GIGAVLWVLTTGLPALISWIKRKRQQ | 0.9 |
| GLPVCGETCVGGTCNTPGCTCSWWPVC TRN | 14 | KWKVLKKKIKMLRNRINGLVKAGPAL KVKLQALAL | 250 | KWCFRVCYRGICYRRCRK | 64 |
| GMWSKILGHLKR | 200 | RLYRRLYRRLYRRLYRRLYRRLYRRLYR RLYRKKKKKKK | 1000 | FIAAIIGGLFSAGKAIHRLIRRRRR | 1.7 |
| SNWLKLGKKMMSAL | 101 | QLIKKLRTWYRKAWHVLGKL | 200 | KNWKKIASIGKEVLKAL | 101 |
| GVVDILKGAAKDLAGHLATKVMNKL | 400 | FFHHIFRGKVHVGKTIHRLVTG | 4.8 | GLMDTVKNAAKNLAGQMLDKLKCKITGSC | 90 |
| GYCFTACYLRNGVRICYRRCN | 100 | VNWKKIILGKIIKVVK | 200 | HKLGTLVNLAKKIL | 500 |
| FPVTWPTKWRKG | 327.2 | INWLKLGKKMMSAL | 101 | GIGAVLKVLTTGLPALISWIKFKRQQ | 29.1 |
| FAKLLAKLAKKFAL | 630.1 | VCYCRRRFCVCVGR | 46.6 | GFLSILKKVLP | 101 |
| FAKKLAKLALKLAKL | 53.5 | GFKDLLKGAAKALVKTVLF | 200 | RGGRLCYCRRRFCVCI | 40.8 |
| GFLSILKKVLKVMAHMK | 22.9 | GVVVRVARVVVRWVRR | 80 | GILDKLKEFGISAARGVAQSLLNTASCKLAKT C | 75 |
| FFRRFFRR | 397.8 | YKLLKLLLPKLKGLLIKL | 13.1 | LRFLKKILKKLF | 125 |
| GLLGKLLKIAAKVGSNLL | 15 | FFGTALKIAANVLPTAICKILKKC | 0.6 | AGTKEWLNKAKDFIKEKGLEMLSAAANAAL N | 150 |
| LKWGAILKHIIK | 200 | ALWMTLLKKVLKA | 19.7 | FAKLLAKALKKLL | 13.5 |
| GLKEIFKAGLGSLVKGIAAHVAN | 210 | GLLSALKAAGKLL | 50 | SMLSVLKNLEKVELEFVACKINKQC | 200 |
| GLPTCGETCFGGTCNTPGCSCSSWPICT RN | 7.7 | WKLFKKILKWL | 26 | KKWMSLLKHILK | 133 |
| RIGVLLARLPKLFSLFKLMGKKV | 35 | WSKILGHLIR | 200 | KRIVQRIKDFLR | 80 |
| SLWENFKNAGKKFILNILDKIRCRVAGG CR | 58.9 | GLFDVIKKLLKKIGGL | 9 | FWGLKGLKKFSKKL | 418.4 |
| LKKLLKKLLKKL | 1000 | GVVVRVGRVVVRWVRR | 94 | FLSAITSILGKFF | 37.5 |
| RRRRRRRKDVY | 609 | KLKLKLKLKLKLKLKL | 1 | FFPIIAGMAAKVICAITKKC | 8 |
| FLPAALAGIGGILGKLF | 44 | FLSAITSLLGKLL | 60 | ALWKTMLKKLGTMALHAGKAALGAAADTI | 101 |
| ALWKTMLKKLGTM | 101 | FAKLLAKLAKAKG | 722.5 | FLGALIKGAIHGGRFIHGMIQNHH | 96.6 |
| FAKLLAKLAKKVL | 37.5 | GMATKAGTAFGKAAKAIIGAAL | 256 | GMWSKILGPLIR | 200 |
| IKKILSKIKKWLK | 500 | INFLKLGKKILGAL | 140 | KALWKTLLKKVLKA | 54.3 |
| GLLKAIKTLL | 147 | FPLTWLKWWKWKK | 267.1 | GLPTCGETCFKGKCYTPGCSCSYPICKKN | 64 |
| GIRCPKSWKCKAFKQRVLKRLLAMLRQ HAF | 7 | GIRKWFKKAAHVGKKVGKVALNAYL | 91.2 | FKRWVQRWKRFLR | 256 |
| KIKWILKYWKWS | 400 | LKLFKKILKVL | 190 | RLYRRLYRRLYRRLYRRLYRRLYRRLYR | 48 |
| KRIVKRIKKWLR | 800 | FPLTWPTKWWKG | 318.1 | FAKKLKKLAKKL | 694 |
| GLLKVIKVLL | 28.5 | LCYCRRRFCVCVGR | 46.2 | FIHHIIGGLFSVGKHIHRLIRRRRR | 0.8 |
| GFLSILKKVLPKVMAHMK | 101 | LLKKLLKKC | 899.3 | IWLLLLKKFWKKW | 6.25 |
| GIFSLFKAGAKFFGKHLLKQAGKAGAE HLACKATNQC | 600 | GLPTCGETCFKGKCYTPGCSCSYPICKK D | 64 | VNWKKVLAKIIKVAK | 200 |
| LNWGALLKHLLK | 63 | GLKDIFKAGLGSLVKGIAAHVAN | 101 | FFFLSRIF | 200 |

| | | | | | |
|---|---|---|---|---|---|
| LLIILRRRWRKQAKAKSK | 200 | GTLPCGESCVWIPCISSVVGCACKSKVCYKD | 10 | GIVKKIVKKIVKKI | 1000 |
| ILGKIWEGIKSIF | 34.4 | FRVTWRTKWWKG | 600 | IPPFIKKVLTTVF | 101 |
| VNWKKVLPKIIKVAK | 200 | GKGRWLERIGKAGGIIIGGALDHL | 101.9 | WKLFKKILKVL | 104 |
| VGALAGFLYWHFLRKGTKMVGK | 10 | GKWLSLLKHILK | 65 | FKDLKKIANIINSIFKK | 1000 |
| VKRWKKFWRKWKKWW | 139.1 | IWRIFR | 256 | AGGKRIVKRIKKFLRGAGGKRIVKRIKKFLRG | 10 |
| GMASLWAKVLPHVVKLIK | 162.5 | GMLKRIKTLM | 500 | GVIKAAKKVVNVLKNLF | 137.2 |
| RRLFRRILRYL | 50.1 | VNWRRILGRIIRVVR | 39.9 | KLALKAALKAWKAAAKLA | 107 |
| FFHHIFRGIVHIGKTIHRLVTG | 7 | SLWENFKNAGKK | 128 | IIKKIIKKIIKK | 1000 |
| FLPILASLAAKFGPKLFSLVTKKS | 75 | GFKRIVQKIRDFLRNLV | 35 | IKKILSKIKKLWK | 500 |
| GLNALKKFFQGIHEAIKLINNHVQ | 270 | AGTKEWLNKAKDFIKEKGLGMLSAAANAALN | 300 | GFWGKLWEGVKSAI | 50.1 |
| KWKLFKKIEKVGQGIGAVLKVLTTGL | 104.2 | ILPWKWCWWPCRR | 98.8 | KWKKLLKKLLKLL | 10 |
| FFRRFFRRFFRRFFRR | 10.5 | FLPIVGKLLSGLL | 75 | GLLSLLSALGKLL | 19 |
| FAKLLAKALKLKL | 371.1 | FLPAIVGAAAKFLPKIFCAISKKC | 14 | RVKRVWPLVIRTVIAGYNLYRAI | 38 |
| GWGSFFKKAAHVAKHVAKAALTHYL | 100 | GIGAVLAVLTTGLPALISWIKRKRQQ | 1 | GLNALKKVFQGIHEAIKLINNHVK | 380 |
| KWKLFKKIGIGAVLKVLTTGLPALKLTK | 16.3 | GKWMKMLKKILK | 200 | GIMSSLMKKLAAHIAK | 400 |
| GALRRLGRKITHAVKKYGPTVLRIIRIAG | 18.4 | ILCWKWPWCPWRR | 33.7 | YRAWRWAWRWR | 331.8 |
| GFKDLLKGAAKALKKTVLF | 500 | DSMGAVKLAKLLIDKMKCEVTKAC | 150 | IKIPAVVKDTLKKVAKGVLSAVAGALTQ | 300 |
| FSISPGKVLDKFGKIVGKVLKQLKKVSAVAKV | 28.3 | FLPVIAGVAANFLPKLFCAISKKC | 16 | GWKDLLKGAAKALVKTVF | 40 |
| GLFGKLIKKFGRKAISYAVKKARGKH | 6 | ALWKTMLKKLGTMALHAGK | 101 | WKKWWKKWWK | 1000 |
| FLSTIWNGIKGLL | 155.6 | GMATAAGTTLGKLAKFVIGAV | 160 | RVCIRVCYRGVCYRRCW | 128 |
| GLFDVIRRVASVIGGL | 10.9 | GIWKKWIKKVVNVLKNLF | 26 | GLLSALKALGKLL | 36 |
| GLNALKKVFKGIHEAIKLINNHVQ | 240 | VRRRRRPR | 87 | KRRKLIKILKLIIKLIRKKR | 8 |
| GCKKWFKKAAHVGKNVGKVALNAYL | 92.9 | FLPIVGRLISGLL | 210 | GIGAVLVVLTTGLPALISWIKRKRQQ | 175.9 |
| GLLKFIKKLL | 13.57 | GLLSNVAGLLKQFAKGGVNAVLNPK | 256 | GMATKAGTALGKVAKAVIGAAL | 150 |
| GLWSKIKDAAKTAGKAALGFVNEMV | 210 | FLSLIPKIAGGIAALAKHL | 22.8 | GMASKAGAIAGKIAKVALKAL | 0.6 |
| GLLRASSKWGRKYYVDLAGCAKA | 670 | YLRLIRYMAKMI | 78.2 | GKILKYLLYLLRKYANLIIR | 128 |
| FALAKKALKKAKKAL | 604.6 | GKWQSLLKHILK | 200 | GFWGKLLEGVKKAI | 1000 |
| LNWGKILKHIIK | 140 | FLPLLLAGLPSFLCLVFKKC | 439 | GVVVRVGRVVVRWV | 43 |
| GKLSSLWKKLKKIIAK | 400 | MDFIIDIIKKIVGLFTGK | 6.34 | GMWSKLLGHLLR | 62 |
| GLMKRIKTML | 500 | RVKRVWPLVIRTVIA | 443 | GLSLLLSLKLKLL | 300 |
| AKVVKKLTKGVAKLLK | 128 | INWKKIFQKVKNLV | 100 | IKPFKFPFKPFRR | 115.4 |
| GLFDIWKKWRWRR | 133.5 | ILPWKWPWWPWRR | 131.5 | KLKKLLKKLLKK | 332 |
| ILGKIIKVVK | 200 | QLLKKIRTWYRKAWHVYGKV | 200 | NLLKSALKTVNKLLAAS | 256 |
| GLLKRIKILL | 99.8 | FLSLALAALPKLFCLIFKKC | 4.7 | QIFKKVKTWYKKAFQILGRL | 101 |
| FKAFKAFKAFKAFKAFKA | 236.9 | KWKVFKKIEKMARNIRNKIVK | 250 | RKDVYRRRRRR | 609 |
| GLFDKWAWWRWRR | 132.4 | KIAGKIAKIAGKIAKIAGKIA | 20.1 | RLMRIFRILKLAR | 17 |
| FAKLLAKLAK | 886.5 | FAKLLAKLAKKAL | 173.6 | GFIFHIIKGLFHAGKMIHGLV | 84.7 |
| QLIHKIRTWYRKAWHVLGKV | 200 | FLSLIPKAISAISALANHF | 34.86 | IIPLPLGYFAKKT | 300 |
| FLGALWNVAKRVF | 12.5 | FFHHIFRAIVHVAKTIHRLVTG | 6 | FLGALWNVAKSLF | 12.5 |
| KAIAKSILKWIKSIAKAI | 1.8 | IKKIVSKIKKVLK | 384 | KLKLKLKLKLKLKLK | 1.1 |
| AILTTLANWARKFL | 75 | GLMDTIKGVAKNVAASLLEKLKCKVTGC | 120 | AAKKGCWTVSIPPKPCF | 1000 |

| | | | | | |
|---|---|---|---|---|---|
| GILSTFKGLAKGVAKDLAGNLLDKFKC KITGC | 120 | GIKDWIKGAAKKLIKTVASHIANQ | 200 | FLKGIVGMLGKLW | 32 |

#### Supplementary Table S2: The entire set of motifs found in hemolytic and non-hemolytic peptides.

| Postitive Motifs | Postitive Motifs | Negative Motifs | Negative Motifs |
|---|---|---|---|
| CGET | AGKAIHRLI | AKD | PTK |
| CGETC | AGKAIHRLIR | SKIK | QGIHE |
| ETC | AGKIAKIAGKIAKI | DLA | QGIHEA |
| GET | AGKIAKIAGKIAKIA | SKIKK | RWKK |
| GETC | AGKIAKIAGKIAKIAG | HVQ | SKIKKLL |
| TLLKK | AGKIAKIAGKIAKIAGK | NKL | TKG |
| TLLKKV | AGKIAKIAGKIAKIAGKI | HRK | TKWWK |
| TLLKKVL | AGKIAKIAGKIAKIAGKIA | INKQ | VAGLL |
| TLLKKVLK | AIHRL | KDLA | VAGLLK |
| TPG | AIHRLI | KINKQ | VFQGI |
| TPGC | AIHRLIR | NHV | VFQGIH |
| GGLF | ANFLPK | NKAK | VNKL |
| GGLFS | AVLVG | NKQ | WFQ |
| GLFS | AVLVGA | NNH | WFQN |
| TLLKKVLKA | AVLVGAN | NNHV | WGAI |
| AIH | AVLVGANA | RIKT | WKKILGK |
| IGGLF | AY | CKINKQ | AAANAAL |
| IGGLFS | CGETCF | CKINKQC | AAANAALN |
| IIGGL | CNTPG | EAI | AAKT |
| IIGGLF | CNTPGC | FQG | AANAAL |
| IIGGLFS | CRRRFCVCVG | GKIIK | AANAALN |
| RLIR | CRRRFCVCVGR | GLGSL | AGLG |
| ANF | CTC | HE | AGLGS |
| HHII | CTR | HEA | AGLGSL |
| HHIIG | CYCRRRFCVCVG | INKQC | AHR |
| HHIIGG | CYCRRRFCVCVGR | INNH | AHRK |
| HIIGG | ETCF | INNHV | AILKH |
| HRLI | FSAGKAIHR | KHII | AKDFIKEKGL |
| HRLIR | GANA | KHIIK | AKDLAG |
| SAGKA | GETCF | KINKQC | AKGVAKD |
| AGKAI | GGLFSAGKAIHR | KLINN | AKGVAKDL |

| | | | |
|---|---|---|---|
| AGKAIH | GGTCN | KLINNH | AKGVAKDLA |
| FIHH | GGTCNT | KLINNHV | AKVV |
| FSAG | GGTCNTP | KRIKT | ALKKVFQGI |
| FSAGK | GGTCNTPG | LINN | ALKKVFQGIH |
| FSAGKA | GGTCNTPGC | LINNH | ANAAL |
| GGLFSA | GIKS | LINNHV | ANAALN |
| GGLFSAG | GKAIHRL | NHVQ | DFIKEKGL |
| GGLFSAGK | GKAIHRLI | NKQC | EAIKLINN |
| GGLFSAGKA | GKAIHRLIR | NNHVQ | EAIKLINNH |
| GKAIH | GKIAKIAGKIAKI | RRFW | EAIKLINNHV |
| GLFSA | GKIAKIAGKIAKIA | SKIKKL | EKGL |
| GLFSAG | GKIAKIAGKIAKIAG | VAKGV | EW |
| GLFSAGK | GKIAKIAGKIAKIAGK | WRRF | EWL |
| GLFSAGKA | GKIAKIAGKIAKIAGKI | WRRFW | EWLN |
| HHIIGGL | GKIAKIAGKIAKIAGKIA | AAAN | EWLNK |
| HHIIGGLF | GLFSAGKAIHR | AAANA | EWLNKA |
| HHIIGGLFS | GTCN | AAANAA | EWLNKAK |
| HIIGGL | GTCNT | AANA | EWLNKAKD |
| HIIGGLF | GTCNTP | AANAA | EWLNKAKDF |
| HIIGGLFS | GTCNTPG | ACKINKQ | EWLNKAKDFI |
| HRLIRR | GTCNTPGC | ACKINKQC | EWLNKAKDFIK |
| HRLIRRR | GVAA | AIKLINN | EWLNKAKDFIKE |
| IHH | HHIIGGLFSA | AIKLINNH | EWLNKAKDFIKEK |
| IRRR | HHIIGGLFSAG | AIKLINNHV | EWLNKAKDFIKEKG |
| KAIH | HHIIGGLFSAGK | AKDF | FIKEKGL |
| KVLKAAAK | HHIIGGLFSAGKA | AKDFI | FKAGL |
| LFSA | HIIGGLFSA | AKDFIK | FKAGLG |
| LFSAG | HIIGGLFSAG | AKDFIKE | FKAGLGS |
| LFSAGK | HIIGGLFSAGK | AKDFIKEK | FKAGLGSL |
| LFSAGKA | HIIGGLFSAGKA | AKDFIKEKG | FQGIHEAI |
| LIRR | IAGKIAKIAGKIAKI | AKDL | FQGIHEAIK |
| LIRRR | IAGKIAKIAGKIAKIA | AKDLA | FQGIHEAIKL |
| RLIRR | IAGKIAKIAGKIAKIAG | ANAA | GAGK |
| RLIRRR | IAGKIAKIAGKIAKIAGK | DFIK | GAILKH |
| TLLKKVLKAA | IAGKIAKIAGKIAKIAGKI | DFIKE | GFVACKINKQ |
| TLLKKVLKAAA | IAGKIAKIAGKIAKIAGKIA | DFIKEK | GFVACKINKQC |
| VLKAAAK | IGGLFSAGKAI | DFIKEKG | GIAAH |
| CNT | IGGLFSAGKAIH | DLAG | GIAAHV |
| CNTP | IHRLIRRRR | EAIK | GIAAHVA |
| CVGR | IHRLIRRRRR | EAIKL | GIHEAIKLI |
| FIHHI | IIGGLFSAGKAI | EKG | GIHEAIKLIN |
| FIHHII | IIGGLFSAGKAIH | FIKE | GLGFVACKINKQ |
| FIHHIIG | ISAL | FIKEK | GLGFVACKINKQC |
| FIHHIIGG | KAAAKAAL | FIKEKG | GLGSLV |
| GLAA | KAIHRL | FQGI | GLGSLVK |
| HRLIRRRR | KAIHRLI | FQGIH | GLGSLVKG |
| HRLIRRRRR | KAIHRLIR | FVACKINKQ | GLGSLVKGI |

| | | | |
|---|---|---|---|
| IGGLFSA | KIAKIAGKIAKI | FVACKINKQC | GLNALKKV |
| IGGLFSAG | KIAKIAGKIAKIA | GIHE | GLNALKKVF |
| IGGLFSAGK | KIAKIAGKIAKIAG | GIHEA | GSLV |
| IGGLFSAGKA | KIAKIAGKIAKIAGK | GKIIKV | GSLVK |
| IHHI | KIAKIAGKIAKIAGKI | GLN | GSLVKG |
| IHHII | KIAKIAGKIAKIAGKIA | GLNA | GSLVKGI |
| IHHIIG | KVLKAAAKA | HEAI | GTLVNL |
| IHHIIGG | KVLKAAAKAA | HEAIK | GVAKD |
| IHRLI | KVLKAAAKAAL | HEAIKL | GVAKDL |
| IHRLIR | LAAN | IHE | GVAKDLA |
| IIGGLFSA | LFSAGKAIHR | IHEA | GVAKGV |
| IIGGLFSAG | LKAAAKAA | IKEK | GVAKGVA |
| IIGGLFSAGK | LKAAAKAAL | IKEKG | GVAKGVAK |
| IIGGLFSAGKA | LPKIFC | IKLINN | HEAIKLINN |
| IRRRR | LPT | IKLINNH | HEAIKLINNH |
| IRRRRR | LVGAN | IKLINNHV | HEAIKLINNHV |
| KKVLKAAAK | LVGANA | ILKHI | HRA |
| LIRRRR | NFLPK | INNHVQ | HRAH |
| LIRRRRR | NTPG | KAKD | HRAHR |
| LKKVLKAAAK | NTPGC | KAKDF | HRAHRK |
| LLKKVLKAAAK | PLA | KAKDFI | IAAH |
| NTP | RRFCVCVG | KAKDFIK | IAAHV |
| PKIF | RRFCVCVGR | KAKDFIKE | IAAHVA |
| RLIRRRR | RRRFCVCVG | KAKDFIKEK | IFKAG |
| RLIRRRRR | RRRFCVCVGR | KAKDFIKEKG | IFKAGL |
| SAGKAI | SAGKAIHRL | KDFIK | IFKAGLG |
| SAGKAIH | SAGKAIHRLI | KDFIKE | IFKAGLGS |
| TCN | SAGKAIHRLIR | KDFIKEK | IFKAGLGSL |
| TLLKKVLKAAAK | SLLGKL | KDFIKEKG | IHEAIKLI |
| AAAKAAL | TCF | KEKG | IHEAIKLIN |
| AAKAAL | TCG | KKVFQ | IIKVV |
| AANF | TCGE | KKVFQG | IIKVVK |
| AGKAIHR | TCGET | KLAKA | IKEKGL |
| AGLAA | TCGETC | KLINNHVQ | ILGKII |
| AIHR | TCNT | KVFQ | ILGKIIK |
| ANFL | TCNTP | KVFQG | ILGKIIKV |
| ANFLP | TCNTPG | LINNHVQ | ILSK |
| CVCVG | TCNTPGC | LKHII | KAGLG |
| CVCVGR | VGAN | LKHIIK | KAGLGS |
| CYT | VGANA | LNKAK | KAGLGSL |
| FCVCVG | VLKAAAKA | LNKAKD | KAKDFIKEKGL |
| FCVCVGR | VLKAAAKAA | LNKAKDF | KDFIKEKGL |
| FIHHIIGGL | VLKAAAKAAL | LNKAKDFI | KDLAG |
| FIHHIIGGLF | VLVG | LNKAKDFIK | KEKGL |
| FIHHIIGGLFS | VLVGA | LNKAKDFIKE | KEW |
| FSAGKAI | VLVGAN | LNKAKDFIKEK | KEWL |
| FSAGKAIH | VLVGANA | LNKAKDFIKEKG | KEWLN |

| | | | |
|---|---|---|---|
| GALA | WGAL | LPHV | KEWLNK |
| GGLFSAGKAI | YCRRRFCVCVG | LTG | KEWLNKA |
| GGLFSAGKAIH | YCRRRFCVCVGR | NKAKD | KEWLNKAK |
| GGRL | AAKF | NKAKDF | KEWLNKAKD |
| GGRLC | AALA | NKAKDFI | KEWLNKAKDF |
| GGRLCY | AGKAIHRLIRR | NKAKDFIK | KEWLNKAKDFI |
| GGRLCYC | AGKAIHRLIRRR | NKAKDFIKE | KEWLNKAKDFIK |
| GGRLCYCR | AIHRLIRR | NKAKDFIKEK | KEWLNKAKDFIKE |
| GGT | AIHRLIRRR | NKAKDFIKEKG | KEWLNKAKDFIKEK |
| GGTC | AISAL | PHV | KEWLNKAKDFIKEKG |
| GKAIHR | ALAKG | QGIH | KGIA |
| GLFSAGKAI | ALWKTLLKK | TWY | KGIAA |
| GLFSAGKAIH | ALWKTLLKKV | VACKINKQ | KGIAAH |
| GLPV | ALWN | VACKINKQC | KGIAAHV |
| GLPVC | ALWNV | VFQ | KGIAAHVA |
| GLPVCG | CTRN | VFQG | KGVAKD |
| GLPVCGE | FCF | WLNK | KGVAKDL |
| GLPVCGET | FLGALW | WLNKA | KGVAKDLA |
| GLPVCGETC | FLPKIF | WLNKAK | KGVAKG |
| GRLC | FLPKIFC | WLNKAKD | KGVAKGV |
| GRLCY | FSAGKAIHRL | WLNKAKDF | KGVAKGVA |
| GRLCYC | FSAGKAIHRLI | WLNKAKDFI | KGVAKGVAK |
| GRLCYCR | FSAGKAIHRLIR | WLNKAKDFIK | KIIKVV |
| IFC | FWGA | WLNKAKDFIKE | KIIKVVK |
| IHHIIGGL | GALAK | WLNKAKDFIKEK | KIKKLLK |
| IHHIIGGLF | GALAKG | WLNKAKDFIKEKG | KILGKI |
| IHHIIGGLFS | GGLFSAGKAIHRL | WSKIK | KKIIAK |
| IHRLIRR | GGLFSAGKAIHRLI | AIKLINNHVQ | KKILGKI |
| IHRLIRRR | GGLFSAGKAIHRLIR | ALKKVFQ | KKLKKI |
| KAIHR | GGRLCYCRR | ALKKVFQG | KKVAK |
| KIFC | GGRLCYCRRR | EAIKLI | KKVFQGIHE |
| KKD | GGRLCYCRRRF | EAIKLIN | KKVFQGIHEA |
| LFSAGKAI | GGRLCYCRRRFC | FKAG | KRWK |
| LFSAGKAIH | GGRLCYCRRRFCV | FQGIHE | KRWKK |
| LLKLL | GGRLCYCRRRFCVC | FQGIHEA | KVAA |
| LLSA | GKAIHRLIRR | FQN | KVFQGIHE |
| LPKIF | GKAIHRLIRRR | GIHEAI | KVFQGIHEA |
| LPVC | GLFSAGKAIHRL | GIHEAIK | LGFVACKINKQ |
| LPVCG | GLFSAGKAIHRLI | GIHEAIKL | LGFVACKINKQC |
| LPVCGE | GLFSAGKAIHRLIR | GLNAL | LGML |
| LPVCGET | GLPT | GLNALK | LGSLV |
| LPVCGETC | GRLCYCRR | GLNALKK | LGSLVK |
| LVGA | GRLCYCRRR | GTLVN | LGSLVKG |
| NFLP | GRLCYCRRRF | HEAIKLI | LGSLVKGI |
| PKIFC | GRLCYCRRRFC | HEAIKLIN | LGTLVN |
| PVCG | GRLCYCRRRFCV | IHEAI | LKKVFQGI |
| PVCGE | GRLCYCRRRFCVC | IHEAIK | LKKVFQGIH |

| | | | |
|---|---|---|---|
| PVCGET | IGC | IHEAIKL | LKVA |
| PVCGETC | IGCS | IKLINNHVQ | LKVAA |
| RFCVCVG | IGCSC | ILKHII | LLKKIL |
| RFCVCVGR | IGGLFSAGKAIHR | ILKHIIK | LLKKILK |
| RGGRL | IIGGLFSAGKAIHR | ILKV | LNALKKV |
| RGGRLC | ILA | IWF | LNALKKVF |
| RGGRLCY | KAIHRLIRR | IWFQ | LNKAKDFIKEKGL |
| RGGRLCYC | KAIHRLIRRR | IWFQN | LSAAA |
| RGGRLCYCR | KKVLKAAAKA | KKIIA | LSAAAN |
| RLC | KKVLKAAAKAA | KKLFK | LSAAANA |
| RLCY | KKVLKAAAKAAL | KKLFKK | LSAAANAA |
| RLCYC | KTLLKK | KKVFQGI | LSKIK |
| RLCYCR | KTLLKKV | KKVFQGIH | LSKIKK |
| SAGKAIHR | LAANF | KPF | LVKGI |
| SY | LAGLA | KVFQGI | NAAL |
| TPGCS | LAGLAA | KVFQGIH | NAALN |
| TPGCSC | LFR | LGKII | NALKKV |
| VCGE | LFSAGKAIHRL | LGKIIK | NALKKVF |
| VCGET | LFSAGKAIHRLI | LGKIIKV | NKAKDFIKEKGL |
| VCGETC | LFSAGKAIHRLIR | LKKVFQ | NKLT |
| VCVG | LGALW | LKKVFQG | NWGAI |
| VCVGR | LKKVLKAAAKA | LNAL | PFK |
| YT | LKKVLKAAAKAA | LNALK | PFR |
| AANFL | LKKVLKAAAKAAL | LNALKK | PFRR |
| AANFLP | LLKKVLKAAAKA | LVKG | PTKW |
| AANFLPK | LLKKVLKAAAKAA | NALK | QGIHEAI |
| AGKAIHRL | LLKKVLKAAAKAAL | NALKK | QGIHEAIK |
| RGGRLCYCRR | LLSAL | VAKD | QGIHEAIKL |
| RGGRLCYCRRR | LLSLL | VAKDL | RAH |
| RGGRLCYCRRRF | LWKTLLKK | VAKDLA | RAHR |
| RGGRLCYCRRRFC | LWKTLLKKV | VAKGVA | RAHRK |
| RGGRLCYCRRRFCV | LWNV | VAKGVAK | RIKTL |
| RGGRLCYCRRRFCVC | MTL | VFQGIHE | RTWY |
| RLCYCRR | PHI | VFQGIHEA | RWKKW |
| RLCYCRRR | PTC | VGLGFVACKINKQ | SAAA |
| RLCYCRRRF | PTCG | VGLGFVACKINKQC | SAAAN |
| RLCYCRRRFC | PTCGE | VKGI | SAAANA |
| RLCYCRRRFCV | PTCGET | VNKLT | SAAANAA |
| RLCYCRRRFCVC | PTCGETC | VVNK | SKIKKLLK |
| SAGKAIHRLIRR | TLLKKVLKAAAKAA | VVNKL | SLVKG |
| SAGKAIHRLIRRR | TLLKKVLKAAAKAAL | VVNKLT | SLVKGI |
| SLFS | TRN | WGAIL | TKWWKG |
| SLLGKLL | WKTLLKK | WGAILK | TWPT |
| TLLKKVLKAAAKA | WKTLLKKV | WKKILGKI | TWPTK |
| | WNV | WKKVL | TWPTKW |
| | | WLNKAKDFIKEKGL | TWYR |
| | | WPT | TWYRK |

| | |
|---|---|
| WPTK | TWYRKA |
| WPTKW | WYRK |
| WYR | WYRKA |
| | YRKA |

#### Supplementary Table S3: A detailed correlation analysis of each feature with the $HC_{50}$ value.

| Features | Correlation | Features | Correlation | Features | Correlation | Features | Correlation |
|---|---|---|---|---|---|---|---|
| **Molecular Weight (kDa)** | **-0.201** | **CTC_211** | **-0.148** | **APAAC1_V** | **-0.018** | **PAAC1_S** | **-0.092** |
| **length** | **-0.201** | **CTC_212** | **-0.074** | **APAAC1_W** | **0.018** | **PAAC1_T** | **-0.039** |
| **AAC_A** | **0.011** | **CTC_213** | **-0.047** | **APAAC1_Y** | **0.010** | **PAAC1_V** | **-0.018** |
| **AAC_C** | **-0.075** | **CTC_214** | **0.028** | **APAAC1_HB_lam1** | **-0.032** | **PAAC1_W** | **0.018** |
| **AAC_D** | **-0.030** | **CTC_215** | **0.027** | **APAAC1_HL_lam1** | **0.019** | **PAAC1_Y** | **0.010** |
| **AAC_E** | **-0.043** | **CTC_216** | **-0.013** | **APAAC1_SC_lam1** | **-0.010** | **PAAC1_lam1** | **0.116** |
| **AAC_F** | **-0.093** | **CTC_217** | **-0.065** | **QSO1_SC_A** | **-0.041** | **APAAC1_A** | **0.011** |
| **AAC_G** | **-0.102** | **CTC_221** | **-0.118** | **QSO1_SC_C** | **-0.100** | **APAAC1_C** | **-0.075** |
| **AAC_H** | **-0.036** | **CTC_222** | **-0.094** | **QSO1_SC_D** | **-0.031** | **APAAC1_D** | **-0.030** |
| **AAC_I** | **-0.007** | **CTC_223** | **-0.094** | **QSO1_SC_E** | **-0.049** | **APAAC1_E** | **-0.043** |
| **AAC_K** | **0.214** | **CTC_224** | **-0.068** | **QSO1_SC_F** | **-0.144** | **APAAC1_F** | **-0.093** |
| **AAC_L** | **-0.020** | **CTC_225** | **-0.006** | **QSO1_SC_G** | **-0.142** | **APAAC1_G** | **-0.102** |
| **AAC_M** | **-0.013** | **CTC_226** | **-0.051** | **QSO1_SC_H** | **-0.052** | **APAAC1_H** | **-0.036** |
| **AAC_N** | **-0.043** | **CTC_227** | **-0.076** | **QSO1_SC_I** | **-0.077** | **APAAC1_I** | **-0.007** |
| **AAC_P** | **-0.075** | **CTC_231** | **-0.093** | **QSO1_SC_K** | **0.121** | **APAAC1_K** | **0.214** |
| **AAC_Q** | **-0.026** | **CTC_232** | **-0.072** | **QSO1_SC_L** | **-0.099** | **APAAC1_L** | **-0.020** |
| **AAC_R** | **0.043** | **CTC_233** | **-0.049** | **QSO1_SC_M** | **-0.031** | **APAAC1_M** | **-0.013** |
| **AAC_S** | **-0.092** | **CTC_234** | **-0.039** | **QSO1_SC_N** | **-0.056** | **APAAC1_N** | **-0.043** |
| **AAC_T** | **-0.039** | **CTC_235** | **0.048** | **QSO1_SC_P** | **-0.112** | **APAAC1_P** | **-0.075** |
| **AAC_V** | **-0.018** | **CTC_236** | **-0.024** | **QSO1_SC_Q** | **-0.040** | **APAAC1_Q** | **-0.026** |
| **AAC_W** | **0.018** | **CTC_237** | **-0.026** | **QSO1_SC_R** | **0.006** | **APAAC1_R** | **0.043** |
| **AAC_Y** | **0.010** | **CTC_241** | **-0.035** | **QSO1_SC_S** | **-0.116** | **APAAC1_S** | **-0.092** |
| **DPC1_AA** | **-0.009** | **CTC_242** | **-0.049** | **QSO1_SC_T** | **-0.089** | **APAAC1_T** | **-0.039** |
| **DPC1_AC** | **-0.012** | **CTC_243** | **0.008** | **QSO1_SC_V** | **-0.055** | **RRI_L** | **-0.028** |

| | | | | | | | |
|---|---|---|---|---|---|---|---|
| **DPC1_AD** | **-0.021** | **CTC_244** | **-0.084** | **QSO1_SC_W** | **-0.024** | **RRI_M** | **-0.053** |
| **DPC1_AE** | **0.030** | **CTC_245** | **-0.089** | **QSO1_SC_Y** | **-0.014** | **RRI_N** | **-0.070** |
| **DPC1_AF** | **0.012** | **CTC_246** | **-0.025** | **QSO1_G_A** | **0.019** | **RRI_P** | **-0.115** |
| **DPC1_AG** | **-0.085** | **CTC_247** | **-0.016** | **QSO1_G_C** | **-0.075** | **RRI_Q** | **-0.052** |
| **DPC1_AH** | **0.035** | **CTC_251** | **-0.029** | **QSO1_G_D** | **0.020** | **RRI_R** | **-0.032** |
| **DPC1_AI** | **-0.087** | **CTC_252** | **-0.090** | **QSO1_G_E** | **-0.022** | **RRI_S** | **-0.080** |
| **DPC1_AK** | **0.165** | **CTC_253** | **0.032** | **QSO1_G_F** | **-0.057** | **RRI_T** | **-0.081** |
| **DPC1_AL** | **-0.045** | **CTC_254** | **-0.037** | **QSO1_G_G** | **-0.080** | **RRI_V** | **-0.070** |
| **DPC1_AM** | **0.000** | **CTC_255** | **0.134** | **QSO1_G_H** | **-0.027** | **RRI_W** | **-0.084** |
| **DPC1_AN** | **-0.001** | **CTC_256** | **0.039** | **QSO1_G_I** | **0.064** | **RRI_Y** | **-0.073** |
| **DPC1_AP** | **0.001** | **CTC_257** | **-0.044** | **QSO1_G_K** | **0.213** | **PRI_PC** | **-0.050** |
| **DPC1_AQ** | **0.010** | **CTC_261** | **-0.043** | **QSO1_G_L** | **0.020** | **PRI_NC** | **-0.047** |
| **DPC1_AR** | **-0.031** | **CTC_262** | **-0.035** | **QSO1_G_M** | **0.015** | **PRI_NE** | **-0.008** |
| **DPC1_AS** | **-0.044** | **CTC_263** | **0.006** | **QSO1_G_N** | **-0.019** | **PRI_PO** | **0.020** |
| **DPC1_AT** | **-0.023** | **CTC_264** | **-0.040** | **QSO1_G_P** | **-0.052** | **PRI_NP** | **0.076** |
| **DPC1_AV** | **-0.010** | **CTC_265** | **-0.032** | **QSO1_G_Q** | **-0.007** | **PRI_AL** | **0.053** |
| **DPC1_AW** | **-0.011** | **CTC_266** | **-0.001** | **QSO1_G_R** | **0.085** | **PRI_CY** | **-0.100** |
| **DPC1_AY** | **-0.033** | **CTC_267** | **-0.018** | **QSO1_G_S** | **-0.057** | **PRI_AR** | **-0.058** |
| **DPC1_CA** | **-0.021** | **CTC_271** | **-0.069** | **QSO1_G_T** | **0.015** | **PRI_AC** | **-0.047** |
| **DPC1_CC** | **0.009** | **CTC_272** | **-0.039** | **QSO1_G_V** | **0.010** | **PRI_BS** | **-0.050** |
| **DPC1_CD** | **-0.023** | **CTC_273** | **-0.055** | **QSO1_G_W** | **0.014** | **PRI_NE_pH** | **-0.008** |
| **DPC1_CE** | **-0.014** | **CTC_274** | **-0.038** | **QSO1_G_Y** | **0.093** | **PRI_HB** | **0.112** |
| **DPC1_CF** | **-0.066** | **CTC_275** | **-0.014** | **QSO1_SC1** | **-0.090** | **PRI_HL** | **0.006** |
| **DPC1_CG** | **0.002** | **CTC_276** | **-0.018** | **QSO1_G1** | **-0.090** | **PRI_NT** | **-0.003** |
| **DPC1_CH** | **0.005** | **CTC_277** | **-0.016** | **SEP_PC** | **0.203** | **PRI_HX** | **-0.015** |
| **DPC1_CI** | **-0.042** | **CTC_311** | **-0.038** | **SEP_NC** | **-0.054** | **PRI_SC** | **-0.072** |
| **DPC1_CK** | **-0.023** | **CTC_312** | **-0.115** | **SEP_NE** | **0.195** | **PRI_SS_HE** | **0.206** |
| **DPC1_CL** | **0.036** | **CTC_313** | **0.004** | **SEP_PO** | **-0.102** | **PRI_SS_ST** | **0.160** |
| **DPC1_CM** | **-0.018** | **CTC_314** | **-0.012** | **SEP_NP** | **0.064** | **PRI_SS_CO** | **0.000** |
| **DPC1_CN** | **-0.063** | **CTC_315** | **0.016** | **SEP_AL** | **-0.060** | **PRI_SA_BU** | **0.085** |
| **DPC1_CP** | **-0.002** | **CTC_316** | **0.034** | **SEP_CY** | **-0.093** | **PRI_SA_EX** | **-0.025** |
| **DPC1_CQ** | **-0.021** | **CTC_317** | **-0.037** | **SEP_AR** | **-0.099** | **PRI_SA_IN** | **-0.016** |
| **DPC1_CR** | **-0.002** | **CTC_321** | **-0.019** | **SEP_AC** | **-0.054** | **PRI_TN** | **-0.014** |
| **DPC1_CS** | **-0.055** | **CTC_322** | **-0.093** | **SEP_BS** | **0.203** | **PRI_SM** | **0.054** |
| **DPC1_CT** | **-0.035** | **CTC_323** | **0.000** | **SEP_NE_pH** | **0.195** | **PRI_LR** | **0.176** |
| **DPC1_CV** | **-0.043** | **CTC_324** | **-0.064** | **SEP_HB** | **0.006** | **DDR_A** | **-0.090** |
| **DPC1_CW** | **-0.036** | **CTC_325** | **0.012** | **SEP_HL** | **0.184** | **DDR_C** | **-0.072** |
| **DPC1_CY** | **-0.083** | **CTC_326** | **-0.015** | **SEP_NT** | **-0.136** | **DDR_D** | **-0.038** |
| **DPC1_DA** | **-0.028** | **CTC_327** | **-0.029** | **SEP_HX** | **-0.095** | **DDR_E** | **-0.063** |
| **DPC1_DC** | **0.046** | **CTC_331** | **-0.073** | **SEP_SC** | **-0.088** | **DDR_F** | **-0.128** |
| **DPC1_DD** | **-** | **CTC_332** | **-0.067** | **SEP_SS_HE** | **-0.080** | **DDR_G** | **-0.135** |
| **DPC1_DE** | **-0.001** | **CTC_333** | **0.029** | **SEP_SS_ST** | **-0.075** | **DDR_H** | **-0.057** |

| | | | | | | | |
|---|---|---|---|---|---|---|---|
| DPC1_DF | -0.011 | CTC_334 | -0.033 | SEP_SS_CO | -0.162 | DDR_I | -0.110 |
| DPC1_DG | -0.031 | CTC_335 | 0.001 | SEP_SA_BU | 0.034 | DDR_K | -0.132 |
| DPC1_DH | -0.024 | CTC_336 | 0.020 | SEP_SA_EX | 0.122 | DDR_L | -0.096 |
| DPC1_DI | -0.037 | CTC_337 | -0.013 | SEP_SA_IN | -0.120 | DDR_M | -0.066 |
| DPC1_DK | -0.030 | CTC_341 | -0.022 | SEP_TN | -0.123 | DDR_N | -0.084 |
| DPC1_DL | 0.052 | CTC_342 | -0.042 | SEP_SM | -0.136 | DDR_P | -0.122 |
| DPC1_DM | 0.019 | CTC_343 | 0.001 | SEP_LR | -0.136 | DDR_Q | -0.062 |
| DPC1_DN | 0.001 | CTC_344 | -0.018 | SOC1_SC1 | 0.135 | DDR_R | -0.103 |
| DPC1_DP | -0.024 | CTC_345 | 0.019 | SOC1_G1 | -0.128 | CeTD_75_p_HB1 | 0.025 |
| DPC1_DQ | -0.027 | CTC_346 | 0.000 | DDR_S | -0.072 | CeTD_100_p_HB1 | 0.013 |
| DPC1_DR | -0.006 | CTC_347 | -0.017 | DDR_T | -0.102 | CeTD_0_p_VW1 | - |
| DPC1_DS | -0.013 | CTC_351 | -0.010 | DDR_V | -0.081 | CeTD_25_p_VW1 | -0.154 |
| DPC1_DT | 0.004 | CTC_352 | 0.056 | DDR_W | -0.152 | CeTD_50_p_VW1 | -0.150 |
| DPC1_DV | -0.008 | CTC_353 | -0.008 | DDR_Y | -0.115 | CeTD_75_p_VW1 | -0.154 |
| DPC1_DW | -0.016 | CTC_354 | 0.007 | SER_A | 0.026 | CeTD_100_p_VW1 | -0.158 |
| DPC1_DY | 0.000 | CTC_355 | 0.012 | SER_C | 0.086 | CeTD_0_p_PO1 | - |
| DPC1_EA | 0.053 | CTC_356 | -0.015 | SER_D | 0.030 | CeTD_25_p_PO1 | -0.298 |
| DPC1_EC | -0.030 | CTC_357 | -0.011 | SER_E | 0.049 | CeTD_50_p_PO1 | -0.318 |
| DPC1_ED | -0.009 | CTC_361 | -0.013 | SER_F | 0.114 | CeTD_75_p_PO1 | -0.327 |
| DPC1_EE | -0.014 | CTC_362 | -0.022 | SER_G | 0.128 | CeTD_100_p_PO1 | -0.324 |
| DPC1_EF | -0.018 | CTC_363 | -0.020 | SER_H | 0.042 | CeTD_0_p_PZ1 | - |
| DPC1_EG | 0.023 | CTC_364 | -0.015 | SER_I | 0.071 | CeTD_25_p_PZ1 | -0.148 |
| DPC1_EH | -0.009 | CTC_365 | -0.011 | SER_K | -0.154 | CeTD_50_p_PZ1 | -0.148 |
| DPC1_EI | -0.024 | CTC_366 | -0.007 | SER_L | 0.025 | CeTD_75_p_PZ1 | -0.147 |
| DPC1_EK | -0.026 | CTC_367 | 0.085 | SER_M | 0.033 | CeTD_100_p_PZ1 | -0.150 |
| DPC1_EL | 0.050 | CTC_371 | -0.062 | SER_N | 0.055 | CeTD_0_p_CH1 | - |
| DPC1_EM | -0.010 | CTC_372 | -0.041 | SER_P | 0.099 | CeTD_25_p_CH1 | -0.145 |
| DPC1_EN | -0.015 | CTC_373 | -0.071 | SER_Q | 0.036 | CeTD_50_p_CH1 | -0.175 |
| DPC1_EP | -0.006 | CTC_374 | -0.058 | SER_R | 0.008 | CeTD_75_p_CH1 | -0.172 |
| DPC1_EQ | 0.002 | CTC_375 | -0.032 | SER_S | 0.089 | CeTD_100_p_CH1 | -0.173 |
| DPC1_ER | -0.008 | CTC_376 | -0.018 | SER_T | 0.054 | CeTD_0_p_SS1 | - |
| DPC1_ES | -0.024 | CTC_377 | - | SER_V | 0.050 | CeTD_25_p_SS1 | -0.046 |
| DPC1_ET | -0.052 | CTC_411 | -0.076 | SER_W | 0.046 | CeTD_50_p_SS1 | -0.048 |
| DPC1_EV | -0.037 | CTC_412 | -0.017 | SER_Y | 0.039 | CeTD_75_p_SS1 | -0.043 |
| DPC1_EW | -0.013 | CTC_413 | -0.027 | SEP | -0.199 | CeTD_100_p_SS1 | -0.043 |
| DPC1_EY | -0.038 | CTC_414 | 0.057 | CTC_111 | -0.035 | CeTD_0_p_SA1 | - |
| DPC1_FA | 0.088 | CTC_415 | 0.039 | CTC_112 | -0.112 | CeTD_25_p_SA1 | -0.282 |
| DPC1_FC | -0.086 | CTC_416 | -0.012 | CTC_113 | -0.029 | CeTD_50_p_SA1 | -0.307 |
| DPC1_FD | -0.045 | CTC_417 | -0.038 | CTC_114 | -0.046 | CeTD_75_p_SA1 | -0.316 |
| DPC1_FE | -0.039 | CTC_421 | -0.002 | CTC_115 | -0.048 | CeTD_100_p_SA1 | -0.315 |
| DPC1_FF | -0.039 | CTC_422 | -0.068 | CTC_116 | 0.066 | CeTD_0_p_HB2 | - |
| DPC1_FG | -0.041 | CTC_423 | -0.007 | CTC_117 | -0.001 | CeTD_25_p_HB2 | -0.142 |

| | | | | | | | |
|---|---|---|---|---|---|---|---|
| **DPC1_FH** | **-0.045** | **CTC_424** | **-0.042** | **CTC_121** | **-0.095** | **CeTD_50_p_HB2** | **-0.149** |
| **DPC1_FI** | **-0.069** | **CTC_425** | **0.002** | **CTC_122** | **-0.058** | **CeTD_75_p_HB2** | **-0.148** |
| **DPC1_FK** | **-0.008** | **CTC_426** | **-0.011** | **CTC_123** | **-0.072** | **CeTD_100_p_HB2** | **-0.153** |
| **DPC1_FL** | **-0.088** | **CTC_427** | **-0.027** | **CTC_124** | **-0.074** | **CeTD_0_p_VW2** | **-** |
| **DPC1_FM** | **-0.015** | **CTC_431** | **0.028** | **CTC_125** | **-0.055** | **CeTD_25_p_VW2** | **0.009** |
| **DPC1_FN** | **-0.012** | **CTC_432** | **-0.022** | **CTC_126** | **-0.040** | **CeTD_50_p_VW2** | **0.008** |
| **DPC1_FP** | **0.009** | **CTC_433** | **-0.051** | **CTC_127** | **-0.038** | **CeTD_75_p_VW2** | **0.006** |
| **DPC1_FQ** | **0.001** | **CTC_434** | **-0.020** | **CTC_131** | **-0.045** | **CeTD_100_p_VW2** | **-0.002** |
| **DPC1_FR** | **-0.045** | **CTC_435** | **-0.014** | **CTC_132** | **-0.027** | **CeTD_0_p_PO2** | **-** |
| **DPC1_FS** | **-0.057** | **CTC_436** | **-0.032** | **CTC_133** | **-0.027** | **CeTD_25_p_PO2** | **-0.133** |
| **DPC1_FT** | **-0.054** | **CTC_437** | **0.026** | **CTC_134** | **0.023** | **CeTD_50_p_PO2** | **-0.132** |
| **DPC1_FV** | **0.006** | **CTC_441** | **-0.025** | **CTC_135** | **0.022** | **CeTD_75_p_PO2** | **-0.131** |
| **DPC1_FW** | **0.048** | **CTC_442** | **-0.097** | **CTC_136** | **-0.001** | **CeTD_100_p_PO2** | **-0.132** |
| **DPC1_FY** | **-0.005** | **CTC_443** | **-0.038** | **CTC_137** | **-0.051** | **CeTD_0_p_PZ2** | **-** |
| **DPC1_GA** | **-0.061** | **CTC_444** | **0.005** | **CTC_141** | **-0.041** | **CeTD_25_p_PZ2** | **-0.198** |
| **DPC1_GC** | **-0.039** | **CTC_445** | **0.067** | **CTC_142** | **0.020** | **CeTD_50_p_PZ2** | **-0.212** |
| **DPC1_GD** | **0.002** | **CTC_446** | **0.006** | **CTC_143** | **-0.029** | **CeTD_75_p_PZ2** | **-0.209** |
| **DPC1_GE** | **-0.044** | **CTC_447** | **-0.014** | **CTC_144** | **-0.007** | **CeTD_100_p_PZ2** | **-0.207** |
| **DPC1_GF** | **-0.003** | **CTC_451** | **0.034** | **CTC_145** | **0.055** | **CeTD_0_p_CH2** | **-** |
| **DPC1_GG** | **-0.094** | **CTC_452** | **-0.034** | **CTC_146** | **-0.008** | **CeTD_25_p_CH2** | **-0.046** |
| **DPC1_GH** | **-0.007** | **CTC_453** | **-0.026** | **CTC_147** | **0.001** | **CeTD_50_p_CH2** | **-0.048** |
| **DPC1_GI** | **0.001** | **CTC_454** | **0.012** | **CTC_151** | **0.019** | **CeTD_75_p_CH2** | **-0.043** |
| **DPC1_GK** | **-0.076** | **CTC_455** | **0.014** | **CTC_152** | **0.011** | **CeTD_100_p_CH2** | **-0.043** |
| **DPC1_GL** | **-0.070** | **CTC_456** | **-0.021** | **CTC_153** | **-0.051** | **CeTD_0_p_SS2** | **-** |
| **DPC1_GM** | **0.008** | **CTC_457** | **0.068** | **CTC_154** | **-0.051** | **CeTD_25_p_SS2** | **0.046** |
| **DPC1_GN** | **-0.024** | **CTC_461** | **0.013** | **CTC_155** | **0.072** | **CeTD_50_p_SS2** | **0.059** |
| **DPC1_GP** | **-0.017** | **CTC_462** | **-0.014** | **CTC_156** | **-0.020** | **CeTD_75_p_SS2** | **0.063** |
| **DPC1_GQ** | **-0.024** | **CTC_463** | **-0.030** | **CTC_157** | **0.002** | **CeTD_100_p_SS2** | **0.053** |
| **DPC1_GR** | **-0.004** | **CTC_464** | **-0.019** | **CTC_161** | **0.003** | **CeTD_0_p_SA2** | **-** |
| **DPC1_GS** | **0.021** | **CTC_465** | **-0.010** | **CTC_162** | **0.027** | **CeTD_25_p_SA2** | **-0.254** |
| **DPC1_GT** | **-0.003** | **CTC_466** | **-0.006** | **CTC_163** | **-0.055** | **CeTD_50_p_SA2** | **-0.249** |
| **DPC1_GV** | **0.050** | **CTC_467** | **-0.004** | **CTC_164** | **-0.014** | **CeTD_75_p_SA2** | **-0.249** |
| **DPC1_GW** | **0.004** | **CTC_471** | **-0.028** | **CTC_165** | **-0.013** | **CeTD_100_p_SA2** | **-0.250** |
| **DPC1_GY** | **-0.028** | **CTC_472** | **-0.014** | **CTC_166** | **-0.015** | **CeTD_0_p_HB3** | **-** |
| **DPC1_HA** | **-0.041** | **CTC_473** | **-0.024** | **CTC_167** | **-0.025** | **CeTD_25_p_HB3** | **0.004** |
| **DPC1_HC** | **0.006** | **CTC_474** | **-0.031** | **CTC_171** | **-0.038** | **CeTD_50_p_HB3** | **-0.025** |
| **DPC1_HD** | **-** | **CTC_475** | **0.019** | **CTC_172** | **-0.036** | **CeTD_75_p_HB3** | **-0.019** |
| **DPC1_HE** | **0.043** | **CTC_476** | **-** | **CTC_173** | **-0.074** | **CeTD_100_p_HB3** | **-0.048** |
| **DPC1_HF** | **0.032** | **CTC_477** | **-0.016** | **CTC_174** | **0.048** | **CeTD_0_p_VW3** | **-** |
| **DPC1_HG** | **-0.028** | **CTC_511** | **0.007** | **CTC_175** | **0.000** | **CeTD_25_p_VW3** | **-0.064** |
| **DPC1_HH** | **-0.061** | **CTC_512** | **-0.060** | **CTC_176** | **-** | **CeTD_50_p_VW3** | **-0.064** |
| **DPC1_HI** | **-0.066** | **CTC_513** | **0.051** | **CTC_177** | **-** | **CeTD_75_p_VW3** | **-0.062** |

| | | | | | | | |
|---|---|---|---|---|---|---|---|
| DPC1_HK | -0.003 | CTC_514 | 0.041 | CeTD_50_p_CH3 | -0.250 | CeTD_100_p_VW3 | -0.066 |
| DPC1_HL | -0.017 | CTC_515 | 0.016 | CeTD_75_p_CH3 | -0.252 | CeTD_0_p_PO3 | - |
| DPC1_HM | -0.033 | CTC_516 | -0.022 | CeTD_100_p_CH3 | -0.256 | CeTD_25_p_PO3 | -0.188 |
| DPC1_HN | -0.017 | CTC_517 | -0.012 | CeTD_0_p_SS3 | - | CeTD_50_p_PO3 | -0.190 |
| DPC1_HP | - | CTC_521 | -0.016 | CeTD_25_p_SS3 | 0.028 | CeTD_75_p_PO3 | -0.196 |
| DPC1_HQ | - | CTC_522 | -0.005 | CeTD_50_p_SS3 | 0.025 | CeTD_100_p_PO3 | -0.195 |
| DPC1_HR | 0.038 | CTC_523 | 0.041 | CeTD_75_p_SS3 | 0.025 | CeTD_0_p_PZ3 | - |
| DPC1_HS | 0.005 | CTC_524 | 0.022 | CeTD_100_p_SS3 | 0.013 | CeTD_25_p_PZ3 | -0.145 |
| DPC1_HT | -0.006 | CTC_525 | 0.108 | CeTD_0_p_SA3 | - | CeTD_50_p_PZ3 | -0.161 |
| DPC1_HV | -0.010 | CTC_526 | 0.005 | CeTD_25_p_SA3 | -0.130 | CeTD_75_p_PZ3 | -0.161 |
| DPC1_HW | 0.033 | CTC_527 | -0.047 | CeTD_50_p_SA3 | -0.151 | CeTD_100_p_PZ3 | -0.162 |
| DPC1_HY | -0.028 | CTC_531 | 0.005 | CeTD_75_p_SA3 | -0.154 | CeTD_0_p_CH3 | - |
| DPC1_IA | -0.037 | CTC_532 | -0.008 | CeTD_100_p_SA3 | -0.158 | CeTD_25_p_CH3 | -0.229 |
| DPC1_IC | -0.068 | CTC_533 | -0.033 | PAAC1_A | 0.011 | DPC1_WP | -0.029 |
| DPC1_ID | -0.021 | CTC_534 | -0.005 | PAAC1_C | -0.075 | DPC1_WQ | -0.013 |
| DPC1_IE | -0.007 | CTC_535 | -0.022 | PAAC1_D | -0.030 | DPC1_WR | 0.047 |
| DPC1_IF | -0.060 | CTC_536 | -0.032 | PAAC1_E | -0.043 | DPC1_WS | 0.002 |
| DPC1_IG | -0.132 | CTC_537 | -0.008 | PAAC1_F | -0.093 | DPC1_WT | 0.019 |
| DPC1_IH | -0.073 | CTC_541 | 0.020 | PAAC1_G | -0.102 | DPC1_WV | -0.008 |
| DPC1_II | 0.094 | CTC_542 | -0.059 | PAAC1_H | -0.036 | DPC1_WW | 0.051 |
| DPC1_IK | 0.191 | CTC_543 | -0.015 | PAAC1_I | -0.007 | DPC1_WY | -0.030 |
| DPC1_IL | -0.038 | CTC_544 | 0.054 | PAAC1_K | 0.214 | DPC1_YA | -0.044 |
| DPC1_IM | 0.020 | CTC_545 | 0.020 | PAAC1_L | -0.020 | DPC1_YC | -0.055 |
| DPC1_IN | -0.007 | CTC_546 | -0.021 | PAAC1_M | -0.013 | DPC1_YD | 0.085 |
| DPC1_IP | -0.024 | CTC_547 | -0.037 | PAAC1_N | -0.043 | DPC1_YE | -0.009 |
| DPC1_IQ | -0.034 | CTC_551 | 0.031 | PAAC1_P | -0.075 | DPC1_YF | -0.018 |
| DPC1_IR | -0.004 | CTC_552 | 0.150 | PAAC1_Q | -0.026 | DPC1_YG | 0.084 |
| DPC1_IS | -0.065 | CTC_553 | -0.008 | PAAC1_R | 0.043 | DPC1_YH | -0.006 |
| DPC1_IT | -0.069 | CTC_554 | 0.062 | CTC_743 | -0.014 | DPC1_YI | -0.016 |
| DPC1_IV | 0.020 | CTC_555 | 0.000 | CTC_744 | -0.036 | DPC1_YK | -0.053 |
| DPC1_IW | -0.042 | CTC_556 | -0.039 | CTC_745 | -0.029 | DPC1_YL | -0.054 |
| DPC1_IY | -0.022 | CTC_557 | -0.093 | CTC_746 | - | DPC1_YM | -0.018 |
| DPC1_KA | 0.047 | CTC_561 | -0.014 | CTC_747 | -0.019 | DPC1_YN | -0.026 |
| DPC1_KC | -0.038 | CTC_562 | 0.012 | CTC_751 | -0.015 | DPC1_YP | -0.015 |
| DPC1_KD | 0.048 | CTC_563 | 0.013 | CTC_752 | 0.008 | DPC1_YQ | 0.002 |
| DPC1_KE | -0.039 | CTC_564 | 0.010 | CTC_753 | -0.013 | DPC1_YR | 0.101 |
| DPC1_KF | -0.042 | CTC_565 | -0.004 | CTC_754 | 0.007 | DPC1_YS | -0.032 |
| DPC1_KG | 0.097 | CTC_566 | -0.008 | CTC_755 | -0.056 | DPC1_YT | -0.043 |
| DPC1_KH | -0.027 | CTC_567 | 0.012 | CTC_756 | -0.018 | DPC1_YV | -0.027 |
| DPC1_KI | 0.081 | CTC_571 | -0.017 | CTC_757 | 0.077 | DPC1_YW | 0.008 |
| DPC1_KK | 0.207 | CTC_572 | 0.020 | CTC_761 | -0.011 | DPC1_YY | 0.029 |
| DPC1_KL | 0.066 | CTC_573 | -0.011 | CTC_762 | -0.018 | ATC_C | -0.063 |

| | | | | | | | |
|---|---|---|---|---|---|---|---|
| **DPC1_KM** | **0.004** | **CTC_574** | **-0.029** | **CTC_763** | **-0.018** | **ATC_H** | **-0.135** |
| **DPC1_KN** | **-0.029** | **CTC_575** | **-0.001** | **CTC_764** | **-0.009** | **ATC_N** | **0.120** |
| **DPC1_KP** | **0.013** | **CTC_576** | **-0.004** | **CTC_765** | **-** | **ATC_O** | **0.073** |
| **DPC1_KQ** | **-0.025** | **CTC_577** | **-0.016** | **CTC_766** | **-** | **ATC_S** | **-0.076** |
| **DPC1_KR** | **0.083** | **CTC_611** | **0.012** | **CTC_767** | **-** | **BTC_T** | **-0.202** |
| **DPC1_KS** | **-0.041** | **CTC_612** | **-0.051** | **CTC_771** | **-** | **BTC_H** | **-0.193** |
| **DPC1_KT** | **0.076** | **CTC_613** | **0.030** | **CTC_772** | **-0.016** | **BTC_S** | **-0.198** |
| **DPC1_KV** | **0.050** | **CTC_614** | **0.010** | **CTC_773** | **0.085** | **BTC_D** | **-0.212** |
| **DPC1_KW** | **0.036** | **CTC_615** | **-0.007** | **CTC_774** | **-0.016** | **PCP_PC** | **0.236** |
| **DPC1_KY** | **-0.038** | **CTC_616** | **-** | **CTC_775** | **-0.016** | **PCP_NC** | **-0.049** |
| **DPC1_LA** | **0.073** | **CTC_617** | **-0.001** | **CTC_776** | **-** | **PCP_NE** | **-0.233** |
| **DPC1_LC** | **-0.026** | **CTC_621** | **-0.007** | **CTC_777** | **-** | **PCP_PO** | **-0.106** |
| **DPC1_LD** | **-0.047** | **CTC_622** | **0.013** | **CeTD_HB1** | **0.229** | **PCP_NP** | **-0.148** |
| **DPC1_LE** | **0.012** | **CTC_623** | **-0.011** | **CeTD_HB2** | **-0.114** | **PCP_AL** | **-0.082** |
| **DPC1_LF** | **-0.102** | **CTC_624** | **-0.019** | **CeTD_HB3** | **-0.131** | **PCP_CY** | **-0.075** |
| **DPC1_LG** | **-0.087** | **CTC_625** | **0.015** | **CeTD_VW1** | **-0.119** | **PCP_AR** | **-0.043** |
| **DPC1_LH** | **0.009** | **CTC_626** | **-0.002** | **CeTD_VW2** | **-0.063** | **PCP_AC** | **-0.049** |
| **DPC1_LI** | **-0.078** | **CTC_627** | **-** | **CeTD_VW3** | **0.134** | **PCP_BS** | **0.236** |
| **DPC1_LK** | **0.027** | **CTC_631** | **-0.028** | **CeTD_PO1** | **-0.130** | **PCP_NE_pH** | **-0.233** |
| **DPC1_LL** | **0.019** | **CTC_632** | **0.012** | **CeTD_PO2** | **-0.110** | **PCP_HB** | **-0.154** |
| **DPC1_LM** | **0.056** | **CTC_633** | **-0.012** | **CeTD_PO3** | **0.220** | **PCP_HL** | **0.220** |
| **DPC1_LN** | **-0.011** | **CTC_634** | **-0.016** | **CeTD_PZ1** | **-0.091** | **PCP_NT** | **-0.130** |
| **DPC1_LP** | **-0.087** | **CTC_635** | **-0.003** | **CeTD_PZ2** | **-0.107** | **PCP_HX** | **-0.098** |
| **DPC1_LQ** | **-0.038** | **CTC_636** | **-** | **CeTD_PZ3** | **0.143** | **PCP_SC** | **-0.074** |
| **DPC1_LR** | **-0.006** | **CTC_637** | **-0.070** | **CeTD_CH1** | **0.247** | **PCP_SS_HE** | **0.157** |
| **DPC1_LS** | **-0.076** | **CTC_641** | **-0.021** | **CeTD_CH2** | **-0.242** | **PCP_SS_ST** | **-0.084** |
| **DPC1_LT** | **0.000** | **CTC_642** | **0.001** | **CeTD_CH3** | **-0.050** | **PCP_SS_CO** | **-0.153** |
| **DPC1_LV** | **-0.020** | **CTC_643** | **-0.025** | **CeTD_SS1** | **0.157** | **CeTD_31_VW** | **-0.031** |
| **DPC1_LW** | **-0.059** | **CTC_644** | **-0.020** | **CeTD_SS2** | **-0.084** | **CeTD_31_PO** | **-0.016** |
| **DPC1_LY** | **0.143** | **CTC_645** | **-0.033** | **CeTD_SS3** | **-0.153** | **CeTD_31_PZ** | **0.027** |
| **DPC1_MA** | **-0.033** | **CTC_646** | **0.007** | **CeTD_SA1** | **-0.167** | **CeTD_31_CH** | **-0.134** |
| **DPC1_MC** | **-0.015** | **CTC_647** | **-** | **CeTD_SA2** | **0.229** | **CeTD_31_SS** | **-0.139** |
| **DPC1_MD** | **-0.029** | **CTC_651** | **-0.024** | **CeTD_SA3** | **-0.110** | **CeTD_31_SA** | **-0.091** |
| **DPC1_ME** | **-0.024** | **CTC_652** | **-0.030** | **CeTD_11_HB** | **0.048** | **CeTD_32_HB** | **-0.145** |
| **DPC1_MF** | **0.015** | **CTC_653** | **0.025** | **CeTD_11_VW** | **-0.011** | **CeTD_32_VW** | **-0.158** |
| **DPC1_MG** | **-0.022** | **CTC_654** | **-0.017** | **CeTD_11_PO** | **-0.024** | **CeTD_32_PO** | **-0.154** |
| **DPC1_MH** | **-0.004** | **CTC_655** | **-0.014** | **CeTD_11_PZ** | **-0.010** | **CeTD_32_PZ** | **-0.116** |
| **DPC1_MI** | **-0.018** | **CTC_656** | **0.007** | **CeTD_11_CH** | **-0.142** | **CeTD_32_CH** | **-0.064** |
| **DPC1_MK** | **0.021** | **CTC_657** | **-** | **CeTD_11_SS** | **-0.183** | **CeTD_32_SS** | **-0.261** |
| **DPC1_ML** | **0.003** | **CTC_661** | **-** | **CeTD_11_SA** | **-0.012** | **CeTD_32_SA** | **-0.007** |
| **DPC1_MM** | **0.010** | **CTC_662** | **-0.009** | **CeTD_12_HB** | **-0.187** | **CeTD_33_HB** | **-0.156** |
| **DPC1_MN** | **0.010** | **CTC_663** | **-0.014** | **CeTD_12_VW** | **-0.282** | **CeTD_33_VW** | **-0.009** |

| | | | | | | | |
|---|---|---|---|---|---|---|---|
| DPC1_MP | 0.023 | CTC_664 | - | CeTD_12_PO | -0.135 | CeTD_33_PO | 0.048 |
| DPC1_MQ | -0.017 | CTC_665 | -0.006 | CeTD_12_PZ | -0.143 | CeTD_33_PZ | -0.048 |
| DPC1_MR | -0.030 | CTC_666 | -0.006 | CeTD_12_CH | -0.061 | CeTD_33_CH | -0.170 |
| DPC1_MS | -0.003 | CTC_667 | - | CeTD_12_SS | -0.167 | CeTD_33_SS | -0.026 |
| DPC1_MT | -0.045 | CTC_671 | - | CeTD_12_SA | -0.111 | CeTD_33_SA | -0.136 |
| DPC1_MV | -0.023 | CTC_672 | -0.017 | CeTD_13_HB | -0.044 | CeTD_0_p_HB1 | - |
| DPC1_MW | -0.007 | CTC_673 | -0.014 | CeTD_13_VW | -0.031 | CeTD_25_p_HB1 | 0.028 |
| DPC1_MY | 0.085 | CTC_674 | - | CeTD_13_PO | -0.069 | CeTD_50_p_HB1 | 0.025 |
| DPC1_NA | -0.026 | CTC_675 | - | CeTD_13_PZ | 0.001 | PCP_Z1 | 0.118 |
| DPC1_NC | -0.013 | CTC_676 | - | CeTD_13_CH | -0.259 | PCP_Z2 | 0.082 |
| DPC1_ND | 0.006 | CTC_677 | 0.085 | CeTD_13_SS | -0.188 | PCP_Z3 | -0.204 |
| DPC1_NE | -0.016 | CTC_711 | -0.049 | CeTD_13_SA | -0.019 | PCP_Z4 | 0.116 |
| DPC1_NF | -0.036 | CTC_712 | -0.052 | CeTD_21_HB | -0.171 | PCP_Z5 | 0.043 |
| DPC1_NG | -0.009 | CTC_713 | -0.036 | CeTD_21_VW | -0.131 | RRI_A | -0.052 |
| DPC1_NH | 0.010 | CTC_714 | -0.028 | CeTD_21_PO | -0.030 | RRI_C | -0.090 |
| DPC1_NI | 0.041 | CTC_715 | 0.106 | CeTD_21_PZ | -0.049 | RRI_D | -0.033 |
| DPC1_NK | -0.002 | CTC_716 | -0.069 | CeTD_21_CH | -0.005 | RRI_E | -0.050 |
| DPC1_NL | -0.005 | CTC_717 | -0.027 | CeTD_21_SS | 0.037 | RRI_F | -0.127 |
| DPC1_NM | -0.001 | CTC_721 | -0.050 | CeTD_21_SA | -0.121 | RRI_G | -0.165 |
| DPC1_NN | 0.014 | CTC_722 | -0.010 | CeTD_22_HB | -0.139 | RRI_H | -0.059 |
| DPC1_NP | -0.036 | CTC_723 | -0.034 | CeTD_22_VW | -0.027 | RRI_I | -0.078 |
| DPC1_NQ | -0.022 | CTC_724 | -0.027 | CeTD_22_PO | -0.152 | RRI_K | 0.112 |
| DPC1_NR | -0.022 | CTC_725 | -0.058 | CeTD_22_PZ | -0.160 | PCP_SA_BU | -0.167 |
| DPC1_NS | 0.083 | CTC_726 | - | CeTD_22_CH | -0.089 | PCP_SA_EX | 0.166 |
| DPC1_NT | -0.025 | CTC_727 | 0.049 | CeTD_22_SS | -0.016 | DPC1_RD | 0.000 |
| DPC1_NV | -0.025 | CTC_731 | -0.007 | CeTD_22_SA | -0.106 | DPC1_RE | -0.014 |
| DPC1_NW | -0.045 | CTC_732 | -0.049 | CeTD_23_HB | 0.009 | DPC1_RF | -0.012 |
| DPC1_NY | -0.025 | CTC_733 | -0.054 | CeTD_23_VW | 0.087 | DPC1_RG | -0.056 |
| DPC1_PA | -0.001 | CTC_734 | -0.045 | CeTD_23_PO | -0.010 | DPC1_RH | 0.046 |
| DPC1_PC | -0.021 | CTC_735 | -0.039 | CeTD_23_PZ | -0.004 | DPC1_RI | 0.057 |
| DPC1_PD | -0.018 | CTC_736 | -0.024 | CeTD_23_CH | 0.018 | DPC1_RK | 0.095 |
| DPC1_PE | -0.003 | CTC_737 | -0.074 | CeTD_23_SS | -0.265 | DPC1_RL | 0.064 |
| DPC1_PF | -0.025 | CTC_741 | - | CeTD_23_SA | -0.062 | DPC1_RM | -0.011 |
| DPC1_PG | -0.030 | CTC_742 | -0.025 | CeTD_31_HB | -0.045 | DPC1_RN | -0.012 |
| DPC1_PH | -0.045 | DPC1_TA | -0.077 | DPC1_VT | 0.010 | DPC1_RP | -0.031 |
| DPC1_PI | -0.058 | DPC1_TC | -0.073 | DPC1_VV | -0.036 | DPC1_RQ | -0.007 |
| DPC1_PK | -0.056 | DPC1_TD | -0.011 | DPC1_VW | -0.003 | DPC1_RR | 0.028 |
| DPC1_PL | 0.013 | DPC1_TE | -0.020 | DPC1_VY | 0.093 | DPC1_RS | -0.022 |
| DPC1_PM | 0.002 | DPC1_TF | -0.026 | DPC1_WA | 0.003 | DPC1_RT | 0.000 |
| DPC1_PN | 0.002 | DPC1_TG | -0.053 | DPC1_WC | -0.052 | DPC1_RV | -0.046 |
| DPC1_PP | 0.060 | DPC1_TH | -0.028 | DPC1_WD | -0.047 | DPC1_RW | 0.014 |
| DPC1_PQ | -0.028 | DPC1_TI | -0.058 | DPC1_WE | -0.014 | DPC1_RY | -0.027 |

| | | | | | | | |
|---|---|---|---|---|---|---|---|
| **DPC1_PR** | **-0.031** | **DPC1_TK** | **0.009** | **DPC1_WF** | **-0.047** | **DPC1_SA** | **-0.044** |
| **DPC1_PS** | **-0.043** | **DPC1_TL** | **0.084** | **DPC1_WG** | **0.020** | **DPC1_SC** | **-0.036** |
| **DPC1_PT** | **0.004** | **DPC1_TM** | **-0.019** | **DPC1_WH** | **0.032** | **DPC1_SD** | **-0.019** |
| **DPC1_PV** | **-0.016** | **DPC1_TN** | **-0.012** | **DPC1_WI** | **-0.015** | **DPC1_SE** | **-0.004** |
| **DPC1_PW** | **-0.022** | **DPC1_TP** | **-0.067** | **DPC1_WK** | **0.024** | **DPC1_SF** | **-0.054** |
| **DPC1_PY** | **-0.008** | **DPC1_TQ** | **-0.018** | **DPC1_WL** | **-0.022** | **DPC1_SG** | **-0.008** |
| **DPC1_QA** | **-0.012** | **DPC1_TR** | **-0.043** | **DPC1_WM** | **-0.030** | **DPC1_SH** | **-0.032** |
| **DPC1_QC** | **-0.011** | **DPC1_TS** | **-0.040** | **DPC1_WN** | **-0.030** | **DPC1_SI** | **-0.020** |
| **DPC1_QD** | **-0.011** | **DPC1_TT** | **-0.056** | **PCP_SA_IN** | **-0.110** | **DPC1_SK** | **0.046** |
| **DPC1_QE** | **-0.004** | **DPC1_TV** | **0.050** | **PCP_TN** | **-0.108** | **DPC1_SL** | **-0.074** |
| **DPC1_QF** | **-0.007** | **DPC1_TW** | **0.035** | **PCP_SM** | **-0.127** | **DPC1_SM** | **-0.031** |
| **DPC1_QG** | **-0.012** | **DPC1_TY** | **-0.014** | **PCP_LR** | **0.127** | **DPC1_SN** | **-0.020** |
| **DPC1_QH** | **-0.035** | **DPC1_VA** | **0.011** | **DPC1_VP** | **-0.030** | **DPC1_SP** | **-0.017** |
| **DPC1_QI** | **-0.043** | **DPC1_VC** | **-0.045** | **DPC1_VQ** | **0.083** | **DPC1_SQ** | **-0.023** |
| **DPC1_QK** | **0.016** | **DPC1_VD** | **0.035** | **DPC1_VR** | **-0.043** | **DPC1_SR** | **-0.003** |
| **DPC1_QL** | **-0.019** | **DPC1_VE** | **0.026** | **DPC1_VS** | **-0.003** | **DPC1_SS** | **0.002** |
| **DPC1_QM** | **0.008** | **DPC1_VF** | **-0.028** | **DPC1_QT** | **-0.001** | **DPC1_ST** | **-0.073** |
| **DPC1_QN** | **-0.033** | **DPC1_VG** | **-0.034** | **DPC1_QV** | **0.050** | **DPC1_SV** | **-0.055** |
| **DPC1_QP** | **-0.005** | **DPC1_VH** | **-0.034** | **DPC1_QW** | **-0.007** | **DPC1_SW** | **-0.002** |
| **DPC1_QQ** | **-0.016** | **DPC1_VI** | **0.003** | **DPC1_QY** | **-0.028** | **DPC1_SY** | **-0.044** |
| **DPC1_QR** | **0.040** | **DPC1_VK** | **0.095** | **DPC1_RA** | **0.003** | **DPC1_VM** | **-0.031** |
| **DPC1_QS** | **-0.002** | **DPC1_VL** | **-0.091** | **DPC1_RC** | **0.000** | **DPC1_VN** | **0.031** |

### Supplementary Table S4: The detailed evaluation metrics of classification models used to predict hemolytic peptides.

#### Supplementary Table S4.1: Evaluation metrics of ML models using compositional features.

**Feature: Amino Acid Composition ( 20 descriptors)**

| Model | Dataset | Sn | Sp | Acc | MCC | AUC |
|---|---|---|---|---|---|---|
| ET | **Cross-validation** | 0.787 | 0.713 | 0.754 | 0.501 | 0.826 |
| | **Independent** | 0.834 | 0.686 | 0.767 | 0.528 | 0.807 |
| SVM | **Cross-validation** | 0.772 | 0.633 | 0.710 | 0.410 | 0.769 |
| | **Independent** | 0.777 | 0.611 | 0.702 | 0.395 | 0.759 |

| | | | | | | |
|---|---|---|---|---|---|---|
| XGB | **Cross-validation** | 0.794 | 0.703 | 0.753 | 0.501 | 0.820 |
| | **Independent** | 0.829 | 0.680 | 0.762 | 0.517 | 0.831 |
| RF | **Cross-validation** | 0.799 | 0.701 | 0.755 | 0.503 | 0.832 |
| | **Independent** | 0.839 | 0.663 | 0.759 | 0.512 | 0.831 |
| ET_tuned | **Cross-validation** | 0.794 | 0.713 | 0.758 | 0.509 | 0.825 |
| | **Independent** | 0.825 | 0.691 | 0.764 | 0.522 | 0.806 |
| MLPC | **Cross-validation** | 0.749 | 0.717 | 0.735 | 0.467 | 0.807 |
| | **Independent** | 0.829 | 0.737 | 0.788 | 0.570 | 0.834 |
| GB | **Cross-validation** | 0.797 | 0.674 | 0.742 | 0.476 | 0.813 |
| | **Independent** | 0.825 | 0.686 | 0.762 | 0.517 | 0.827 |
| DT | **Cross-validation** | 0.724 | 0.647 | 0.690 | 0.373 | 0.687 |
| | **Independent** | 0.678 | 0.657 | 0.668 | 0.334 | 0.663 |
| LR | **Cross-validation** | 0.785 | 0.623 | 0.712 | 0.414 | 0.772 |
| | **Independent** | 0.796 | 0.600 | 0.707 | 0.406 | 0.755 |

#### Feature: Di-peptide Composition ( 400 descriptors)

| Model | Dataset | Sn | Sp | Acc | MCC | AUC |
|---|---|---|---|---|---|---|
| ET | **Cross-validation** | 0.773 | 0.697 | 0.739 | 0.471 | 0.814 |
| | **Independent** | 0.829 | 0.646 | 0.746 | 0.486 | 0.823 |
| SVM | **Cross-validation** | 0.722 | 0.652 | 0.691 | 0.375 | 0.732 |
| | **Independent** | 0.773 | 0.669 | 0.725 | 0.444 | 0.770 |
| XGB | **Cross-validation** | 0.785 | 0.684 | 0.740 | 0.471 | 0.814 |
| | **Independent** | 0.858 | 0.697 | 0.785 | 0.565 | 0.845 |
| RF | **Cross-validation** | 0.807 | 0.703 | 0.760 | 0.513 | 0.825 |
| | **Independent** | 0.877 | 0.651 | 0.775 | 0.547 | 0.837 |
| ET_tuned | **Cross-validation** | 0.767 | 0.685 | 0.731 | 0.454 | 0.813 |
| | **Independent** | 0.839 | 0.669 | 0.762 | 0.518 | 0.823 |
| MLPC | **Cross-validation** | 0.745 | 0.691 | 0.721 | 0.435 | 0.788 |
| | **Independent** | 0.796 | 0.691 | 0.749 | 0.491 | 0.806 |
| GB | **Cross-validation** | 0.793 | 0.644 | 0.727 | 0.444 | 0.796 |
| | **Independent** | 0.844 | 0.646 | 0.754 | 0.502 | 0.816 |
| DT | **Cross-validation** | 0.733 | 0.637 | 0.690 | 0.372 | 0.685 |
| | **Independent** | 0.782 | 0.691 | 0.741 | 0.476 | 0.736 |
| LR | **Cross-validation** | 0.728 | 0.668 | 0.701 | 0.396 | 0.747 |
| | **Independent** | 0.796 | 0.669 | 0.738 | 0.470 | 0.795 |

**Feature: AAC and DPC ( 420 descriptors)**

| Model | Dataset | Sn | Sp | Acc | MCC | AUC |
|---|---|---|---|---|---|---|
| ET | Cross-validation | 0.702 | 0.775 | 0.742 | 0.479 | 0.812 |
| | Independent | 0.749 | 0.792 | 0.772 | 0.541 | 0.852 |
| SVM | Cross-validation | 0.642 | 0.696 | 0.671 | 0.338 | 0.705 |
| | Independent | 0.704 | 0.758 | 0.733 | 0.463 | 0.784 |
| XGB | Cross-validation | 0.697 | 0.780 | 0.742 | 0.481 | 0.815 |
| | Independent | 0.793 | 0.792 | 0.793 | 0.585 | 0.865 |
| RF | Cross-validation | 0.698 | 0.791 | 0.748 | 0.493 | 0.818 |
| | Independent | 0.749 | 0.855 | 0.806 | 0.609 | 0.875 |
| ET_tuned | Cross-validation | 0.708 | 0.764 | 0.738 | 0.473 | 0.808 |
| | Independent | 0.704 | 0.831 | 0.772 | 0.541 | 0.843 |
| MLPC | Cross-validation | 0.708 | 0.710 | 0.709 | 0.418 | 0.784 |
| | Independent | 0.676 | 0.754 | 0.718 | 0.431 | 0.808 |
| GB | Cross-validation | 0.663 | 0.783 | 0.727 | 0.450 | 0.789 |
| | Independent | 0.732 | 0.807 | 0.772 | 0.541 | 0.856 |
| DT | Cross-validation | 0.650 | 0.704 | 0.679 | 0.355 | 0.677 |
| | Independent | 0.659 | 0.725 | 0.694 | 0.385 | 0.692 |
| LR | Cross-validation | 0.669 | 0.713 | 0.692 | 0.382 | 0.742 |
| | Independent | 0.659 | 0.787 | 0.728 | 0.451 | 0.806 |

**Feature: Physico-chemical properties (30 descriptors)**

| Model | Dataset | Sn | Sp | Acc | MCC | AUC |
|---|---|---|---|---|---|---|
| ET | Cross-validation | 0.794 | 0.706 | 0.755 | 0.503 | 0.820 |
| | Independent | 0.853 | 0.674 | 0.772 | 0.539 | 0.820 |
| SVM | Cross-validation | 0.769 | 0.637 | 0.710 | 0.411 | 0.774 |
| | Independent | 0.782 | 0.623 | 0.710 | 0.411 | 0.772 |
| XGB | Cross-validation | 0.790 | 0.706 | 0.753 | 0.498 | 0.825 |
| | Independent | 0.853 | 0.680 | 0.775 | 0.544 | 0.833 |
| RF | Cross-validation | 0.803 | 0.693 | 0.754 | 0.500 | 0.832 |
| | Independent | 0.848 | 0.674 | 0.769 | 0.534 | 0.854 |
| ET_tuned | Cross-validation | 0.792 | 0.714 | 0.757 | 0.509 | 0.816 |
| | Independent | 0.848 | 0.634 | 0.751 | 0.498 | 0.815 |
| MLPC | Cross-validation | 0.778 | 0.649 | 0.720 | 0.432 | 0.791 |
| | Independent | 0.820 | 0.634 | 0.736 | 0.464 | 0.799 |
| GB | Cross-validation | 0.801 | 0.690 | 0.751 | 0.495 | 0.823 |
| | Independent | 0.848 | 0.674 | 0.769 | 0.534 | 0.854 |
| DT | Cross-validation | 0.689 | 0.681 | 0.685 | 0.368 | 0.689 |
| | Independent | 0.725 | 0.600 | 0.668 | 0.328 | 0.661 |
| LR | Cross-validation | 0.787 | 0.615 | 0.710 | 0.410 | 0.770 |
| | Independent | 0.787 | 0.617 | 0.710 | 0.411 | 0.774 |

**Feature: Composition-enhanced Transition and Distribution (189 descriptors)**

| Model | Dataset | Sn | Sp | Acc | MCC | AUC |
|---|---|---|---|---|---|---|
| ET | Cross-validation | 0.786 | 0.732 | 0.762 | 0.519 | 0.815 |
| | Independent | 0.839 | 0.720 | 0.785 | 0.565 | 0.840 |
| SVM | Cross-validation | 0.779 | 0.655 | 0.723 | 0.438 | 0.785 |
| | Independent | 0.815 | 0.623 | 0.728 | 0.448 | 0.815 |
| XGB | Cross-validation | 0.789 | 0.710 | 0.754 | 0.502 | 0.841 |
| | Independent | 0.825 | 0.709 | 0.772 | 0.538 | 0.841 |
| | Cross-validation | 0.838 | 0.733 | 0.784 | 0.532 | 0.835 |

| | | | | | | |
|---|---|---|---|---|---|---|
| RF | **Independent** | 0.867 | 0.720 | 0.801 | 0.597 | 0.856 |
| ET_tuned | **Cross-validation** | 0.787 | 0.728 | 0.760 | 0.516 | 0.813 |
| | **Independent** | 0.839 | 0.714 | 0.782 | 0.559 | 0.845 |
| MLPC | **Cross-validation** | 0.746 | 0.659 | 0.707 | 0.431 | 0.807 |
| | **Independent** | 0.829 | 0.691 | 0.767 | 0.528 | 0.827 |
| GB | **Cross-validation** | 0.797 | 0.704 | 0.756 | 0.505 | 0.838 |
| | **Independent** | 0.825 | 0.680 | 0.759 | 0.512 | 0.844 |
| DT | **Cross-validation** | 0.720 | 0.650 | 0.689 | 0.370 | 0.684 |
| | **Independent** | 0.777 | 0.663 | 0.725 | 0.444 | 0.719 |
| LR | **Cross-validation** | 0.790 | 0.633 | 0.720 | 0.430 | 0.793 |
| | **Independent** | 0.806 | 0.629 | 0.725 | 0.443 | 0.793 |

**Feature: AAC + DPC + CeTD (450 descriptors)**

| Model | Dataset | Sn | Sp | Acc | MCC | AUC |
|---|---|---|---|---|---|---|
| SVM | **Cross-validation** | 0.680 | 0.780 | 0.734 | 0.464 | 0.800 |
| | **Independent** | 0.726 | 0.831 | 0.782 | 0.562 | 0.858 |
| XGB | **Cross-validation** | 0.706 | 0.780 | 0.746 | 0.489 | 0.820 |
| | **Independent** | 0.788 | 0.831 | 0.811 | 0.619 | 0.875 |
| RF | **Cross-validation** | 0.749 | 0.822 | 0.790 | 0.623 | 0.828 |
| | **Independent** | 0.737 | 0.850 | 0.775 | 0.598 | 0.841 |
| ET | **Cross-validation** | 0.762 | 0.788 | 0.747 | 0.492 | 0.817 |
| | **Independent** | 0.726 | 0.831 | 0.782 | 0.562 | 0.859 |
| MLPC | **Cross-validation** | 0.701 | 0.732 | 0.718 | 0.433 | 0.787 |
| | **Independent** | 0.698 | 0.758 | 0.731 | 0.458 | 0.814 |
| GB | **Cross-validation** | 0.685 | 0.799 | 0.747 | 0.490 | 0.805 |
| | **Independent** | 0.749 | 0.826 | 0.790 | 0.577 | 0.867 |
| DT | **Cross-validation** | 0.657 | 0.725 | 0.694 | 0.383 | 0.691 |
| | **Independent** | 0.665 | 0.763 | 0.718 | 0.431 | 0.714 |
| LR | **Cross-validation** | 0.664 | 0.716 | 0.692 | 0.382 | 0.742 |
| | **Independent** | 0.642 | 0.792 | 0.723 | 0.441 | 0.811 |

**Feature: AAC + DPC + PCP (450 descriptors)**

| Model | Dataset | Sn | Sp | Acc | MCC | AUC |
|---|---|---|---|---|---|---|
| SVM | **Cross-validation** | 0.680 | 0.780 | 0.734 | 0.464 | 0.800 |
| | **Independent** | 0.726 | 0.831 | 0.782 | 0.562 | 0.858 |
| XGB | **Cross-validation** | 0.706 | 0.780 | 0.746 | 0.489 | 0.820 |
| | **Independent** | 0.788 | 0.831 | 0.811 | 0.619 | 0.875 |
| RF | **Cross-validation** | 0.749 | 0.822 | 0.790 | 0.623 | 0.828 |
| | **Independent** | 0.737 | 0.850 | 0.801 | 0.598 | 0.881 |
| ET | **Cross-validation** | 0.716 | 0.774 | 0.747 | 0.492 | 0.817 |
| | **Independent** | 0.726 | 0.831 | 0.782 | 0.562 | 0.859 |
| MLPC | **Cross-validation** | 0.701 | 0.732 | 0.718 | 0.433 | 0.787 |
| | **Independent** | 0.698 | 0.758 | 0.731 | 0.458 | 0.814 |
| GB | **Cross-validation** | 0.685 | 0.799 | 0.747 | 0.490 | 0.805 |
| | **Independent** | 0.749 | 0.826 | 0.790 | 0.577 | 0.867 |
| DT | **Cross-validation** | 0.657 | 0.725 | 0.694 | 0.383 | 0.691 |
| | **Independent** | 0.665 | 0.763 | 0.718 | 0.431 | 0.714 |
| LR | **Cross-validation** | 0.664 | 0.716 | 0.692 | 0.382 | 0.742 |
| | **Independent** | 0.642 | 0.792 | 0.723 | 0.441 | 0.811 |

**Feature: ALLCOMP (1192 descriptors)**

| Model | Dataset | Sn | Sp | Acc | MCC | AUC |
|---|---|---|---|---|---|---|
| RF | Cross-validation | 0.229 | 0.828 | 0.805 | 0.628 | 0.842 |
| | Independent | 0.749 | 0.839 | 0.801 | 0.610 | 0.878 |
| SVM | Cross-validation | 0.663 | 0.705 | 0.686 | 0.368 | 0.749 |
| | Independent | 0.676 | 0.787 | 0.736 | 0.467 | 0.819 |
| XGB | Cross-validation | 0.760 | 0.796 | 0.779 | 0.556 | 0.842 |
| | Independent | 0.771 | 0.821 | 0.798 | 0.593 | 0.863 |
| ET | Cross-validation | 0.743 | 0.809 | 0.779 | 0.554 | 0.845 |
| | Independent | 0.765 | 0.870 | 0.821 | 0.641 | 0.868 |
| ET_tuned | Cross-validation | 0.737 | 0.791 | 0.766 | 0.529 | 0.839 |
| | Independent | 0.754 | 0.836 | 0.798 | 0.593 | 0.860 |
| MLPC | Cross-validation | 0.634 | 0.686 | 0.662 | 0.368 | 0.781 |
| | Independent | 0.743 | 0.778 | 0.762 | 0.521 | 0.830 |
| GB | Cross-validation | 0.695 | 0.797 | 0.750 | 0.496 | 0.825 |
| | Independent | 0.760 | 0.836 | 0.801 | 0.598 | 0.870 |
| DT | Cross-validation | 0.691 | 0.731 | 0.712 | 0.422 | 0.711 |
| | Independent | 0.709 | 0.783 | 0.749 | 0.494 | 0.746 |
| LR | Cross-validation | 0.656 | 0.767 | 0.716 | 0.428 | 0.779 |
| | Independent | 0.648 | 0.812 | 0.736 | 0.468 | 0.806 |

**Feature: ALLCOMP-excluding SOC (1192 descriptors)**

| Model | Dataset | Sn | Sp | Acc | MCC | AUC |
|---|---|---|---|---|---|---|
| RF | Cross-validation | 0.743 | 0.809 | 0.779 | 0.554 | 0.845 |
| | Independent | 0.765 | 0.870 | 0.821 | 0.641 | 0.888 |
| SVM | Cross-validation | 0.663 | 0.705 | 0.686 | 0.368 | 0.749 |
| | Independent | 0.676 | 0.787 | 0.736 | 0.467 | 0.819 |
| XGB | Cross-validation | 0.760 | 0.796 | 0.779 | 0.556 | 0.842 |
| | Independent | 0.771 | 0.821 | 0.798 | 0.593 | 0.863 |
| ET | Cross-validation | 0.743 | 0.809 | 0.779 | 0.554 | 0.845 |
| | Independent | 0.765 | 0.870 | 0.821 | 0.641 | 0.868 |
| ET_tuned | Cross-validation | 0.737 | 0.791 | 0.766 | 0.529 | 0.839 |
| | Independent | 0.754 | 0.836 | 0.798 | 0.593 | 0.860 |
| MLPC | Cross-validation | 0.634 | 0.686 | 0.662 | 0.368 | 0.781 |
| | Independent | 0.743 | 0.778 | 0.762 | 0.521 | 0.830 |
| GB | Cross-validation | 0.695 | 0.797 | 0.750 | 0.496 | 0.825 |
| | Independent | 0.760 | 0.836 | 0.801 | 0.598 | 0.870 |
| DT | Cross-validation | 0.691 | 0.731 | 0.712 | 0.422 | 0.711 |
| | Independent | 0.709 | 0.783 | 0.749 | 0.494 | 0.746 |
| LR | Cross-validation | 0.656 | 0.767 | 0.716 | 0.428 | 0.779 |
| | Independent | 0.648 | 0.812 | 0.736 | 0.468 | 0.806 |

#### Supplementary Table S4.2: Evaluation metrics of ML models using Protein language models embeddings.

ProtBERT embeddings

| Model | Dataset | Sn | Sp | Acc | MCC | AUC |
|---|---|---|---|---|---|---|

| | | | | | | |
|---|---|---|---|---|---|---|
| ET | **Cross-validation** | 0.695 | 0.751 | 0.726 | 0.446 | 0.789 |
| | **Independent** | 0.651 | 0.820 | 0.744 | 0.480 | 0.815 |
| SVM | **Cross-validation** | 0.780 | 0.778 | 0.779 | 0.556 | 0.847 |
| | **Independent** | 0.846 | 0.834 | 0.839 | 0.678 | 0.874 |
| XGB | **Cross-validation** | 0.723 | 0.751 | 0.738 | 0.474 | 0.808 |
| | **Independent** | 0.703 | 0.825 | 0.769 | 0.533 | 0.843 |
| RF | **Cross-validation** | 0.736 | 0.755 | 0.747 | 0.490 | 0.814 |
| | **Independent** | 0.737 | 0.825 | 0.785 | 0.565 | 0.842 |
| RT_tuned | **Cross-validation** | 0.688 | 0.755 | 0.725 | 0.444 | 0.786 |
| | **Independent** | 0.663 | 0.825 | 0.751 | 0.496 | 0.810 |
| MLPC | **Cross-validation** | 0.767 | 0.787 | 0.778 | 0.553 | 0.850 |
| | **Independent** | 0.840 | 0.839 | 0.839 | 0.677 | 0.882 |
| GB | **Cross-validation** | 0.771 | 0.759 | 0.764 | 0.528 | 0.831 |
| | **Independent** | 0.789 | 0.829 | 0.811 | 0.618 | 0.860 |
| DT | **Cross-validation** | 0.672 | 0.710 | 0.693 | 0.383 | 0.691 |
| | **Independent** | 0.669 | 0.773 | 0.725 | 0.444 | 0.721 |
| LR | **Cross-validation** | 0.774 | 0.785 | 0.780 | 0.557 | 0.849 |
| | **Independent** | 0.834 | 0.844 | 0.839 | 0.677 | 0.880 |

ESM2-t33 embeddings

| **Model** | **Dataset** | **Sn** | **Sp** | **Acc** | **MCC** | **AUC** |
|---|---|---|---|---|---|---|
| ET | **Cross-validation** | 0.715 | 0.831 | 0.777 | 0.552 | 0.854 |
| | **Independent** | 0.732 | 0.855 | 0.798 | 0.594 | 0.873 |
| SVM | **Cross-validation** | 0.736 | 0.787 | 0.764 | 0.526 | 0.831 |
| | **Independent** | 0.749 | 0.855 | 0.806 | 0.609 | 0.864 |
| XGB | **Cross-validation** | 0.729 | 0.805 | 0.770 | 0.538 | 0.854 |
| | **Independent** | 0.737 | 0.831 | 0.788 | 0.572 | 0.861 |
| RF | **Cross-validation** | 0.691 | 0.824 | 0.762 | 0.522 | 0.847 |
| | **Independent** | 0.693 | 0.841 | 0.772 | 0.541 | 0.855 |
| RT_tuned | **Cross-validation** | 0.705 | 0.826 | 0.770 | 0.537 | 0.847 |
| | **Independent** | 0.704 | 0.870 | 0.793 | 0.585 | 0.862 |
| MLPC | **Cross-validation** | 0.775 | 0.780 | 0.778 | 0.555 | 0.835 |
| | **Independent** | 0.832 | 0.821 | 0.826 | 0.652 | 0.871 |
| GB | **Cross-validation** | 0.711 | 0.804 | 0.761 | 0.519 | 0.837 |
| | **Independent** | 0.715 | 0.865 | 0.795 | 0.589 | 0.857 |
| DT | **Cross-validation** | 0.638 | 0.715 | 0.679 | 0.354 | 0.676 |
| | **Independent** | 0.609 | 0.676 | 0.645 | 0.286 | 0.643 |
| LR | **Cross-validation** | 0.718 | 0.766 | 0.744 | 0.485 | 0.823 |
| | **Independent** | 0.715 | 0.826 | 0.775 | 0.546 | 0.851 |

BioBERT embeddings

| **Model** | **Dataset** | **Sn** | **Sp** | **Acc** | **MCC** | **AUC** |
|---|---|---|---|---|---|---|
| ET | **Cross-validation** | 0.561 | 0.809 | 0.699 | 0.385 | 0.781 |
| | **Independent** | 0.571 | 0.791 | 0.692 | 0.374 | 0.776 |
| SVM | **Cross-validation** | 0.647 | 0.723 | 0.689 | 0.371 | 0.756 |
| | **Independent** | 0.646 | 0.777 | 0.718 | 0.427 | 0.764 |
| XGB | **Cross-validation** | 0.620 | 0.783 | 0.710 | 0.410 | 0.778 |
| | **Independent** | 0.623 | 0.791 | 0.715 | 0.422 | 0.783 |
| RF | **Cross-validation** | 0.570 | 0.809 | 0.703 | 0.393 | 0.777 |
| | **Independent** | 0.537 | 0.777 | 0.668 | 0.325 | 0.752 |
| RT_tuned | **Cross-validation** | 0.569 | 0.820 | 0.708 | 0.404 | 0.775 |
| | **Independent** | 0.594 | 0.787 | 0.699 | 0.390 | 0.788 |
| MLPC | **Cross-validation** | 0.666 | 0.735 | 0.705 | 0.402 | 0.771 |
| | **Independent** | 0.686 | 0.815 | 0.756 | 0.507 | 0.792 |
| GB | **Cross-validation** | 0.612 | 0.762 | 0.695 | 0.380 | 0.756 |

| | | | | | | |
|---|---|---|---|---|---|---|
| | **Independent** | 0.623 | 0.782 | 0.710 | 0.411 | 0.768 |
| DT | **Cross-validation** | 0.570 | 0.638 | 0.608 | 0.208 | 0.604 |
| | **Independent** | 0.577 | 0.649 | 0.617 | 0.226 | 0.613 |
| LR | **Cross-validation** | 0.643 | 0.754 | 0.705 | 0.400 | 0.774 |
| | **Independent** | 0.691 | 0.806 | 0.754 | 0.501 | 0.788 |

ESM2-t36 embeddings

| **Model** | **Dataset** | **Sn** | **Sp** | **Acc** | **MCC** | **AUC** |
|---|---|---|---|---|---|---|
| ET | **Cross-validation** | 0.676 | 0.835 | 0.761 | 0.520 | 0.837 |
| | **Independent** | 0.698 | 0.845 | 0.777 | 0.552 | 0.854 |
| SVM | **Cross-validation** | 0.715 | 0.789 | 0.755 | 0.506 | 0.832 |
| | **Independent** | 0.726 | 0.874 | 0.806 | 0.610 | 0.868 |
| XGB | **Cross-validation** | 0.713 | 0.808 | 0.764 | 0.525 | 0.829 |
| | **Independent** | 0.749 | 0.860 | 0.808 | 0.614 | 0.869 |
| RF | **Cross-validation** | 0.671 | 0.822 | 0.753 | 0.502 | 0.823 |
| | **Independent** | 0.715 | 0.855 | 0.790 | 0.578 | 0.842 |
| RT_tuned | **Cross-validation** | 0.661 | 0.832 | 0.753 | 0.504 | 0.833 |
| | **Independent** | 0.698 | 0.870 | 0.790 | 0.580 | 0.856 |
| MLPC | **Cross-validation** | 0.712 | 0.801 | 0.760 | 0.517 | 0.840 |
| | **Independent** | 0.765 | 0.826 | 0.798 | 0.593 | 0.869 |
| GB | **Cross-validation** | 0.676 | 0.802 | 0.744 | 0.484 | 0.818 |
| | **Independent** | 0.721 | 0.855 | 0.793 | 0.583 | 0.863 |
| DT | **Cross-validation** | 0.636 | 0.670 | 0.655 | 0.307 | 0.653 |
| | **Independent** | 0.659 | 0.657 | 0.658 | 0.315 | 0.658 |
| LR | **Cross-validation** | 0.705 | 0.802 | 0.757 | 0.511 | 0.832 |
| | **Independent** | 0.698 | 0.860 | 0.785 | 0.568 | 0.859 |

ESM2-t6 embeddings

| **Model** | **Dataset** | **Sn** | **Sp** | **Acc** | **MCC** | **AUC** |
|---|---|---|---|---|---|---|
| ET | **Cross-validation** | 0.715 | 0.825 | 0.774 | 0.545 | 0.848 |
| | **Independent** | 0.732 | 0.845 | 0.793 | 0.583 | 0.857 |
| SVM | **Cross-validation** | 0.727 | 0.778 | 0.755 | 0.508 | 0.812 |
| | **Independent** | 0.698 | 0.836 | 0.772 | 0.541 | 0.832 |
| XGB | **Cross-validation** | 0.736 | 0.814 | 0.778 | 0.553 | 0.846 |
| | **Independent** | 0.743 | 0.797 | 0.772 | 0.541 | 0.855 |
| RF | **Cross-validation** | 0.691 | 0.820 | 0.760 | 0.517 | 0.835 |
| | **Independent** | 0.687 | 0.855 | 0.777 | 0.553 | 0.852 |
| RT_tuned | **Cross-validation** | 0.713 | 0.824 | 0.773 | 0.542 | 0.843 |
| | **Independent** | 0.676 | 0.836 | 0.762 | 0.521 | 0.842 |
| MLPC | **Cross-validation** | 0.768 | 0.767 | 0.768 | 0.535 | 0.839 |
| | **Independent** | 0.709 | 0.841 | 0.780 | 0.557 | 0.853 |
| GB | **Cross-validation** | 0.704 | 0.804 | 0.758 | 0.513 | 0.834 |
| | **Independent** | 0.698 | 0.831 | 0.769 | 0.536 | 0.824 |
| DT | **Cross-validation** | 0.681 | 0.720 | 0.702 | 0.402 | 0.700 |
| | **Independent** | 0.642 | 0.720 | 0.684 | 0.363 | 0.681 |
| LR | **Cross-validation** | 0.704 | 0.777 | 0.743 | 0.483 | 0.802 |
| | **Independent** | 0.665 | 0.831 | 0.754 | 0.505 | 0.820 |

#### Supplementary Table S4.3: Evaluation metrics of protein language models

| **Model** | **Dataset** | **Sn** | **Sp** | **Acc** | **MCC** | **AUC** | **model size** |
|---|---|---|---|---|---|---|---|

| | | | | | | | |
|---|---|---|---|---|---|---|---|
| **ESM2-t33** | **Independent** | 0.861 | 0.700 | 0.791 | 0.573 | 0.870 | 2.488 gb |
| **ESM2-t30** | **Independent** | 0.842 | 0.780 | 0.813 | 0.599 | 0.851 | 567.79 mb |
| **ESM2-t12** | **Independent** | 0.700 | 0.790 | 0.742 | 0.490 | 0.831 | 129.75 mb |
| **ESM2-t6** | **Independent** | 0.761 | 0.840 | 0.795 | 0.591 | 0.870 | 29.95 mb |
| **BioBERT** | **Independent** | 0.637 | 0.805 | 0.715 | 0.474 | 0.800 | 413 mb |
| **ProtBERT** | **Independent** | 0.849 | 0.736 | 0.800 | 0.602 | 0.875 | 1.8 gb |

#### Supplementary Table S5: The detailed evaluation metrics of hybrid models.

---

##### Supplementary Table S5.1: Evaluation metrics of ML models developed using compositional features combined with MERCI.

| Model | Sn | Sp | Acc | MCC | AUC |
|---|---|---|---|---|---|
| **RF** | 0.816 | 0.826 | 0.821 | 0.641 | 0.900 |
| **DT** | 0.710 | 0.768 | 0.741 | 0.478 | 0.779 |
| **ET** | 0.760 | 0.821 | 0.793 | 0.583 | 0.881 |
| **MLP** | 0.782 | 0.807 | 0.795 | 0.589 | 0.875 |
| **SVC** | 0.832 | 0.739 | 0.782 | 0.571 | 0.855 |
| **XGBC** | 0.799 | 0.836 | 0.819 | 0.635 | 0.890 |
| **LR** | 0.743 | 0.821 | 0.785 | 0.567 | 0.864 |

##### Supplementary Table S5.2: Evaluation metrics of ML models developed using PLM embedding combined with MERCI.

| Embedding source | ML model | Sn | Sp | Acc | MCC | AUC |
|---|---|---|---|---|---|---|
| **BioBERT** | **MLPC** | 0.715 | 0.836 | 0.780 | 0.557 | 0.837 |
| **ESM2t33** | **ET** | 0.816 | 0.816 | 0.816 | 0.631 | 0.899 |
| **ESM2t36** | **XGBC** | 0.788 | 0.860 | 0.826 | 0.651 | 0.892 |
| **ProtBERT** | **MLPC** | 0.765 | 0.874 | 0.824 | 0.646 | 0.892 |
| **ESM2t6** | **ET** | 0.777 | 0.865 | 0.824 | 0.646 | 0.892 |
| **ProtBERT+BiLSTM** | **ET** | 0.754 | 0.858 | 0.597 | 0.879 | 0.848 |

#### Supplementary Table S5.3: Evaluation metrics of PLM combined with MERCI.

| Metric | Sn | Sp | Acc | MCC | AUC |
|---|---|---|---|---|---|
| **BioBERT** | 0.676 | 0.792 | 0.738 | 0.472 | 0.767 |
| **ProtBert** | 0.659 | 0.903 | 0.790 | 0.585 | 0.871 |
| **ESM2-t6** | 0.883 | 0.792 | 0.834 | 0.674 | 0.909 |
| **ESM2-t33** | 0.687 | 0.942 | 0.824 | 0.658 | 0.890 |

### Supplementary Table S6: The detailed evaluation metrics of regression models used to predict $HC_{50}$ value of peptides.

---

#### Supplementary Table S6.1: Evaluation metrics of ML models using compositional features.

**Feature: Amino Acid Composition ( 20 descriptors)**

| Model | Dataset | R | $R^2$ | MAE | MSE |
|---|---|---|---|---|---|
| XGBRegressor | **Cross-validation** | 0.607 | 0.324 | 0.894 | 1.555 |
| XGBRegressor | **Independent** | 0.611 | 0.320 | 0.901 | 1.461 |
| Random Forest Regression | **Cross-validation** | 0.648 | 0.412 | 0.839 | 1.351 |
| Random Forest Regression | **Independent** | 0.681 | 0.456 | 0.826 | 1.166 |
| Gradient Boosting Regression | **Cross-validation** | 0.596 | 0.348 | 0.922 | 1.499 |
| Gradient Boosting Regression | **Independent** | 0.648 | 0.410 | 0.885 | 1.267 |
| ExtraTreesRegressor | **Cross-validation** | 0.652 | 0.415 | 0.811 | 1.344 |
| ExtraTreesRegressor | **Independent** | 0.687 | 0.469 | 0.788 | 1.141 |
| Decision Tree Regression | **Cross-validation** | 0.418 | -0.200 | 1.143 | 2.734 |
| Decision Tree Regression | **Independent** | 0.476 | -0.153 | 1.096 | 2.479 |
| AdaBoost Regression | **Cross-validation** | 0.446 | 0.165 | 1.110 | 1.913 |
| AdaBoost Regression | **Independent** | 0.427 | 0.144 | 1.132 | 1.839 |
| Support Vector Regression | **Cross-validation** | 0.574 | 0.308 | 0.907 | 1.590 |
| Support Vector Regression | **Independent** | 0.589 | 0.336 | 0.893 | 1.426 |
| K-Nearest Neighbors Regression | **Cross-validation** | 0.569 | 0.303 | 0.923 | 1.602 |
| K-Nearest Neighbors Regression | **Independent** | 0.613 | 0.368 | 0.856 | 1.358 |
| Linear Regression | **Cross-validation** | 0.444 | 0.189 | 1.057 | 1.861 |
| Linear Regression | **Independent** | 0.449 | 0.200 | 1.046 | 1.719 |

**Feature: Di-peptide composition ( 20 descriptors)**

| Model | Dataset | R | $R^2$ | MAE | MSE |
|---|---|---|---|---|---|
| XGBRegressor | **Cross-validation** | 0.609 | 0.320 | 0.880 | 1.561 |
| XGBRegressor | **Independent** | 0.696 | 0.468 | 0.785 | 1.144 |
| Random Forest Regression | **Cross-validation** | 0.629 | 0.387 | 0.850 | 1.411 |
| Random Forest Regression | **Independent** | 0.702 | 0.469 | 0.837 | 1.139 |
| Gradient Boosting Regression | **Cross-validation** | 0.618 | 0.376 | 0.895 | 1.432 |
| Gradient Boosting Regression | **Independent** | 0.639 | 0.406 | 0.862 | 1.275 |
| ExtraTreesRegressor | **Cross-validation** | 0.612 | 0.347 | 0.852 | 1.500 |
| ExtraTreesRegressor | **Independent** | 0.739 | 0.546 | 0.718 | 0.976 |
| Decision Tree Regression | **Cross-validation** | 0.451 | -0.139 | 1.119 | 2.611 |
| Decision Tree Regression | **Independent** | 0.544 | 0.036 | 1.017 | 2.071 |
| AdaBoost Regression | **Cross-validation** | 0.516 | 0.233 | 1.049 | 1.759 |
| AdaBoost Regression | **Independent** | 0.495 | 0.207 | 1.062 | 1.704 |
| Support Vector Regression | **Cross-validation** | 0.597 | 0.342 | 0.899 | 1.513 |
| Support Vector Regression | **Independent** | 0.610 | 0.366 | 0.881 | 1.362 |
| K-Nearest Neighbors Regression | **Cross-validation** | 0.604 | 0.347 | 0.893 | 1.500 |
| K-Nearest Neighbors Regression | **Independent** | 0.662 | 0.434 | 0.796 | 1.217 |
| Linear Regression | **Cross-validation** | 0.523 | 0.238 | 1.005 | 1.750 |
| Linear Regression | **Independent** | 0.527 | 0.244 | 0.977 | 1.625 |

**Feature: Amino acid composition and di-peptide composition ( 639 descriptors)**

| Model | Dataset | R | $R^2$ | MAE | MSE |
|---|---|---|---|---|---|
| XGBRegressor | **Cross-validation** | 0.655 | 0.405 | 0.854 | 1.369 |
| XGBRegressor | **Independent** | 0.686 | 0.461 | 0.801 | 1.159 |
| Random Forest Regression | **Cross-validation** | 0.679 | 0.453 | 0.819 | 1.259 |
| Random Forest Regression | **Independent** | 0.719 | 0.515 | 0.772 | 1.043 |
| Gradient Boosting Regression | **Cross-validation** | 0.648 | 0.398 | 0.905 | 1.382 |
| Gradient Boosting Regression | **Independent** | 0.616 | 0.370 | 0.923 | 1.354 |
| ExtraTreesRegressor | **Cross-validation** | 0.665 | 0.434 | 0.811 | 1.300 |
| ExtraTreesRegressor | **Independent** | 0.712 | 0.505 | 0.762 | 1.064 |
| Decision Tree Regression | **Cross-validation** | 0.478 | -0.067 | 1.087 | 2.452 |
| Decision Tree Regression | **Independent** | 0.508 | -0.060 | 1.005 | 2.278 |
| AdaBoost Regression | **Cross-validation** | 0.494 | 0.166 | 1.119 | 1.910 |
| AdaBoost Regression | **Independent** | 0.491 | 0.157 | 1.133 | 1.811 |
| Support Vector Regression | **Cross-validation** | 0.585 | 0.321 | 0.879 | 1.561 |
| Support Vector Regression | **Independent** | 0.613 | 0.358 | 0.859 | 1.378 |
| K-Nearest Neighbors Regression | **Cross-validation** | 0.557 | 0.273 | 0.934 | 1.672 |
| K-Nearest Neighbors Regression | **Independent** | 0.629 | 0.387 | 0.847 | 1.316 |
| Linear Regression | **Cross-validation** | 0.355 | -0.606 | 1.250 | 3.725 |
| Linear Regression | **Independent** | 0.229 | -3.360 | 1.515 | 9.370 |

**Feature: AAC+DPC+CeTD+PCP ( 20 descriptors)**

| Model | Dataset | R | $R^2$ | MAE | MSE |
|---|---|---|---|---|---|
| XGBRegressor | **Cross-validation** | 0.683 | 0.434 | 0.879 | 1.526 |
| XGBRegressor | **Independent** | 0.690 | 0.487 | 0.816 | 1.317 |

| | | | | | |
|---|---|---|---|---|---|
| Random Forest Regression | **Cross-validation** | 0.708 | 0.492 | 0.839 | 1.151 |
| Random Forest Regression | **Independent** | 0.729 | 0.532 | 0.730 | 1.004 |
| Gradient Boosting Regression | **Cross-validation** | 0.596 | 0.448 | 0.922 | 1.497 |
| Gradient Boosting Regression | **Independent** | 0.649 | 0.411 | 0.886 | 1.265 |
| ExtraTreesRegressor | **Cross-validation** | 0.645 | 0.406 | 0.815 | 1.364 |
| ExtraTreesRegressor | **Independent** | 0.683 | 0.462 | 0.802 | 1.157 |
| Decision Tree Regression | **Cross-validation** | 0.648 | 0.412 | 0.839 | 1.351 |
| Decision Tree Regression | **Independent** | 0.680 | 0.457 | 0.812 | 1.166 |
| AdaBoost Regression | **Cross-validation** | 0.469 | 0.180 | 1.105 | 1.880 |
| AdaBoost Regression | **Independent** | 0.462 | 0.168 | 1.105 | 1.788 |
| Support Vector Regression | **Cross-validation** | 0.574 | 0.308 | 0.907 | 1.590 |
| Support Vector Regression | **Independent** | 0.589 | 0.336 | 0.893 | 1.426 |
| K-Nearest Neighbors Regression | **Cross-validation** | 0.569 | 0.303 | 0.923 | 1.602 |
| K-Nearest Neighbors Regression | **Independent** | 0.613 | 0.368 | 0.856 | 1.358 |
| Linear Regression | **Cross-validation** | 0.444 | 0.189 | 1.057 | 1.861 |
| Linear Regression | **Independent** | 0.449 | 0.200 | 1.046 | 1.719 |

#### Feature: CeTD ( 20 descriptors)

| Model | Dataset | R | $R^2$ | MAE | MSE |
|---|---|---|---|---|---|
| XGBRegressor | **Cross-validation** | 0.626 | 0.347 | 0.858 | 1.501 |
| XGBRegressor | **Independent** | 0.691 | 0.455 | 0.775 | 1.171 |
| Random Forest Regression | **Cross-validation** | 0.629 | 0.387 | 0.850 | 1.411 |
| Random Forest Regression | **Independent** | 0.702 | 0.492 | 0.770 | 1.093 |
| Gradient Boosting Regression | **Cross-validation** | 0.617 | 0.375 | 0.895 | 1.434 |
| Gradient Boosting Regression | **Independent** | 0.638 | 0.405 | 0.862 | 1.278 |
| ExtraTreesRegressor | **Cross-validation** | 0.614 | 0.350 | 0.850 | 1.493 |
| ExtraTreesRegressor | **Independent** | 0.738 | 0.543 | 0.727 | 0.981 |
| Decision Tree Regression | **Cross-validation** | 0.448 | -0.155 | 1.119 | 2.652 |
| Decision Tree Regression | **Independent** | 0.531 | 0.003 | 1.027 | 2.141 |
| AdaBoost Regression | **Cross-validation** | 0.529 | 0.242 | 1.044 | 1.739 |
| AdaBoost Regression | **Independent** | 0.557 | 0.282 | 0.997 | 1.542 |
| Support Vector Regression | **Cross-validation** | 0.597 | 0.342 | 0.899 | 1.513 |
| Support Vector Regression | **Independent** | 0.610 | 0.366 | 0.881 | 1.362 |
| K-Nearest Neighbors Regression | **Cross-validation** | 0.604 | 0.347 | 0.893 | 1.500 |
| K-Nearest Neighbors Regression | **Independent** | 0.662 | 0.434 | 0.796 | 1.217 |
| Linear Regression | **Cross-validation** | 0.523 | 0.238 | 1.005 | 1.750 |
| Linear Regression | **Independent** | 0.527 | 0.244 | 0.977 | 1.625 |

#### Feature: AAC+DPC+CeTD ( 609 descriptors)

| Model | Dataset | R | $R^2$ | MAE | MSE |
|---|---|---|---|---|---|
| XGBRegressor | **Cross-validation** | 0.640 | 0.425 | 0.808 | 1.072 |
| XGBRegressor | **Independent** | 0.718 | 0.511 | 0.697 | 0.989 |
| Random Forest Regression | **Cross-validation** | 0.703 | 0.484 | 0.788 | 1.185 |
| Random Forest Regression | **Independent** | 0.735 | 0.531 | 0.752 | 1.007 |
| Gradient Boosting Regression | **Cross-validation** | 0.603 | 0.476 | 0.852 | 1.255 |
| Gradient Boosting Regression | **Independent** | 0.607 | 0.488 | 0.825 | 1.148 |
| ExtraTreesRegressor | **Cross-validation** | 0.611 | 0.488 | 0.785 | 1.223 |

| | | | | | |
|---|---|---|---|---|---|
| ExtraTreesRegressor | **Independent** | 0.668 | 0.581 | 0.72 | 0.947 |
| Decision Tree Regression | **Cross-validation** | 0.528 | 0.316 | 1.075 | 2.307 |
| Decision Tree Regression | **Independent** | 0.517 | 0.325 | 1.063 | 2.25 |
| AdaBoost Regression | **Cross-validation** | 0.597 | 0.327 | 1.01 | 1.593 |
| AdaBoost Regression | **Independent** | 0.555 | 0.286 | 1.025 | 1.582 |
| Support Vector Regression | **Cross-validation** | 0.177 | 0.004 | 1.135 | 2.334 |
| Support Vector Regression | **Independent** | 0.173 | 0.019 | 1.16 | 2.156 |
| K-Nearest Neighbors Regression | **Cross-validation** | 0.391 | 0.108 | 1.137 | 2.089 |
| K-Nearest Neighbors Regression | **Independent** | 0.343 | 0.056 | 1.112 | 2.075 |
| Linear Regression | **Cross-validation** | 0.022 | 0.137 | 1.184 | 2.305 |
| Linear Regression | **Independent** | 0.05 | 0.209 | 1.344 | 2.689 |

#### Feature: ALLCOMP ( 1192 descriptors)

| Model | Dataset | R | $R^2$ | MAE | MSE |
|---|---|---|---|---|---|
| XGBRegressor | **Cross-validation** | 0.680 | 0.445 | 0.808 | 1.272 |
| XGBRegressor | **Independent** | 0.738 | 0.541 | 0.737 | 0.987 |
| Random Forest Regression | **Cross-validation** | 0.705 | 0.486 | 0.791 | 1.178 |
| Random Forest Regression | **Independent** | 0.730 | 0.531 | 0.741 | 1.008 |
| Gradient Boosting Regression | **Cross-validation** | 0.688 | 0.461 | 0.837 | 1.240 |
| Gradient Boosting Regression | **Independent** | 0.692 | 0.473 | 0.810 | 1.133 |
| ExtraTreesRegressor | **Cross-validation** | 0.696 | 0.473 | 0.770 | 1.208 |
| ExtraTreesRegressor | **Independent** | 0.753 | 0.566 | 0.705 | 0.932 |
| Decision Tree Regression | **Cross-validation** | 0.513 | 0.001 | 1.060 | 2.292 |
| Decision Tree Regression | **Independent** | 0.502 | -0.040 | 1.048 | 2.235 |
| AdaBoost Regression | **Cross-validation** | 0.582 | 0.312 | 0.995 | 1.578 |
| AdaBoost Regression | **Independent** | 0.540 | 0.271 | 1.010 | 1.567 |
| Support Vector Regression | **Cross-validation** | 0.162 | -0.011 | 1.120 | 2.319 |
| Support Vector Regression | **Independent** | 0.158 | 0.004 | 1.145 | 2.141 |
| K-Nearest Neighbors Regression | **Cross-validation** | 0.376 | 0.093 | 1.122 | 2.074 |
| K-Nearest Neighbors Regression | **Independent** | 0.328 | 0.041 | 1.097 | 2.060 |
| Linear Regression | **Cross-validation** | 0.007 | 0.122 | 1.169 | 2.290 |
| Linear Regression | **Independent** | 0.035 | 0.194 | 1.329 | 2.674 |

#### Feature: ALLCOMP-ex SPC ( 1167 descriptors)

| Model | Dataset | R | $R^2$ | MAE | MSE |
|---|---|---|---|---|---|
| XGBRegressor | **Cross-validation** | 0.693 | 0.463 | 0.792 | 1.229 |
| XGBRegressor | **Independent** | 0.707 | 0.481 | 0.770 | 1.115 |
| Random Forest Regression | **Cross-validation** | 0.714 | 0.517 | 0.783 | 1.103 |
| Random Forest Regression | **Independent** | 0.739 | 0.543 | 0.735 | 0.981 |
| Gradient Boosting Regression | **Cross-validation** | 0.687 | 0.460 | 0.844 | 1.239 |
| Gradient Boosting Regression | **Independent** | 0.700 | 0.485 | 0.806 | 1.107 |
| ExtraTreesRegressor | **Cross-validation** | 0.696 | 0.474 | 0.766 | 1.206 |
| ExtraTreesRegressor | **Independent** | 0.741 | 0.549 | 0.715 | 0.970 |
| Decision Tree Regression | **Cross-validation** | 0.509 | 0.006 | 1.047 | 2.285 |

| | | | | | |
|---|---|---|---|---|---|
| Decision Tree Regression | **Independent** | 0.534 | 0.048 | 1.023 | 2.046 |
| AdaBoost Regression | **Cross-validation** | 0.583 | 0.309 | 0.996 | 1.584 |
| AdaBoost Regression | **Independent** | 0.548 | 0.267 | 1.015 | 1.575 |
| Support Vector Regression | **Cross-validation** | 0.628 | 0.375 | 0.857 | 1.439 |
| Support Vector Regression | **Independent** | 0.682 | 0.443 | 0.815 | 1.197 |
| K-Nearest Neighbors Regression | **Cross-validation** | 0.575 | 0.294 | 0.930 | 1.619 |
| K-Nearest Neighbors Regression | **Independent** | 0.623 | 0.376 | 0.829 | 1.340 |
| Linear Regression | **Cross-validation** | 0.693 | 0.463 | 0.792 | 1.229 |
| Linear Regression | **Independent** | 0.707 | 0.481 | 0.770 | 1.115 |

#### Supplementary Table S6.2: Evaluation metrics of ML models using Protein language models embeddings

##### Feature: ProtBERT embeddings

| Model | Dataset | R | $R^2$ | MAE | MSE |
|---|---|---|---|---|---|
| XGBRegressor | **Cross-validation** | 0.507 | 0.176 | 1.145 | 1.345 |
| XGBRegressor | **Independent** | 0.379 | 0.019 | 1.096 | 1.451 |
| Random Forest Regression | **Cross-validation** | 0.545 | 0.265 | 1.019 | 1.308 |
| Random Forest Regression | **Independent** | 0.410 | 0.115 | 1.076 | 1.407 |
| Gradient Boosting Regression | **Cross-validation** | 0.574 | 0.318 | 1.405 | 1.286 |
| Gradient Boosting Regression | **Independent** | 0.445 | 0.181 | 1.252 | 1.377 |
| ExtraTreesRegressor | **Cross-validation** | 0.479 | 0.097 | 1.462 | 1.378 |
| ExtraTreesRegressor | **Independent** | 0.377 | 0.015 | 1.509 | 1.453 |
| Decision Tree Regression | **Cross-validation** | 0.382 | -0.284 | 1.549 | 1.836 |
| Decision Tree Regression | **Independent** | 0.288 | -0.375 | 1.603 | 1.632 |
| AdaBoost Regression | **Cross-validation** | 0.587 | 0.320 | 1.410 | 1.286 |
| AdaBoost Regression | **Independent** | 0.460 | 0.190 | 1.459 | 1.773 |
| Support Vector Regression | **Cross-validation** | 0.607 | 0.358 | 1.387 | 1.269 |
| Support Vector Regression | **Independent** | 0.488 | 0.235 | 1.441 | 1.352 |
| K-Nearest Neighbors Regression | **Cross-validation** | 0.532 | 0.244 | 1.424 | 1.317 |
| K-Nearest Neighbors Regression | **Independent** | 0.432 | 0.141 | 1.472 | 1.395 |
| Linear Regression | **Cross-validation** | 0.127 | -5.789 | 1.259 | 2.833 |
| Linear Regression | **Independent** | 0.252 | -1.299 | 1.790 | 2.058 |

##### Feature: ESM-2t6 embeddings

| Model | Dataset | R | $R^2$ | MAE | MSE |
|---|---|---|---|---|---|
| XGBRegressor | **Cross-validation** | 0.622 | 0.374 | 0.885 | 1.439 |
| XGBRegressor | **Independent** | 0.644 | 0.404 | 0.862 | 1.280 |
| Random Forest Regression | **Cross-validation** | 0.673 | 0.427 | 0.854 | 1.318 |
| Random Forest Regression | **Independent** | 0.649 | 0.412 | 0.860 | 1.263 |
| Gradient Boosting Regression | **Cross-validation** | 0.653 | 0.418 | 0.875 | 1.339 |

| | | | | | |
|---|---|---|---|---|---|
| Gradient Boosting Regression | **Independent** | 0.629 | 0.393 | 0.884 | 1.305 |
| ExtraTreesRegressor | **Cross-validation** | 0.698 | 0.471 | 0.808 | 1.218 |
| ExtraTreesRegressor | **Independent** | 0.677 | 0.452 | 0.820 | 1.177 |
| Decision Tree Regression | **Cross-validation** | 0.413 | -0.230 | 1.179 | 2.823 |
| Decision Tree Regression | **Independent** | 0.440 | -0.219 | 1.137 | 2.619 |
| AdaBoost Regression | **Cross-validation** | 0.560 | 0.295 | 0.999 | 1.622 |
| AdaBoost Regression | **Independent** | 0.546 | 0.260 | 1.031 | 1.590 |
| Support Vector Regression | **Cross-validation** | 0.450 | 0.151 | 1.039 | 1.952 |
| Support Vector Regression | **Independent** | 0.476 | 0.185 | 1.044 | 1.752 |
| K-Nearest Neighbors Regression | **Cross-validation** | 0.589 | 0.332 | 0.905 | 1.538 |
| K-Nearest Neighbors Regression | **Independent** | 0.599 | 0.336 | 0.878 | 1.427 |
| Linear Regression | **Cross-validation** | 0.564 | 0.219 | 0.989 | 1.794 |
| Linear Regression | **Independent** | 0.607 | 0.316 | 0.927 | 1.470 |

#### Feature: ESM-2t6 embeddings

| Model | Dataset | R | $R^2$ | MAE | MSE |
|---|---|---|---|---|---|
| XGBRegressor | **Cross-validation** | 0.658 | 0.425 | 0.853 | 1.323 |
| XGBRegressor | **Independent** | 0.667 | 0.438 | 0.821 | 1.207 |
| Random Forest Regression | **Cross-validation** | 0.675 | 0.437 | 0.845 | 1.297 |
| Random Forest Regression | **Independent** | 0.702 | 0.478 | 0.814 | 1.122 |
| Gradient Boosting Regression | **Cross-validation** | 0.663 | 0.431 | 0.858 | 1.309 |
| Gradient Boosting Regression | **Independent** | 0.661 | 0.435 | 0.850 | 1.215 |
| ExtraTreesRegressor | **Cross-validation** | 0.695 | 0.467 | 0.810 | 1.226 |
| ExtraTreesRegressor | **Independent** | 0.711 | 0.495 | 0.786 | 1.084 |
| Decision Tree Regression | **Cross-validation** | 0.418 | -0.228 | 1.188 | 2.810 |
| Decision Tree Regression | **Independent** | 0.353 | -0.344 | 1.254 | 2.887 |
| AdaBoost Regression | **Cross-validation** | 0.611 | 0.354 | 0.955 | 1.484 |
| AdaBoost Regression | **Independent** | 0.566 | 0.316 | 0.970 | 1.471 |
| Support Vector Regression | **Cross-validation** | 0.514 | 0.227 | 0.988 | 1.779 |
| Support Vector Regression | **Independent** | 0.569 | 0.283 | 0.956 | 1.540 |
| K-Nearest Neighbors Regression | **Cross-validation** | 0.602 | 0.345 | 0.885 | 1.506 |
| K-Nearest Neighbors Regression | **Independent** | 0.610 | 0.350 | 0.863 | 1.397 |
| Linear Regression | **Cross-validation** | 0.155 | -24.258 | 5.387 | 58.390 |
| Linear Regression | **Independent** | 0.333 | -3.464 | 2.192 | 9.592 |

#### Feature: ESM-2t33 embeddings

| Model | Dataset | R | $R^2$ | MAE | MSE |
|---|---|---|---|---|---|
| XGBRegressor | **Cross-validation** | 0.658 | 0.425 | 0.853 | 1.323 |
| XGBRegressor | **Independent** | 0.667 | 0.438 | 0.821 | 1.207 |
| Random Forest Regression | **Cross-validation** | 0.675 | 0.437 | 0.845 | 1.297 |
| Random Forest Regression | **Independent** | 0.702 | 0.478 | 0.814 | 1.122 |
| Gradient Boosting Regression | **Cross-validation** | 0.663 | 0.431 | 0.858 | 1.309 |
| Gradient Boosting Regression | **Independent** | 0.661 | 0.435 | 0.850 | 1.215 |
| ExtraTreesRegressor | **Cross-validation** | 0.695 | 0.467 | 0.810 | 1.226 |

| | | | | | |
|---|---|---|---|---|---|
| ExtraTreesRegressor | **Independent** | 0.711 | 0.495 | 0.786 | 1.084 |
| Decision Tree Regression | **Cross-validation** | 0.418 | -0.228 | 1.188 | 2.810 |
| Decision Tree Regression | **Independent** | 0.353 | -0.344 | 1.254 | 2.887 |
| AdaBoost Regression | **Cross-validation** | 0.611 | 0.354 | 0.955 | 1.484 |
| AdaBoost Regression | **Independent** | 0.566 | 0.316 | 0.970 | 1.471 |
| Support Vector Regression | **Cross-validation** | 0.514 | 0.227 | 0.988 | 1.779 |
| Support Vector Regression | **Independent** | 0.569 | 0.283 | 0.956 | 1.540 |
| K-Nearest Neighbors Regression | **Cross-validation** | 0.602 | 0.345 | 0.885 | 1.506 |
| K-Nearest Neighbors Regression | **Independent** | 0.610 | 0.350 | 0.863 | 1.397 |
| Linear Regression | **Cross-validation** | 0.155 | -24.258 | 5.387 | 58.390 |
| Linear Regression | **Independent** | 0.333 | -3.464 | 2.192 | 9.592 |

#### Feature: ESM-2t36 embeddings

| Model | Dataset | R | $R^2$ | MAE | MSE |
|---|---|---|---|---|---|
| XGBRegressor | **Cross-validation** | 0.634 | 0.391 | 0.867 | 1.398 |
| XGBRegressor | **Independent** | 0.687 | 0.472 | 0.794 | 1.135 |
| Random Forest Regression | **Cross-validation** | 0.665 | 0.419 | 0.864 | 1.334 |
| Random Forest Regression | **Independent** | 0.688 | 0.457 | 0.831 | 1.167 |
| Gradient Boosting Regression | **Cross-validation** | 0.647 | 0.411 | 0.868 | 1.354 |
| Gradient Boosting Regression | **Independent** | 0.683 | 0.459 | 0.839 | 1.163 |
| ExtraTreesRegressor | **Cross-validation** | 0.679 | 0.445 | 0.828 | 1.273 |
| ExtraTreesRegressor | **Independent** | 0.706 | 0.486 | 0.808 | 1.105 |
| Decision Tree Regression | **Cross-validation** | 0.401 | -0.206 | 1.202 | 2.762 |
| Decision Tree Regression | **Independent** | 0.386 | -0.228 | 1.151 | 2.639 |
| AdaBoost Regression | **Cross-validation** | 0.589 | 0.324 | 0.978 | 1.550 |
| AdaBoost Regression | **Independent** | 0.571 | 0.312 | 0.989 | 1.479 |
| Support Vector Regression | **Cross-validation** | 0.478 | 0.178 | 1.020 | 1.890 |
| Support Vector Regression | **Independent** | 0.514 | 0.215 | 0.999 | 1.686 |
| K-Nearest Neighbors Regression | **Cross-validation** | 0.585 | 0.321 | 0.900 | 1.557 |
| K-Nearest Neighbors Regression | **Independent** | 0.614 | 0.366 | 0.845 | 1.363 |
| Linear Regression | **Cross-validation** | 0.450 | -0.610 | 1.406 | 3.666 |
| Linear Regression | **Independent** | 0.423 | -1.194 | 1.587 | 4.715 |

#### Feature: BioBERT embeddings

| Model | Dataset | R | $R^2$ | MAE | MSE |
|---|---|---|---|---|---|
| XGBRegressor | **Cross-validation** | 0.499 | 0.231 | 1.000 | 1.765 |
| XGBRegressor | **Independent** | 0.556 | 0.303 | 0.963 | 1.497 |
| Random Forest Regression | **Cross-validation** | 0.562 | 0.286 | 0.973 | 1.640 |
| Random Forest Regression | **Independent** | 0.583 | 0.317 | 0.977 | 1.467 |
| Gradient Boosting Regression | **Cross-validation** | 0.563 | 0.309 | 0.949 | 1.585 |
| Gradient Boosting Regression | **Independent** | 0.593 | 0.346 | 0.954 | 1.406 |
| ExtraTreesRegressor | **Cross-validation** | 0.574 | 0.313 | 0.944 | 1.578 |
| ExtraTreesRegressor | **Independent** | 0.616 | 0.366 | 0.927 | 1.362 |
| Decision Tree Regression | **Cross-validation** | 0.301 | -0.403 | 1.333 | 3.207 |
| Decision Tree Regression | **Independent** | 0.234 | -0.609 | 1.359 | 3.458 |

| | | | | | |
|---|---|---|---|---|---|
| AdaBoost Regression | **Cross-validation** | 0.471 | 0.212 | 1.046 | 1.807 |
| AdaBoost Regression | **Independent** | 0.435 | 0.183 | 1.086 | 1.755 |
| Support Vector Regression | **Cross-validation** | 0.523 | 0.228 | 0.965 | 1.772 |
| Support Vector Regression | **Independent** | 0.554 | 0.267 | 0.967 | 1.575 |
| K-Nearest Neighbors Regression | **Cross-validation** | 0.460 | 0.190 | 1.027 | 1.860 |
| K-Nearest Neighbors Regression | **Independent** | 0.458 | 0.183 | 1.025 | 1.756 |
| Linear Regression | **Cross-validation** | 0.485 | 0.026 | 1.136 | 2.232 |
| Linear Regression | **Independent** | 0.505 | -0.033 | 1.137 | 2.220 |

##### Feature: ProtBERT+BiLSTM embeddings

| Model | Dataset | R | $R^2$ | MAE | MSE |
|---|---|---|---|---|---|
| XGBRegressor | **Cross-validation** | 0.475 | 0.241 | 0.640 | 1.113 |
| XGBRegressor | **Independent** | 0.486 | 0.264 | 0.621 | 0.984 |
| Random Forest Regression | **Cross-validation** | 0.602 | 0.471 | 0.739 | 1.048 |
| Random Forest Regression | **Independent** | 0.646 | 0.449 | 0.825 | 1.265 |
| Gradient Boosting Regression | **Cross-validation** | 0.495 | 0.268 | 0.650 | 1.056 |
| Gradient Boosting Regression | **Independent** | 0.525 | 0.315 | 0.590 | 0.874 |
| ExtraTreesRegressor | **Cross-validation** | 0.629 | 0.415 | 0.982 | 0.945 |
| ExtraTreesRegressor | **Independent** | 0.548 | 0.452 | 0.837 | 0.794 |
| Decision Tree Regression | **Cross-validation** | 0.238 | -0.371 | 0.979 | 2.507 |
| Decision Tree Regression | **Independent** | 0.255 | -0.260 | 0.860 | 2.108 |
| AdaBoost Regression | **Cross-validation** | 0.433 | 0.175 | 0.752 | 1.267 |
| AdaBoost Regression | **Independent** | 0.421 | 0.171 | 0.762 | 1.184 |
| Support Vector Regression | **Cross-validation** | -0.029 | -0.201 | 0.930 | 2.130 |
| Support Vector Regression | **Independent** | -0.032 | -0.186 | 0.955 | 1.950 |
| K-Nearest Neighbors Regression | **Cross-validation** | 0.178 | -0.104 | 0.935 | 1.899 |
| K-Nearest Neighbors Regression | **Independent** | 0.144 | -0.143 | 0.910 | 1.857 |
| Linear Regression | **Cross-validation** | 0.152 | -2.292 | 1.681 | 6.897 |
| Linear Regression | **Independent** | 0.156 | -3.821 | 1.892 | 9.761 |

#### List of Abbreviations

| **Abbreviation** | **Definition** |
|---|---|
| AAC | Amino Acid Composition (20 descriptors) |
| DPC | Di-peptide Composition (400 descriptors) |

| | |
|---|---|
| PCP | Physico-chemical properties (30 descriptors) |
| RRI | Residue Repeats Index (20 descriptors) |
| PRI | Property Repeats Index (25 descriptors) |
| SER & SEP | Shannon Entropy measures (46 descriptors |
| CeTD | Composition enhanced Transition and Distribution (189 descriptors) |
| PAAC | Pseudo Amino Acid Composition (21 descriptors) |
| APAAC | Amphiphilic Pseudo Amino Acid Composition (23 descriptors) |
| SOC | Sequence Order Coupling Number (2 descriptors) |
| Sp | Specificity |
| Sn | Sensitivity |
| Acc | Accuracy |
| MCC | Matthews correlation coefficient |
| AUC | Area under receiver operating characteristic |
| R | Pearson Correlation Coefficient |
| $R^2$ | Coefficient of Determination |
| MAE | Mean Absolute Error |
| MSE | Mean Squared Error |
